# Using deep long-read RNAseq in Alzheimer’s disease brain to assess medical relevance of RNA isoform diversity

**DOI:** 10.1101/2023.08.06.552162

**Authors:** Bernardo Aguzzoli Heberle, J. Anthony Brandon, Madeline L. Page, Kayla A. Nations, Ketsile I. Dikobe, Brendan J. White, Lacey A. Gordon, Grant A. Fox, Mark E. Wadsworth, Patricia H. Doyle, Brittney A. Williams, Edward J. Fox, Anantharaman Shantaraman, Mina Ryten, Sara Goodwin, Elena Ghiban, Robert Wappel, Senem Mavruk-Eskipehlivan, Justin B. Miller, Nicholas T. Seyfried, Peter T. Nelson, John D. Fryer, Mark T. W. Ebbert

## Abstract

Due to alternative splicing, human protein-coding genes average over eight RNA isoforms, resulting in nearly four distinct protein coding sequences per gene. Long-read RNAseq (IsoSeq) enables more accurate quantification of isoforms, shedding light on their specific roles. To assess the medical relevance of measuring RNA isoform expression, we sequenced 12 aged human frontal cortices (6 Alzheimer’s disease cases and 6 controls; 50% female) using one Oxford Nanopore PromethION flow cell per sample. Our study uncovered 53 new high-confidence RNA isoforms in medically relevant genes, including several where the new isoform was one of the most highly expressed for that gene. Specific examples include *WDR4* (61%; microcephaly), *MYL3* (44%; hypertrophic cardiomyopathy), and *MTHFS* (25%; major depression, schizophrenia, bipolar disorder). Other notable genes with new high-confidence isoforms include *CPLX2* (10%; schizophrenia, epilepsy) and *MAOB* (9%; targeted for Parkinson’s disease treatment). We identified 1,917 medically relevant genes expressing multiple isoforms in human frontal cortex, where 1,018 had multiple isoforms with different protein coding sequences, demonstrating the need to better understand how individual isoforms from a single gene body are involved in human health and disease, if at all. Exactly 98 of the 1,917 genes are implicated in brain-related diseases, including Alzheimer’s disease genes such as *APP* (Aβ precursor protein; five), *MAPT* (tau protein; four), and *BIN1* (eight). As proof of concept, we also found 99 differentially expressed RNA isoforms between Alzheimer’s cases and controls, despite the genes themselves not exhibiting differential expression. Our findings highlight the significant knowledge gaps in RNA isoform diversity and their medical relevance. Deep long-read RNA sequencing will be necessary going forward to fully comprehend the medical relevance of individual isoforms for a “single” gene.

## Main

Due to alternative splicing, human protein-coding genes average over eight RNA isoforms, resulting in nearly four distinct protein coding sequences^1,2^, yet because of practical limitations in standard short-read sequencing technologies, researchers have historically been forced to collapse all isoforms into a single gene expression measurement—a major oversimplification of the underlying biology. Given the large number of isoforms derived from a single gene body, we question whether these genes have a single function, as it is possible that many of the distinct RNA and resulting protein isoforms have distinct functions. In fact, Yang et al. demonstrated that many unique isoforms from a single gene body appear to have unique interactomes at the protein level^3^. Additionally, distinct functions for individual isoforms from a single gene body have already been demonstrated for a handful of genes^4–6^.

Detailed analysis of individual isoforms has been limited to highly studied genes because studying individual RNA isoforms via high-throughput methods has been challenging since short-read data relies on heuristics to assemble isoforms^7–9^, but heuristics cannot outperform directly measuring the trait in question. In principle, long reads have the potential to sequence entire isoforms, providing a direct measurement for each isoform, but long-read data is still far from perfect^10^ and also requires its own heuristics—specifically to estimate expression for each isoform^10,11^. Still, the significantly longer reads has clearly improved our ability to characterize and quantify RNA isoforms beyond what short reads enable^10^.

While determining a given RNA isoform’s downstream function will ultimately require experimental validation, long-read sequencing enables researchers to better understand their role in human health and disease by quantifying expression for each isoform across different tissues and cell types. Knowing which tissues and cell types express each isoform is an important first step in understanding their function. Notably, isoforms can also play entirely different, or even opposite roles within a given cell; a classic example includes two well-studied *BCL-X (BCL2L1)* transcripts with opposite functions, where *BCL-X_L_*is anti-apoptotic while *BCL-X_S_* is pro-apoptotic^6^. Changes in expression ratio between the *BCL-X* isoforms are implicated in cancer and are being studied as therapeutic targets^12^, demonstrating the importance of understanding individual RNA isoform function rather than treating them as a “single” gene (i.e., a single function).

Traditionally, RNAseq studies relied on short-read sequencing approaches that excel at quantifying gene-level expression, but cannot accurately assemble and quantify individual RNA isoforms^8,13^ (**Fig. 1a**), forcing researchers to collapse RNA isoforms into a single gene measurement. Long-read sequencing, however, can sequence entire RNA molecules, providing more accurate quantification of RNA isoforms, including *de novo* isoforms and genes (**Fig. 1a**). With this technology, researchers can start uncovering individual RNA isoform function and their relationship to human diseases for poorly understood genes and isoforms. For example, recent long-read RNAseq studies used targeted approaches to uncover aberrant splicing events in sporadic Alzheimer’s disease^14^, dystrophinopathies^15^, and cancers^16,17^.

**Fig. 1:**
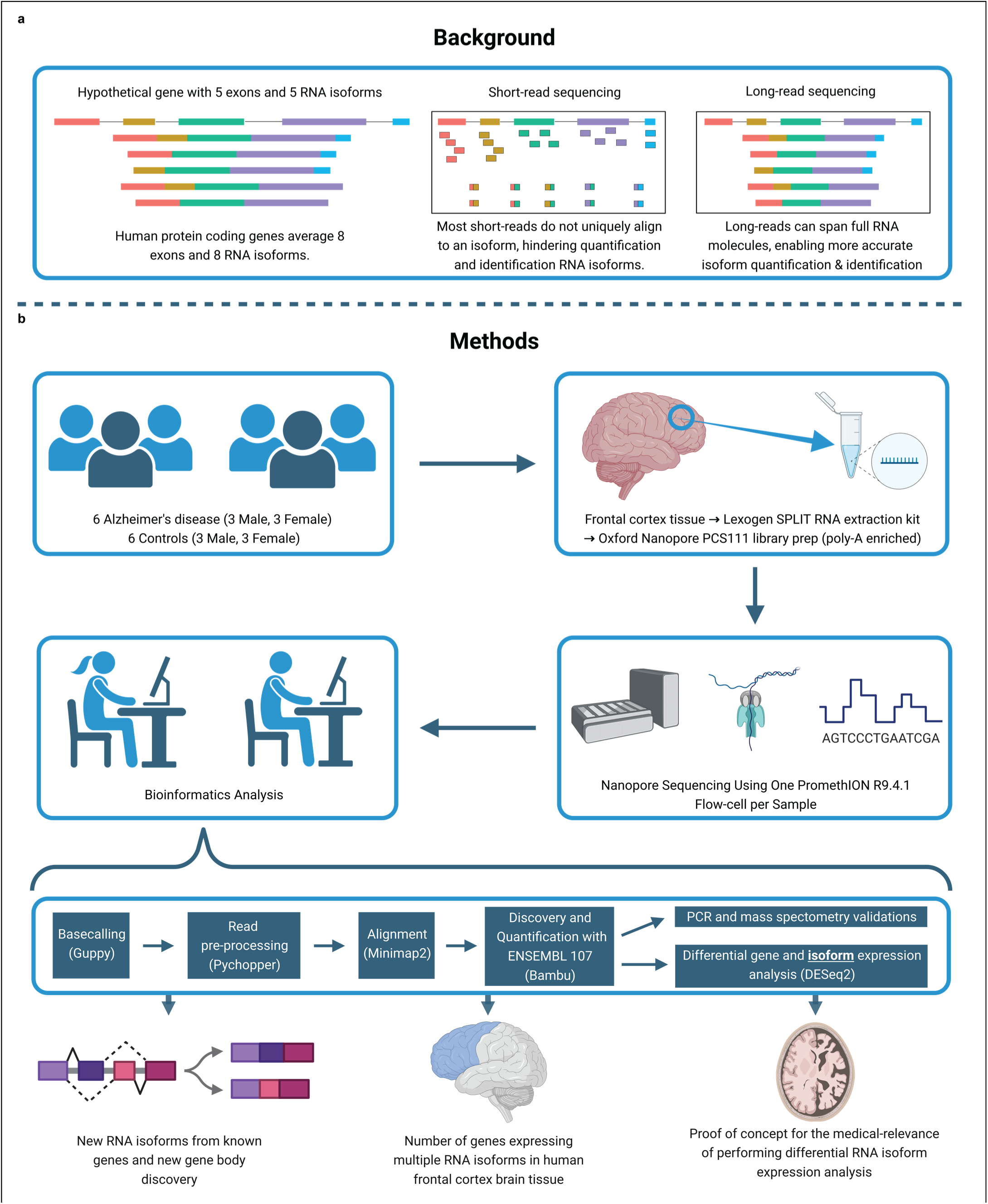
Study design and rationale. **a,** Background explaining the improvements long-read sequencing brings to the study of RNA isoforms. **b,** Details for experimental design, methods, and a summary of the topics explored in this article. Created with <u>BioRender.com</u>.

Two other studies demonstrated that long-read sequencing can discover new RNA isoforms across several human tissues, including the brain^18,19^; while both studies revealed important biology, including reporting *de novo* RNA isoforms, they had limited sequencing coverage (averaging <6 million aligned reads per sample). Read depth is essential to accurately quantify individual RNA isoforms, given that a total of more than 250,000 annotated RNA isoforms have been reported, as of July 2023^2^. Additionally, neither study focused on the medical relevance of using long read RNAseq. While long-read sequencing does not resolve all challenges related to isoform sequencing (e.g., those related to RNA degradation, etc.), our goal is to demonstrate the utility and importance of using long-read sequencing for both academic research and clinical diagnostics in the context of RNA isoforms (e.g., reporting newly discovered RNA isoforms in medically relevant genes, and variant interpretation in genes expressing multiple RNA isoforms).

In this study, we demonstrate that accurate RNA isoform quantification through deep (∼35.5 million aligned reads per sample) long-read sequencing can improve our understanding of individual RNA isoform function, and provide insights into how they may impact human health and disease. Specifically, in addition to discovering (1) *de novo* (i.e., unannotated) RNA isoforms in known medically relevant genes, we also discovered (2) *de novo* spliced mitochondria-encoded RNA isoforms, (3) entirely new gene bodies in nuclear DNA, and (4) demonstrate the complexity of RNA isoform diversity for medically relevant genes within a single tissue (human frontal cortex from Alzheimer’s disease cases and controls). Lastly, (5) we provide a proof of concept for differential RNA isoform expression analysis, demonstrating its potential to reveal disease-relevant transcriptomic signatures unavailable at the gene level (i.e., collapsing all isoforms into a single expression measurement). To do so, we sequenced human frontal cortex brain tissue using Oxford Nanopore Technologies deep long-read sequencing (**Fig. 1b**). Summary data from this study are readily explorable through a public web application to visualize individual RNA isoform expression in aged human frontal cortex tissue (https://ebbertlab.com/brain_rna_isoform_seq.html).

## Results

### Methodological and results overview

We sequenced 12 post-mortem aged brain samples (Brodmann area 9/46) from six Alzheimer’s disease cases and six cognitively unimpaired controls (50% female; **Fig. 1b**). All samples had postmortem intervals < 5hrs and RNA Integrity Score (RIN) ≥ 9.0; demographics, summary sequencing statistics, and read length distributions are shown in **Supplementary Table 1** and **Supplementary** Fig. 1-4. Total RNA was extracted using the Lexogen SPLIT RNA extraction kit, and cDNA was prepared using the Oxford Nanopore Technologies PCS111 library preparation kit. The PCS111 library preparation kit includes polyA enrichment. Each sample was sequenced using one PromethION flow cell. Sequencing yielded an average of 35.5 million aligned reads per sample after filtering out all reads that did not contain the primers on both ends, and those with a mapping quality < 10 (**Extended Data** Fig. 1a). By excluding all reads missing primers, we believe reads included in this study closely represent the RNA as it was at extraction. Also, even though our study involves polyA selection, we still refer to the isoforms as “RNA isoforms” rather than “mRNA isoforms” because mRNA is often treated as synonymous with protein-coding, but many long non-coding RNAs are polyadenylated and are not considered mRNAs, but mRNA-like^20^

We performed stringent RNA isoform quantification and discovery (including *de novo* gene bodies) using bambu^11^ (**Fig. 1b**)—a tool with emphasis on reducing false-positive RNA isoform discovery compared to other commonly used tools^11^. Bambu was highlighted as one of the top performers in a recent benchmark study^10^. However, as a tradeoff for higher precision, bambu is unable to discover new RNA isoforms that only differ from an annotated RNA isoform because of an alternative transcription start and/or end site (e.g., shortened 5’ UTR). When it comes to quantification, the increasing complexity of annotations can impact quantification due to non-unique reads being split between multiple transcripts. For example, if a read maps equally well to two RNA isoforms, each isoform will receive credit for 0.5 reads.

Isoform and gene body quantification and discovery were based on version 107 of Ensembl GRCh38 gene annotations (July 2022). For our 12 samples, bambu reported an average of 42.4% reads uniquely assigned to an RNA isoform and 17.5% reads spanning a full-length RNA isoform (**Extended Data** Fig. 1c). We only considered an isoform to be expressed above noise levels if its median counts per million (CPM) was greater than one (i.e., at least half of the samples had a CPM > 1); this threshold is dependent on overall depth, as lower depths will require a higher, more stringent CPM threshold. Using this threshold, we observed 28,989 expressed RNA isoforms from 18,041 gene bodies in our samples (**Extended Data** Fig. 2a,b**,c**). Based on official ENSEMBL isoform annotation biotypes, out of the mRNA isoforms expressed with median CPM > 1, 20,183 were classified protein coding, 2,303 were long non-coding RNAs, 3,213 were classified as having a retained intron, and the remaining 3,290 were scattered across other biotypes—including novel transcripts (**Extended Data** Fig. 3). Examples of medically relevant genes with highly expressed intron retention transcripts are: *PSEN1*, *MTOR*, *HACE1*, *TPM3*, *HNRNPU*.

We used publicly available mass spectrometry data from aged human dorsolateral frontal cortex tissue and human cell-lines to validate new RNA isoforms at the protein level. We included the cell-line data because, as of 2023, it is the largest human proteome with the highest sequence coverage ever reported across multiple digestion enzymes, giving us the best chance to validate our newly discovered RNA isoforms at the protein level. We carefully confirmed isoforms with uniquely matching peptide hits, resulting in a small number of successful validations. We also leveraged existing short-read RNAseq data from the Religious Orders Study Memory and Aging Project (ROSMAP)^21^ and long-read RNAseq data from Glinos et al.^18^ to validate our newly discovered RNA isoforms and gene bodies. For interest, we also include a brief analysis using the new Telomere-to-Telomere (T2T) CHM13 reference genome.

### Discovery of new RNA isoforms from known gene bodies, including medically relevant genes

Our first goal was to identify and quantify *de novo* RNA isoforms expressed in human frontal cortex. In total, bambu discovered 1,534 new transcripts from known nuclear gene bodies (i.e., annotated nuclear gene bodies). Of these 1,534 new RNA isoforms, 636 had a median CPM between 0 and 0.25 and 470 had a median CPM between 0.26 and 1. While we expect that many of these new RNA isoforms with a median CPM ≤ 1 are legitimate, we consider them low-confidence discoveries and exclude them throughout the remainder of our analyses, except where explicitly noted, to err on the side of caution.

After excluding all isoforms with a median CPM ≤ 1, 428 isoforms remained that we consider high confidence (**Fig. 2a,b**). Of these 428 isoforms, 303 were from protein-coding genes (**Fig. 2a**). We report significantly fewer new isoforms compared to Glinos et al.^18^ (∼70,000) and Leung et al.^19^ (∼12,000) because of (1) differences in the reference database, (2) discovery tool employed^10,22^, and (3) stringency in what constitutes a new isoform. Specifically, Glinos et al. used gene annotations from 2016 when determining new isoforms. This is likely because they were trying to maintain consistency with previous GTEx releases, but approximately 50,000 new isoforms have already been annotated since then^2^. We also used a recently released tool (bambu) for isoform and gene discovery, which has shown to dramatically reduce false positives^10,11^. Finally, we set a stricter threshold for what we consider high-confidence isoforms, using a median CPM > 1. Given the depth of our data, a CPM = 1 corresponds to an average of 24 observed copies (i.e., counts) per sample. Other tools and papers, however, report certain isoforms as new, even if the isoform is only observed twice, or if it is only slightly different from other isoforms (i.e., within sequencing error rate). Approximately 69.4% of our newly discovered isoforms are unique to our data, when compared to Ensembl v107, Glinos et al., and Leung et al. (**Supplementary Table 2,3**). This is likely because of the differences in sequencing depth, discovery tool employed (i.e., bambu^11^ vs. FLAIR^23^, vs Cupcake^24^), and differences in subject age.

**Fig. 2:**
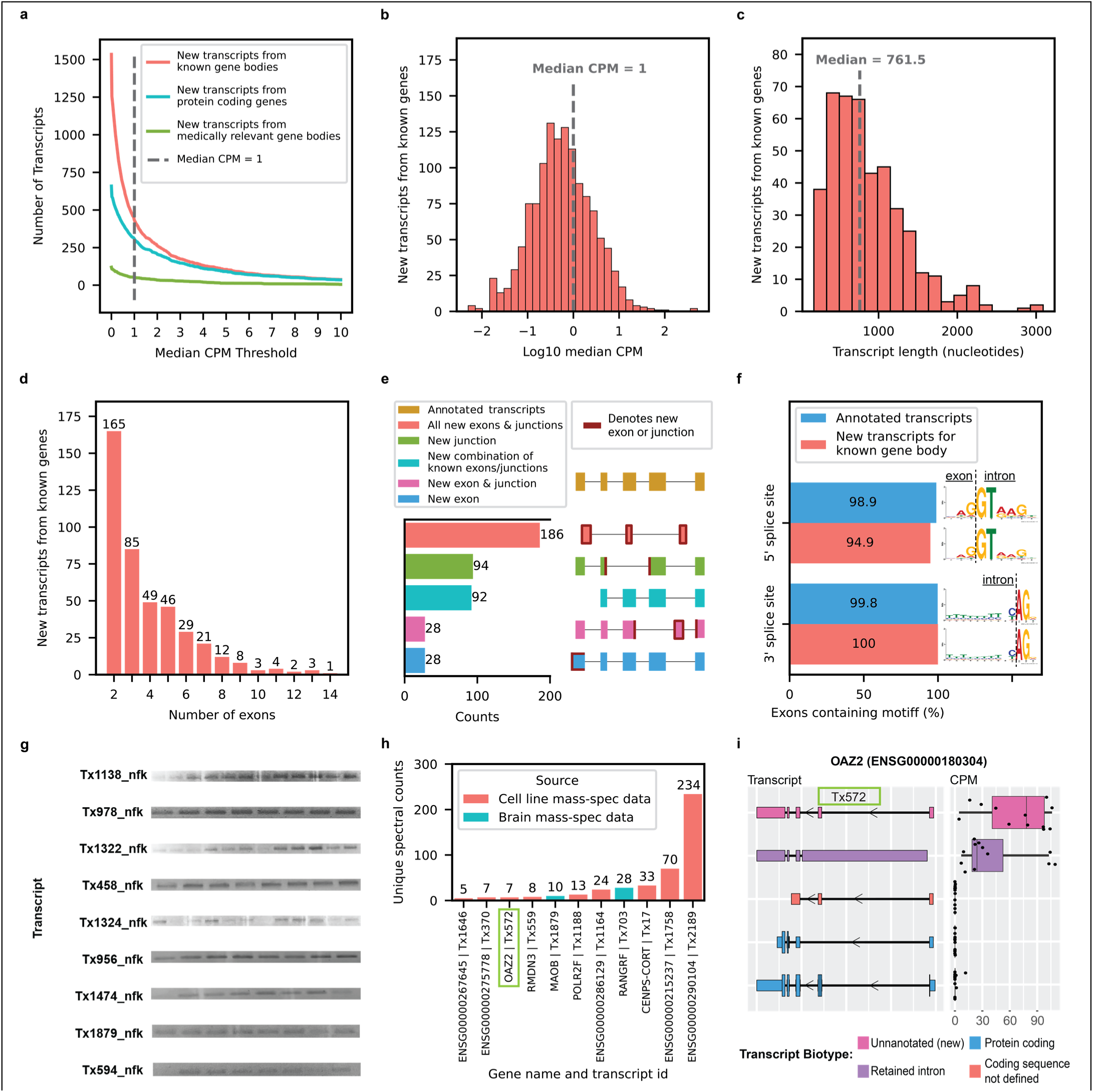
New high-confidence RNA isoforms and new spliced mitochondrial RNA isoforms expressed in human frontal cortex. Figures **a-f** refer to new transcripts from annotated gene bodies. **a,** Number of newly discovered transcripts across median CPM threshold. Cutoff shown as the dotted line set at median CPM = 1. **b,** Distribution of log10 median CPM values for newly discovered transcripts from annotated gene bodies, dotted line shows cutoff point of median CPM = 1. Figures **c-f** only include data from transcripts above this expression cutoff. **c,** Histogram showing distribution of transcripts length for new transcripts from annotated gene bodies. **d,** Bar plot showing the distribution of the number of exons for newly discovered transcripts from annotated gene bodies. **e,** Bar plot showing the kinds of events that gave rise to new transcripts from annotated gene bodies. For context, bambu considers modified exons (e.g., significantly longer or shorter) as new exons, including lengthened UTR regions. **f,** Bar plot showing the prevalence of canonical splice site motifs for annotated exons from transcripts with median CPM > 1 versus new exons from new transcripts in annotated gene bodies. **g,** Gel electrophoresis validation using PCR amplification for a subset of new RNA isoforms from known genes. This is an aggregate figure showing bands for several different gels. Individual gel figures are available in **Supplementary** Figures 1-26. **h**, Protein level validation using publicly available mass spectrometry proteomics data. Y-axis shows number of spectral counts from uniquely matching peptides (unique spectral counts); new transcripts from known gene bodies were considered validated at the protein level if they had more than 5 unique spectral counts. BambuTx1879, BambuTx1758, BambuTx2189 are unique to our study. **i**, RNA isoform structure and CPM expression for isoforms from *OAZ2* (cellular growth/proliferation). The new isoform Tx572 was most expressed and validated at the protein level (highlighted with the green box).

To determine what primarily drives the large difference in newly discovered RNA isoforms in our data vs. Glinos et al., we processed the Glinos et al. data through our pipeline using bambu with two different annotations: (1) ENSEMBL v88, as used in Glinos et al.; and (2) ENSEMBL v107, as used in this work. Using ENSEMBL v88, we identified a total of 4,594 new RNA isoforms whereas with ENSEMBL v107, we identified 3,796 new isoforms (difference of 798), demonstrating that, while the updated annotations do affect what is truly a new discovery, it is not the primary source of the difference.

Similarly, we performed a down sampling analysis to assess the importance of depth on our discoveries. Including all discoveries (i.e., including our low-confidence discoveries with median CPM ≤ 1), we only discovered 490 new isoforms from known genes with 20% of our aligned reads compared to 1,534 using 100% of our aligned reads (difference of 1,044; **Extended Data** Fig. 4a). Looking only at high-confidence discoveries in known genes, we discover 238 and 428 at 20% and 100% of reads, respectively (**Extended Data** Fig 4b), showing the importance of depth in our data. While both annotations and read depth were important to maximize discovery without reporting isoforms that have already been annotated since ENSEMBL v88, these do not explain the dramatic difference in reported discoveries between our work and Glinos et al. Thus, we conclude the primary driver is the discovery tool employed, which again, recent work suggests bambu is less likely to report false positives^10,11^. We also observed a 33.8% increase in transcript discovery overlap between our dataset and GTEx when using the same tools and annotation, supporting the idea that these are large drivers of differences between our findings (**Extended Data** Fig. 5).

New high-confidence isoforms had a median of 761.5 nucleotides in length, ranging from 179 to 3089 nucleotides (**Fig. 2c**); the number of exons in new high-confidence isoforms ranged between 2 and 14, with most isoforms falling on the lower end of the distribution (**Fig. 2d**). Our data were enriched for new RNA isoforms containing all new exons and exon-exon boundaries (i.e., exon junctions; **Fig. 2e**). For context, bambu considers modified exons as new (e.g., significantly longer, or shorter), excluding modified exons from transcripts that do not have any new exon junctions (e.g., shortened 3’ UTR). The 428 new high-confidence isoforms contained 737 new exon-intron boundaries where 94.9% (356/370) and 100% (367/367) of the 5’ and 3’ splice site matched canonical splice site motifs, respectively, supporting their biological feasibility (**Fig. 2f**). We attempted to validate 17 new high-confidence isoforms through PCR and gel electrophoresis and successfully validated nine of them (**Fig. 2g, Supplementary** Fig. 5-26**, Supplementary Table 4**). We then attempted to validate the eight RNA isoform that failed via standard PCR (no visible band on gel) using RT-qPCR—a more sensitive method compared to PCR and gel electrophoresis—and successfully validated six of them (**Supplementary Table 5**). MIQE guidelines by Bustin et al.^25^ suggests Ct < 40 as a cutoff for RT-qPCR validation, but we used a more stringent cutoff of Ct < 35 to be conservative. Out of the 15 transcripts that were successfully validated through PCR and gel electrophoresis or RT-qPCR, 11 are unique to this study. For interest, we compared relative abundance for known and new RNA isoforms between long-read sequencing and RT-qPCR for *MAOB, SLC26A1*, and *MT-RNR2*. The expression patterns were concordant between RT-qPCR and long-read sequencing for the isoforms of all three genes tested—*SLC26A1*, *MT-RNR2*, and *MAOB* (**Extended Data** Fig. 6**, Supplementary Table 6,7**).

We attempted to validate our new high-confidence transcripts from known genes using publicly available long-read RNAseq data from five GTEx^18^ brain (Brodmann area 9) samples and short-read RNAseq data from 251 ROSMAP^21^ brain samples (Brodmann area 9/46). We observed that 98.8% of the new high-confidence transcripts from known gene bodies had at least one uniquely mapped read in either GTEx or ROSMAP data and 69.6% had at least 100 uniquely mapped reads in either dataset (**Extended Data** Fig. 7, **Supplementary Table 8**). Importantly, while a single unique read within short-read data may seem like a soft threshold, only a small percentage of short reads are unique to a single isoform for genes expressing multiple isoforms.

For interest, we also include a brief analysis using the new Telomere-to-Telomere (T2T) CHM13 reference genome and validated 6 RNA isoforms across the 99 newly predicted protein-coding genes reported in Nurk et al.^26^ (**Extended Data** Fig. 8). Our validation threshold for the CHM13 analysis was at least 10 uniquely mapped reads in total across our 12 frontal cortex samples.

We also validated 11 of the new isoforms from known genes at the protein level using mass spectrometry data from the same brain region and human cell-lines (**Fig. 2h,i**). Three of the 11 we validated are unique to our study (BambuTx1879, BambuTx1758, BambuTx2189). We were particularly careful not to include new isoforms with a CPM ≤ 1 when performing protein validation via mass spectrometry because the sensitivity of mass spectrometry is exceptionally low for lowly expressed proteins, and including them would dramatically increase the search space and severely penalize statistical power.

### Medically relevant genes

Because identifying and quantifying all individual isoforms for a given gene is a critical first step to understanding all possible functions for a “single” gene, we wanted to assess how many new high-confidence isoforms originate from known medically relevant genes. For consistency, we used the list of medically relevant genes defined in Wagner et. al^27^, and we then further annotated with genes relevant to brain-related diseases^28–37^. Identifying and quantifying all isoforms is especially important for known medically relevant genes because, for example, when clinicians interpret the consequence of a genetic mutation, it is interpreted in the context of a single isoform of the parent gene body. That isoform may not even be expressed in the relevant tissue or cell type, however. Having a firm understanding of which specific tissues and cell types express each isoform for a given gene body is expressed in will be important to fully understanding its function or relevance to human health and disease.

Of the 428 new high-confidence isoforms, 53 originated from 49 medically relevant genes and we quantified the proportion of total expression for the gene that came from the new isoform(s) (**Fig. 3a**). The genes where the largest percentage of reads came from a newly discovered isoform include *SLC26A1* (86%; kidney stones^38^ and musculoskeletal health^39^), *CAMKMT* (61%; hypotonia-cystinuria syndrome, neonatal seizures, severe developmental delay, etc.^40^), *WDR4* (61%; microcephaly^41^ and Galloway-Mowat syndrome-6^42^), *MYL3* (44%; hypertrophic cardiomyopathy^43^), and *MTHFS* (25%; major depression, schizophrenia, and bipolar disorder^44^). Other notable genes with new high-confidence isoforms include *CPLX2* (10%; schizophrenia, epilepsy, and synaptic vesicle pathways^45^) and *MAOB* (9%; currently targeted for Parkinson’s disease treatment^46^; **Fig. 3c**); the new *MAOB* isoform includes a retained intron. We also found an unannotated RNA isoform for *TREM2* (16%; **Fig. 3b**), one of the top Alzheimer’s disease risk genes^47^, that skips exon two. This isoform was reported as new in our data because it remains unannotated by Ensembl as of June 2023^2^, but it has previously been reported by two groups^48,49^. Notably, the articles identifying this new *TREM2* isoform reported a relative abundance around 10%, corroborating our long-read sequencing results^48,49^. The new isoform for *POLB*—a gene implicated in base-excision repair for nuclear and mitochondrial genomes^50,51^— accounted for 28% of the gene’s expression (**Fig. 3d**). We also discovered 34 new transcripts from medically relevant genes with median CPM between 0 and 0.25 and 32 with median CPM between 0.26 and 1, including new RNA isoforms for *SMN1* and *SMN2* (spinal muscular atrophy^52^; **Supplementary** Fig. 27,28). For interest, we show all medically relevant genes with new RNA isoforms that did not meet our high-confidence threshold in **Supplementary** Fig. 29.The medically relevant genes where we identified multiple new isoforms were *PSENEN* (3; part of the gamma-secretase protein complex^53^), *CPLX2* (2; part of synaptic vesicles and associated with Schizophrenia), and *DGUOK* (2; associated with mitochondrial DNA depletion syndrome-3, including liver and brain effects^54^).

**Fig. 3:**
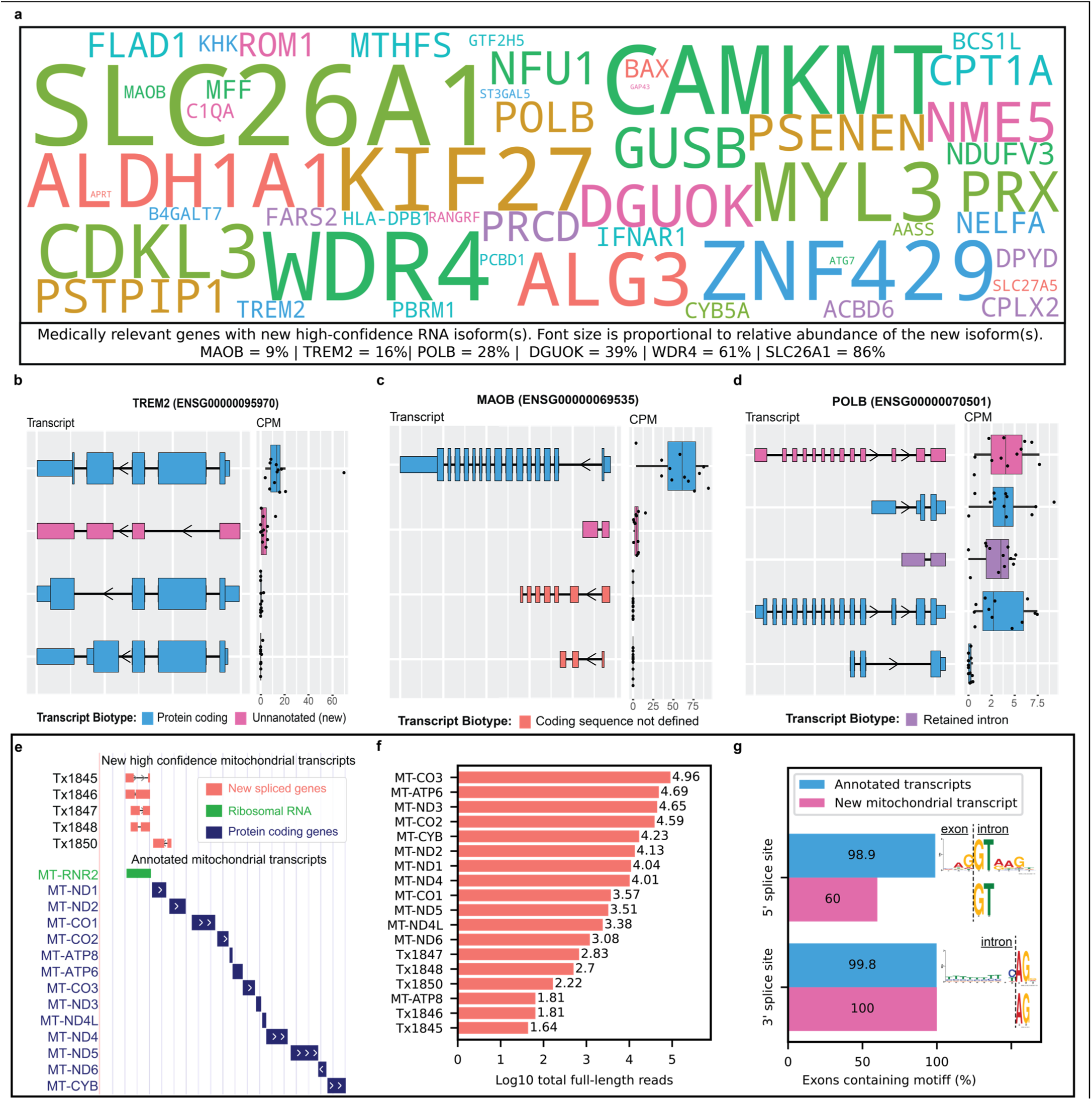
Medically relevant genes with new high-confidence RNA isoforms expressed in human frontal cortex. **a,** Gene names for medically relevant genes where we discovered a new RNA isoform that was not annotated in Ensembl version 107. Only included new RNA isoforms with a median CPM > 1. The size of gene name is proportional to relative abundance of the new RNA isoform. Relative abundance values relevant to this figure can be found in **Supplementary** Fig. 30. **b-d,** RNA isoform structure and CPM expression for isoforms from *TREM2*, *MAOB*, and *POLB*. For *TREM2 and MAOB* all isoforms are shown (4 each). For *POLB* only the top 5 most highly expressed isoforms in human frontal cortex are shown. Figures **e-g** refer to new spliced mitochondrial transcripts, we only included new mitochondrial transcripts with median full-length counts > 40. **e**, Structure for new spliced mitochondrial transcripts in red/coral denoted by “Tx”, MT-RNR2 ribosomal RNA represented in green (overlapping 4 out of 5 spliced mitochondrial isoforms) and known protein coding transcripts in blue. **f**, Bar plot showing number of full-length counts (log10) for new spliced mitochondrial transcripts and known protein coding transcripts. **g**, Bar plot showing the prevalence of canonical splice site motifs for annotated exons from nuclear transcripts with median CPM > 1 versus new exon from spliced mitochondrial transcripts.

### Spliced mitochondrially encoded isoforms

We identified a new set of spliced mitochondrially encoded isoforms containing two exons (**Fig 3e**), a highly unexpected result given that annotated mitochondrial transcripts only contain one exon. New mitochondrial isoforms were filtered using a count threshold based on full-length reads rather than a median CPM threshold due to technical difficulties in quantification arising from the polycistronic nature of mitochondrial transcription. Bambu identified a total of 34 new spliced mitochondrial isoforms, but after filtering using a strict median full-length count threshold of 40, only five high-confidence isoforms remained. Four of the new high-confidence isoforms span the *MT-RNR2* transcript. Not only does *MT-RNR2* encode the mitochondrial 16S ribosomal RNA, but it is also partially translated into an anti-apoptotic 24 amino acid peptide (humanin; HN) by inhibiting the Bax protein^55^. The fifth new high-confidence isoform spans the *MT-ND1* and *MT-ND2* genes, but on the opposite strand. Our results support previous important work by Herai et al. demonstrating splicing events in mitochondrial RNA^56^.

For context, while expression for the new mitochondrial isoforms was low compared to known mitochondrial genes (**Fig. 3f**), their expression is relatively high when compared to all nuclear isoforms. Because we were more stringent in how we estimated expression for the mitochondrial isoforms (using only full-length counts), comparing their overall expression directly to nuclear isoforms is challenging, but all five new high-confidence mitochondrial isoforms are in the top 3,500 most expressed isoforms, including nuclear genome isoforms, when only considering full-length counts (i.e., reads matching all exon-exon boundaries from its assigned transcript). All five exons from new high-confidence mitochondrial isoforms contained the main nucleotides from the canonical 3’ splice site motif (AG), while three out of five (60%) contained the main nucleotides from the canonical 5’ splice site motif (GT) (**Fig. 3g**).

We attempted to validate three new high-confidence mitochondrially encoded isoforms through PCR and successfully validated two of them (**Supplementary** Fig. 25,26). It was not possible to design specific primers for the other two new high-confidence mitochondrial isoforms because of low sequence complexity or overlap with other lowly expressed (low-confidence) mitochondrial RNA isoforms found in our data; thus, we did not attempt to validate them using PCR. While there are more advanced and traditional methods to validate these experimentally, we feel that direct validation of the other spliced mitochondrial isoforms we discovered via PCR, combined with the exceptional and thorough work by Herai et al.^56^ demonstrates that at least some, if not all of those we observed are real, which is our primary objective. Notably, however, we were able to validate all five high-confidence spliced mitochondrial transcripts in the data from Glinos et al.^18^, as each had at least 100 uniquely aligned counts across each of the 5 GTEx brain samples (**Extended Data** Fig. 7). Mitochondria have been implicated in a range of human diseases, including seizure disorders^57^, ataxias^58^, neurodegeneration^59^, and other age related diseases^60^. While function for the new isoforms is not clear, mitochondria are essential to human cell life (and most eukaryotes). Thus, if there are any isoforms that must be fully understood, it is isoforms from the mitochondria because these new spliced mitochondrial isoforms could have important biological roles or serve as biomarkers for mitochondrial function.

### Discovery of transcripts from new gene bodies

RNA isoforms from new gene bodies refer to polyadenylated RNA species coming from regions of the genome where transcription was unexpected (i.e., unannotated). Bambu identified a total of 1,860 isoforms from 1,676 new gene bodies. We observed 1060 new RNA isoforms from new gene bodies with median CPM between 0 and 0.25 and 533 with median CPM between 0.26 and 1, for a total of 1593 potential new gene bodies with a CPM ≤ 1. To err on the side of caution, we consider these potential discoveries as low confidence and excluded them from the remainder of our analyses. After excluding isoforms with a median CPM ≤ 1, 267 high-confidence isoforms from 245 gene bodies remained (**Fig. 4a,b**). Glinos et al. did not specifically report on new gene bodies, but Leung et al. reported 54 new gene bodies in human cortex where 5 overlapped with our high-confidence isoforms from new genes. The new isoforms from new gene bodies had a median length of 1529 nucleotides, ranging between 109 and 5291 nucleotides (**Fig. 4c**). The number of exons ranged between 2 and 4, with 96.6% of isoforms only having two exons (**Fig. 4d**).

**Fig. 4:**
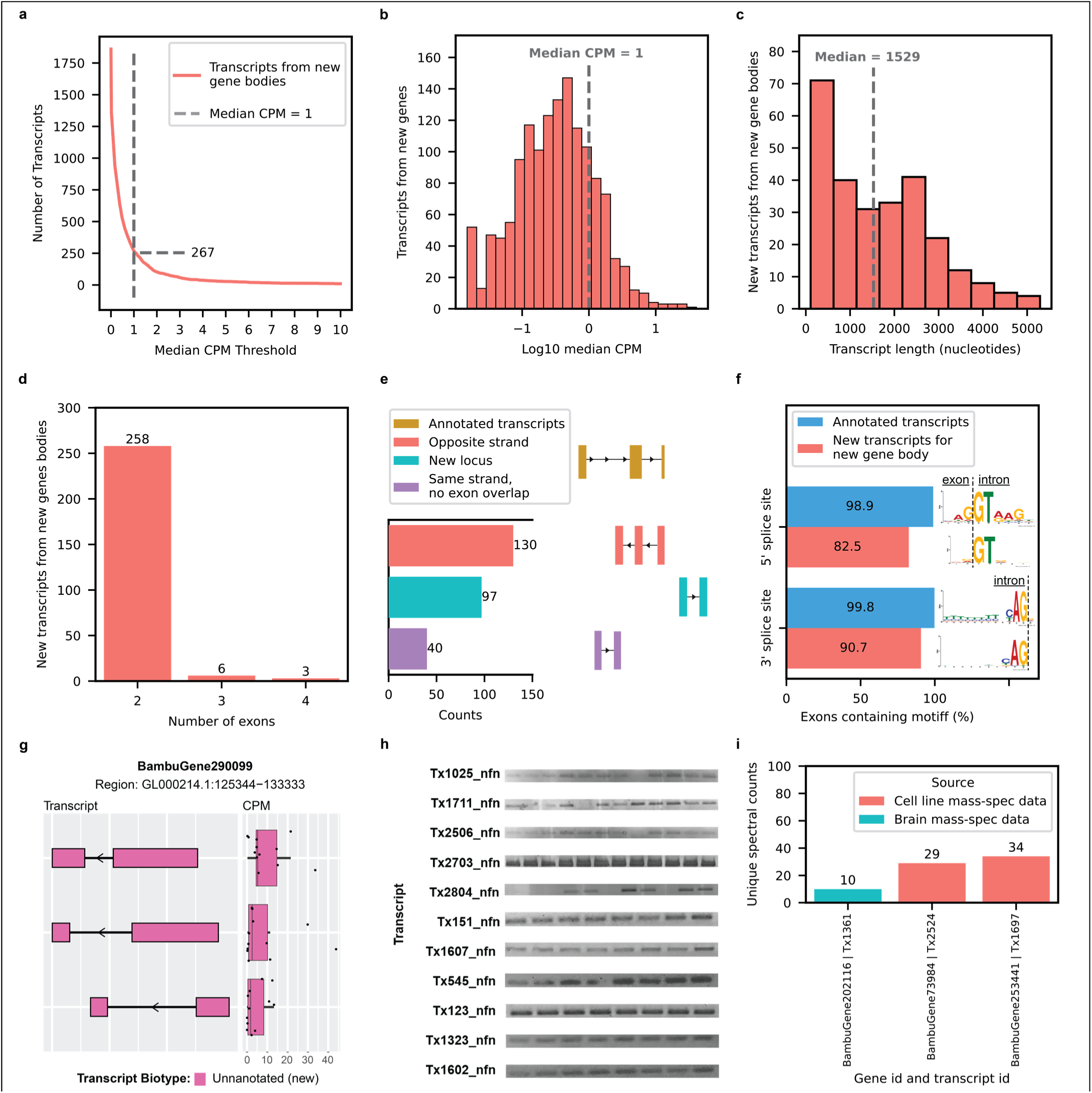
New high-confidence gene bodies in human frontal cortex tissue. **a,** Number of newly discovered transcripts from new gene bodies represented across median CPM threshold. Cutoff shown as the dotted line set at median CPM = 1. **b,** Distribution of log10 median CPM values for new transcripts from new gene bodies, dotted line shows cutoff point of median CPM = 1. Figures **c-g** only include data from transcripts above this expression cutoff. **c,** Histogram showing length distribution for new transcripts from new gene bodies. **d,** Bar plot showing the distribution of the number of exons for new transcripts from new gene bodies. Given the large proportion transcripts containing only two exons, it is possible that we only sequenced a fragment of larger RNA molecules. **e,** Bar plot showing the kinds of events that gave rise to new transcripts from new gene bodies. **f,** Bar plot showing the prevalence of canonical splice site motifs for annotated exons from transcripts with median CPM > 1 versus new exons from new gene bodies. **g,** RNA isoform structure and CPM expression for isoforms from new gene body (*BambuGene290099)*. **h,** Gel electrophoresis validation using PCR amplification for a subset of new isoforms from new genes. This is an aggregate figure showing bands for several different gels. Individual gel figures are available in **Supplementary** Figures 5-26**. i,** Protein level validation using publicly available mass spectrometry proteomics data. Y-axis shows number of spectral counts from uniquely matching peptides (unique spectral counts); new transcripts from new genes were considered validated at the protein level if they had more than 5 unique spectral counts.

Given the large proportion transcripts containing only two exons, it is possible that we only sequenced a fragment of larger RNA molecules. Of the 267 new high-confidence isoforms from new gene bodies, 130 overlapped a known gene body on the opposite strand, 97 came from a completely new locus, 40 came from within a known gene body, but did not overlap a known exon (**Fig. 4e**). These 170 new transcripts from new gene bodies located in intragenic regions could be a result of leaky transcription and splicing. A recent article by Sen et al.^61^ suggests that spurious intragenic transcription may be a feature of aging in mammalian tissues. In new isoforms from new gene bodies, 82.5% (222/269) of exons contained the primary “GT” nucleotides from the canonical 5’ splice site motif, while 90.7% (244/269) contained the primary “AG” nucleotides from the canonical 3’ splice site motif (**Fig. 4f**). Interestingly, the new gene body, *BambuGene290099*, had three high-confidence RNA isoforms (**Fig. 4g**). We attempted to validate 12 new high-confidence RNA isoforms from new gene bodies through PCR and gel electrophoresis and successfully validated 11 of them (**Fig. 4h, Supplementary** Fig. 5-26**, Supplementary Table 4**). The one RNA isoform that failed to validate via standard PCR (no clear band on gel) successfully validated through RT-qPCR (mean Ct = 23.2; **Supplementary Table 5**). MIQE guidelines by Bustin et al.^25^ suggests Ct < 40 as a cutoff for RT-qPCR validation, but we used a more stringent cutoff of CT < 35 to be conservative. All 12 new RNA isoforms from new gene bodies that were validated through PCR and gel electrophoresis and RT-qPCR are unique to this study.

We attempted to validate our new high-confidence transcripts using publicly available long-read RNAseq data from five GTEx^18^ brain samples (Brodmann area 9) and short-read RNAseq data from 251 ROSMAP^21^ brain samples (Brodmann area 9/46). Over 94.4% of the new high-confidence transcripts from new gene bodies had at least one uniquely mapped read in either GTEx or ROSMAP data and over 44.2% had at least 100 uniquely mapped reads in either dataset (**Extended Data** Fig. 7**, Supplementary Table 8)**. The rate of validation was higher for new transcripts from known gene bodies than for new transcripts from new gene bodies, indicating that some of our newly discovered genes could be aging related. Whether these newly discovered gene bodies are biologically meaningful or “biological noise”, is unclear. We validated three RNA isoforms from new gene bodies at the protein level using mass spectrometry data from the same brain region and human cell-lines (**Fig. 4i**); all three are unique to this study.

We also added External RNA Controls Consortium (ERCC) spike-ins to our samples to benchmark RNA quantification. Mean ERCC CPM values had a strong correlation with their known concentrations, achieving a Spearman correlation coefficient of 0.98 (**Supplementary** Fig. 31). Interestingly, we identified a new low abundance RNA isoform (median CPM < 1) with two exons for ERCC RNA spike-ins (**Supplementary** Fig.32). We were skeptical about this discovery since ERCCs only contain one exon, but we validated these results by PCR using two different batches of ERCC mix and several sample library preparations (**Supplementary** Fig. 33,34).

### Medically relevant genes expressing multiple RNA isoforms in human frontal cortex tissue

As part of understanding the function of individual RNA isoforms, we need to understand the context of their expression, including tissue type. Quantifying the relative abundance of specific isoforms within individual tissues is a critical step to understanding their individual functions. Notably, we found 7,042 genes expressing two or more RNA isoforms with a median CPM > 1, where 3,387 genes expressed two or more isoforms with clearly distinct protein sequences (**Fig. 5a,b**). Of the 5,035 medically relevant genes included here^27^ 1,917 expressed multiple isoforms and 1,018 expressed isoforms with different protein coding sequences (**Fig. 5c**), demonstrating the isoform diversity of medically relevant genes in a single tissue and the importance of interpreting genetic variants in the proper context of tissue-specific isoforms. Isoforms with clearly distinct protein sequences are perhaps most likely to have different functions, but we still need to understand the effects of isoforms that are technically different only in noncoding regions (i.e., modified UTR regions can have large impacts on translation^62,63^). In addition, we observed 7,418 transcripts from medically relevant genes expressed with median CPM > 1, where 5,695 are longer than 2,000 nucleotides (**Supplementary** Fig. 35). Given the length of these 5,695 RNA isoforms it is likely that their quantification is less accurate, despite the advantages long-read sequencing offers.

Interestingly, 98 genes implicated in brain related diseases expressed multiple RNA isoforms in human frontal cortex, including Alzheimer’s disease genes such as *APP* (Aβ precursor protein) with five, *MAPT* (tau protein) with four, and *BIN1* with eight (**Fig. 5d,e1-e4**). Notably, we only observed four *MAPT* isoforms with a median CPM > 1, where two were expressed at levels many times greater than the others, while substantial previous research suggests there are six tau proteins expressed in the central nervous system^64–66^. Researchers have been studying these and other top Alzheimer’s disease genes for years, yet there is still a lot to learn about most of their isoforms and their medical relevance. To truly understand how any of the top Alzheimer’s disease genes are involved in disease, we need to understand exactly why brain tissue is actively expressing multiple isoforms of the “same” gene and what functions they serve. Similarly, several genes implicated in other neurogenerative diseases and neuropsychiatric disorders expressed multiple isoforms in human frontal cortex, including *SOD1* (amyotrophic lateral sclerosis, ALS; frontotemporal dementia, FTD; **Fig. 5f1**) with two isoforms expressed with a median CPM > 1, *SNCA* (Parkinson’s disease; **Fig. 5f1**) with four, *TARDBP* (TDP-43 protein; involved in several neurodegenerative diseases; **Fig. 5f1,f2**) with four, and *SHANK3* (autism spectrum disorder; **Fig. 5g1,g2**) with three.

**Fig. 5:**
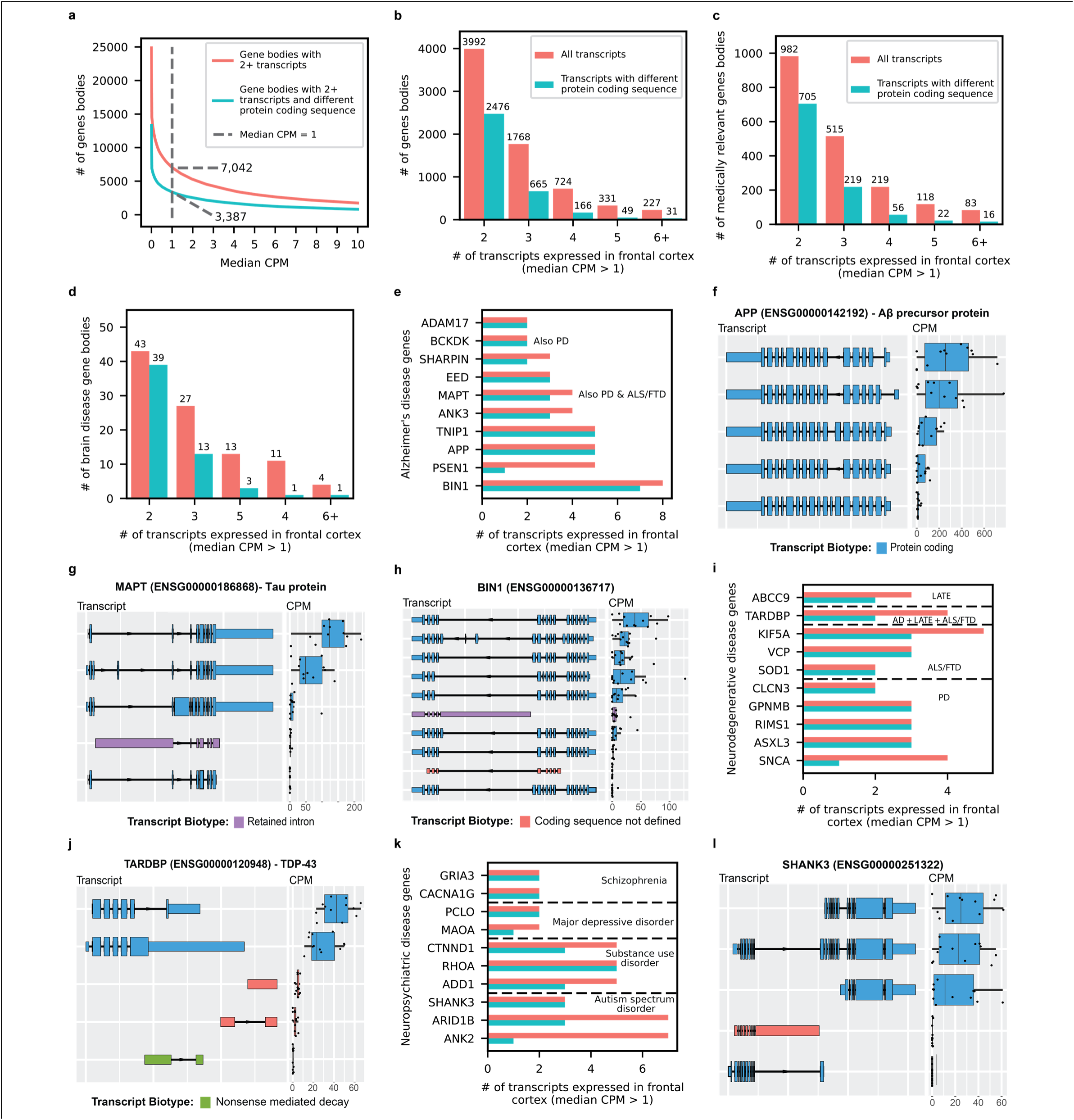
Gene bodies expressing multiple transcripts in human frontal cortex tissue. **a,** Gene bodies with multiple transcripts across median CPM threshold. Dotted line is at median CPM = 1, figures **b-g2** only include gene bodies with multiple transcripts at median CPM > 1. **b,** Gene bodies expressing multiple transcripts. **c,** Medically relevant gene bodies expressing multiple transcripts. **d,** Brain disease relevant gene bodies expressing multiple transcripts. **e1,** Transcripts expressed in frontal cortex for a subset of genes implicated in Alzheimer’s disease. AD: Alzheimer’s disease. ALS/FTD: Amyotrophic lateral sclerosis and frontotemporal dementia. PD: Parkinson’s disease. **e2,** *APP* transcript expression. **e3,** *MAPT* transcript expression. **e4,** *BIN1* transcript expression. **f1,** Same as **e1** but for a subset of genes implicated in other neurodegenerative diseases. LATE: Limbic-predominant age-related *TDP-43* encephalopathy. **f2,** *TARDBP* transcript expression. **g1,** Same as **e1** but for a subset of genes implicated in neuropsychiatric disorders. **g2,** *SHANK3* transcript expression.

### Isoform analysis reveals disease-associated transcriptomic signatures unavailable at gene level

Perhaps the most compelling value in long-read isoform sequencing is the ability to perform differential *isoform* expression analyses. Through these analyses, we can begin to distinguish which isoforms are involved in specific pathways, cell types, tissue types, and ultimately human health and disease, such as Alzheimer’s disease. Thus, as proof of principle, we performed differential gene and isoform expression analyses comparing six pathologically confirmed Alzheimer’s disease cases and six cognitively unimpaired controls. The data set is not large enough to draw firm disease-specific conclusions, but it does demonstrate the need for larger studies.

For the gene-level analysis we included all gene bodies with a median CPM > 1, for a total of 20,448. For the RNA isoform-level analysis, we included all transcripts with median CPM > 1 coming from genes with more than one transcript expressed at CPM > 1, for a total of 19,423 RNA isoforms from 7,042 genes. To be conservative, the threshold for differential expression was set at a relatively strict |log2foldchange| > 1 and FDR corrected p-value (q-value) < 0.05. We found 176 differentially expressed genes and 105 differentially expressed RNA isoforms (**Fig. 6a,b; Supplementary Table 9,10**). Out of these 105 isoforms, 99 came from genes that were not differentially expressed when collapsing all isoforms into a single gene measurement (**Fig. 6a,b**), demonstrating the utility of differential isoform expression analyses. Interestingly, there were two differentially expressed isoforms from the same gene (*TNFSF12)*, with opposite trends. The *TNFSF12-219* isoform was upregulated in Alzheimer’s disease cases while *TNFSF12-203* was upregulated in controls (**Fig. 6c,d,e**), even though the *TNFSF12* gene was not differentially expressed when collapsing all transcripts into a single gene measurement (**Fig. 6c**).

**Fig. 6:**
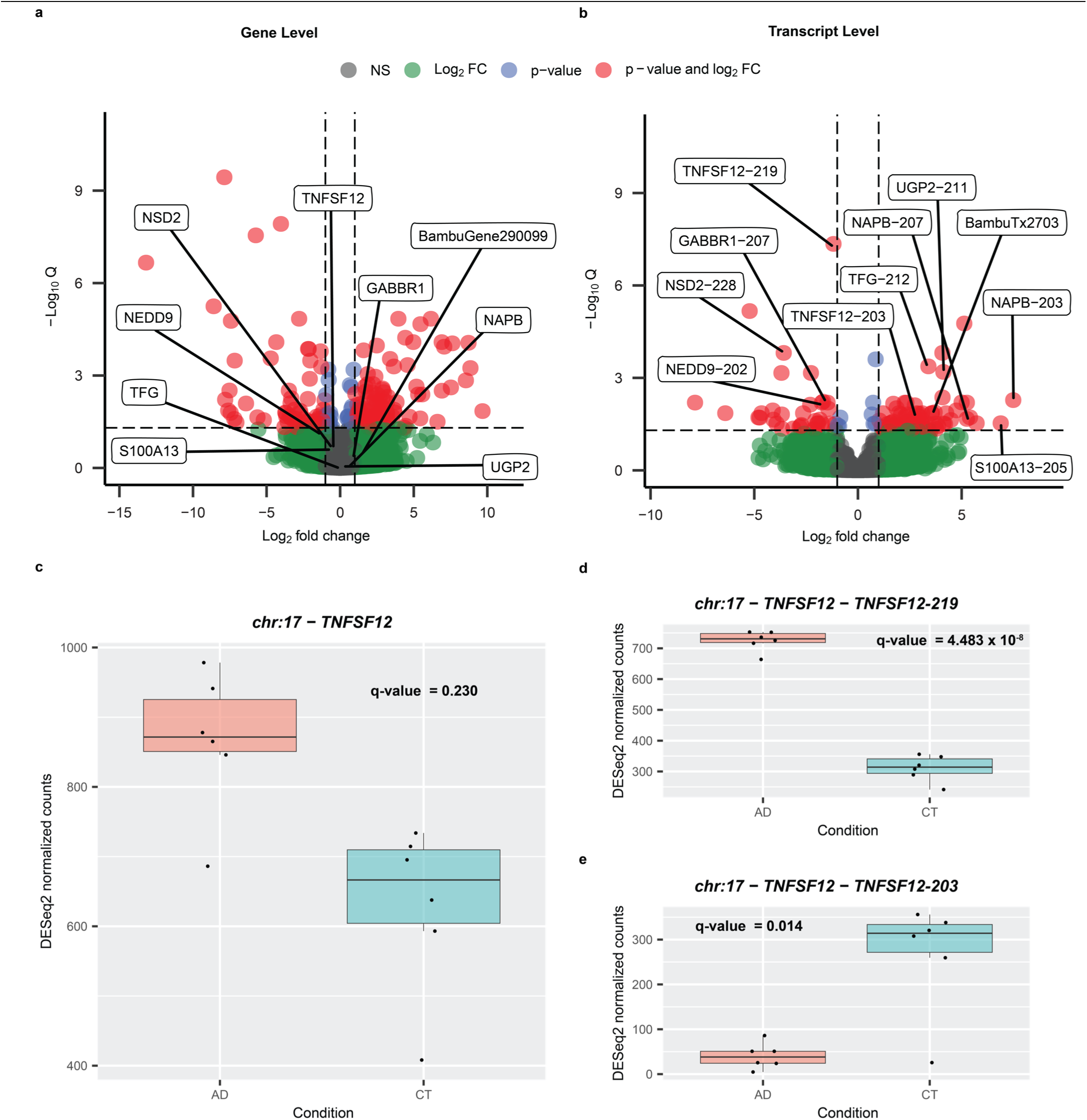
RNA isoform analysis can reveal disease expression patterns unavailable at gene level. **a,** Differential gene expression between Alzheimer’s disease cases and cognitively unimpaired controls. The horizontal line is at FDR corrected p-value (q-value) = 0.05. Vertical lines are at Log2 fold change = -1 and +1. The threshold for differential gene expression was set at q-value < 0.05 and |Log2 fold change| > 1. Names displayed represent a subset of genes that are not differentially expressed but have at least one RNA isoform that is differentially expressed. **b,** Same as **a** but for differential RNA isoform expression analysis. NEDD9-202, NAPB-203, and S100A13-205 are examples of protein coding RNA isoforms that were differentially expressed even though the gene was not. **c,** Expression for *TNFSF12* between Alzheimer’s disease cases and controls. The *TNFSF12* gene does not meet the differential expression threshold. **d,** *TNFSF12-219* transcript expression between cases and controls; *TNFSF12-219* is upregulated in cases. **e,** Expression for the *TNFSF12-203* transcript between cases and controls; *TNFSF12-203* is upregulated in controls. These differentially expressed *TNFSF12* RNA isoforms are not thought to be protein coding, but understanding why cells actively transcribe non-coding RNAs remains an important question in biology. Definition of boxplot elements: center line, median; box limits, upper and lower quartiles; whiskers, 1.5x interquartile range.

The *TNFSF12* gene is a pro-inflammatory cytokine that has been associated with both apoptosis and angiogenesis^67^, suggesting that this “single” gene has multiple distinct functions; it is also part of the Tumor Necrosis Factor super family that has been implicated in Alzheimer’s disease for driving inflammation^68^. These differentially expressed *TNFSF12* RNA isoforms are not thought to be protein coding, but understanding why cells actively transcribe different non-coding RNAs remains an important question in biology. NEDD9-202, NAPB-203, and S100A13-205 are examples of protein coding RNA isoforms that were differentially expressed even though the gene was not. While we cannot make any meaningful conclusions about whether specific genes or isoforms are involved in Alzheimer’s disease (because of sample size), we clearly demonstrate the need to perform larger long-read RNA sequencing studies that perform differential *isoform* expression analysis across human cell types, tissues, and disease.

For interest, we measured the expression patterns for the TNFSF12-203 and TNFSF12-219 isoforms in the 5 GTEx long-read RNAseq samples from Brodmann area 9 to assess whether the expression pattern matched what we observed in our Alzheimer’s disease controls (**Extended Data** Fig. 9). We found that the expression for both TNFSF12 isoforms shows greater variability than either of our groups, but arguably more closely resembles the pattern in our controls. Specifically, the overall trend between isoforms TNFSF12-203 and TNFSF12-219 are relatively close to each other in the GTEx controls, but certainly not as tightly distributed when compared to what we observed in our controls. The expression pattern is also opposite what we observed in our cases; i.e., TNFSF12-219 is higher in our cases and lower in the GTEx controls when compared to TNFSF12-203 within the same group.

Importantly, while we find these results interesting, we emphasize this is only a proof-of-concept study and urge caution not to overinterpret these results. We do not claim, and are not convinced this specific association represents reality in Alzheimer’s disease cases and controls. Equally important, we have no demographic or clinical data on the GTEx controls, including Alzheimer’s disease status, age, sex, genetic background, cause of death, medical history, or other background that could impact the comparison. For interest, we also provide plots from a principal component analysis at both the gene and isoform level where we observed a potential separation between cases and controls (**Supplementary** Fig. 36). We encourage caution to avoid overinterpreting this potential separation between cases and controls given the small sample size, however.

### Web application for human frontal cortex RNA isoform expression

We used the data from this study to create a web application where other researchers can perform queries and see RNA isoform expression in human frontal cortex for any given gene. The web application can be accessed here: https://ebbertlab.com/brain_rna_isoform_seq.html.

## Discussion

We identified hundreds of new human brain-expressed RNA isoforms and new gene bodies, including spliced isoforms in a critical mitochondrial gene (*MT-RNR2*), demonstrating that significant gaps remain in our understanding of RNA isoform diversity (**Fig. 2a,3e,4a**). More importantly, we began to explore RNA isoform function for every gene by quantifying individual RNA isoform expression levels in human frontal cortex. We found 7,042 genes expressing multiple RNA isoforms, with 1,917 being medically relevant genes (i.e., implicated in human disease) (**Fig. 5a,b,c**). Some of these medically relevant genes expressing multiple RNA isoforms in human frontal cortex are implicated in brain-related diseases, including Alzheimer’s disease, Parkinson’s disease, autism spectrum disorder, substance use disorder, and others (**Fig. 5d**). For example, the *APP* gene which encodes the beta-amyloid precursor protein—a defining hallmark of Alzheimer’s disease^69^— expressed 5 different RNA isoforms encoding 5 different proteins (**Fig. 5e2**). The *SHANK3* gene—a major autism spectrum disorder risk gene^70,71^—expressed 4 RNA isoforms encoding 3 different proteins (**Fig. 5g2**). Together, these findings highlight the importance of measuring individual RNA isoform expression accurately so we can start to discern the roles (if any) of each isoform within human health and disease, and so researchers and clinicians alike can better interpret the effects of a given genetic variant. Genetic variants are generally interpreted in the context of a single RNA isoform, and may even be an isoform that is not predominant in the diseased tissue or cell type. Thus, the better we understand the RNA isoform expression landscape, the more prepared researchers and clinicians will be to interpret genetic variants. Additionally, as we better understand the RNA expression landscape and which specific RNA isoforms are involved in disease, those isoforms can be targeted directly.

As proof of concept, we performed differential RNA isoform expression analysis to reveal expression patterns associated with disease that are unavailable when performing gene level analysis (**Fig. 6a,b**). Given the 99 isoforms that were differentially expressed where the gene as a whole was not, we demonstrated that performing differential gene-level expression is important, but may be insufficient in many cases if we want to truly understand the biological complexities, subtleties, and potentially distinct functions afforded by alternative splicing. Indeed, the very existence and breadth of alternative splicing should be a clear signal from biology indicating the importance of accurately characterizing and quantifying individual RNA isoform expression and function. We further suggest that deep long-read RNAseq is necessary to understand the full complexity of transcriptional changes during disease. The gene *TNFSF12* is a key example for the importance of studying RNA isoform expression because the gene itself is not differentially expressed in our data, but in this proof-of-concept study the *TNFSF12-219* isoform is significantly upregulated in Alzheimer’s disease cases while *TNFSF12-203* isoform is significantly upregulated in controls (**Fig. 6c,d,e**). While these isoforms are not thought to be protein coding, understanding why cells actively transcribe non-coding isoforms remains an important question in biology, and biology regularly reveals surprises that scientists do not expect. By characterizing and quantifying expression for all isoforms, we can better determine what functions they have, if any. Findings such as this can lead to new and more precise targets for disease treatment and diagnosis across a broad range of complex human diseases because, rather than targeting an entire gene, we can specifically target the isoform(s) that are either promoting cellular health or that are driving cellular dysfunction.

We also identified five new, high-confidence, and spliced mitochondrially encoded RNA isoforms with two exons each. This is a surprising finding given that all annotated human mitochondrial transcripts only have one exon (**Fig. 2e,f**). We were skeptical of these spliced mitochondrially encoded isoforms, but validated two of them through PCR and gel electrophoresis (**Supplementary** Fig. 25,26) and previous work by Herai et al. in human cell cultures corroborates our findings^56^. To our knowledge, this is the first study identifying spliced mitochondrial RNA isoform expression directly in human tissue. Although their expression was low compared to known mitochondrial protein coding genes, these new isoforms could potentially have important biological roles or serve as biomarkers for mitochondrial function. Given the involvement of mitochondria in many age-related diseases^60^, we believe it is important to determine exactly what function these spliced mitochondrial RNA isoforms play, if any. In other words, given the number of diseases associated with mitochondrial dysfunction, we cannot afford to overlook the potential importance of these new isoforms simply because we do not yet know what function they serve.

### Study limitations

While long-read sequencing can span entire RNA molecules, most reads in this study are not full-length RNA isoforms despite using template-switching reverse transcription and only including reads with primers present on both ends. It is unclear exactly what causes RNAs to not be full-length, but likely culprits include RNA degradation, mechanical shearing during library preparation and sequencing, and limitations in library preparation protocols; working with aged brain tissue has also proven especially challenging. Our data showed a pronounced 3’ bias that can hinder RNA isoform quantification, especially for genes where the exon diversity is closer to the 5’ end (**Supplementary** Fig. 37). Even with RNA integrity scores ≥ 9, our samples averaged 17.5% full-length reads and 42.4% reads uniquely assigned to an RNA isoform (**Extended Data** Fig. 1c); these biases may be specific to the tissues or subjects used in this study, so additional work across tissue types and tissue banks will be important to better characterize the biological and technical limitations of isoform sequencing.

Long-reads present an improvement over short-read RNAseq, but it remains challenging to accurately quantify RNA isoforms in genes with many large and similar isoforms (**<u>Extended Data</u>** Fig. 10). Thus, while this work is a significant improvement over short-read sequencing, the data are not perfect and future improvements in sequencing, transcriptome annotation, and bioinformatic quantification will continue to improve the accuracy of long-read RNA sequencing.

For isoform discovery and quantification, we used bambu^11^, which has been shown to minimize false positives^10,11^. Like all methods there are important caveats to consider when analyzing and interpreting these data. Specifically, as a tradeoff for higher precision, bambu is unable to discover new RNA isoforms that only differ from an annotated RNA isoform because of an alternative transcription start and/or end site (e.g., shortened 5’ UTR). Additionally, the increasing complexity of annotations can impact quantification due to non-unique reads being split between multiple transcripts. For example, if a read maps equally well to two RNA isoforms, each isoform will receive credit for 0.5 reads.

A high-proportion of new transcripts from new gene bodies were intragenic—either on the opposite strand of a known gene, or in the same strand but without exon overlap; this pattern is consistent with potential spurious transcription associated with mammalian tissue aging^61^. The advanced age of subjects included in our study and the lower rate of validation of our newly discovered gene bodies in other datasets suggests that some of these transcripts could be a result of aging, though their biological significance remains unclear. In addition, most new RNA isoforms from new gene bodies only had 2 exons, therefore it is possible that these are fragments from longer RNA molecules.

In this work, our sequencing was specific to prefrontal cortex from aged subjects, thus discoveries and interpretations are generally limited to this tissue and phenotypes associated with this tissue and cohort. Hence, most of our results focused on genes and diseases primarily associated with the central nervous system.

The differential RNA isoform expression analysis performed in this study cannot be used to infer RNA expression patterns associated with Alzheimer’s disease in the general population due to its small sample size. It only serves as a proof of concept for the value of measuring individual RNA isoform expression in disease tissue. We refrained from performing differential isoform usage analysis and pathway analysis to avoid overinterpretation of results from only 12 samples, however, these analyses could provide valuable insights in larger studies. Additionally, this study is based on “bulk” RNA sequencing, rather than single-cell sequencing; bulk sequencing is likely to obscure critical cell type-specific expression patterns that single-cell sequencing can elucidate, though the cost of single-cell sequencing combined with long-read sequencing is still a major hurdle in making a large study of this kind feasible.

While presenting this work at conferences, some were skeptical regarding the idea that the various RNA isoforms a given gene expresses are performing distinct functions, and some even suggested the large number of isoforms observed across various studies are simply biological noise. While we agree that we should be cautious not to overstate or assume that most of the isoforms a given gene body expresses are performing distinct functions (even if closely related), we are equally convinced of the importance of fully characterizing and quantifying all RNA isoforms across tissues and cell types to help determine whether individual isoforms do have unique functions, even if, and perhaps especially if the differences are biologically subtle.

In our opinion, while assuming most isoforms have a distinct function would be irresponsible, we also think it would be a mistake to assume that most of the isoforms are simply noise. Assuming that most isoforms are noise may be akin to assuming alternative splicing is simply a biological accident rather than an evolutionarily evolved biological process that enables biological diversity and complexity^72–74^. In our opinion, humans and other complex organisms clearly require a remarkable level of biological complexity, and to assume that such complexity across a vast range of tissue and cell types, combined with a vast range of developmental and aging stages is possible from a mere 20,000+ protein-coding genes all performing a single function seems unlikely. It may be that any differences in function are subtle, but these subtleties may be important to understand variation in various phenotypes (e.g., response to vaccines or drug treatments) that remain a mystery. Nevertheless, only a deeper interrogation into how individual isoforms play into biology across tissues, cell types, and developmental stages will ultimately reveal the reality.

## Conclusion

Individual RNA isoform expression has been overlooked due to technical limitations of short-read sequencing. Now with long-reads we can more accurately quantify the complete set of RNA isoforms. More importantly, we can start to understand the role of individual RNA isoforms from understudied genes and isoforms in the development and progression of diseases, along with what role they play in various developmental and aging stages. We demonstrate a large proportion of medically relevant genes express multiple RNA isoforms in human frontal cortex, with many encoding different protein coding sequences that could potentially perform different functions. As proof of principle, we demonstrate that differential RNA isoform analysis can reveal transcriptomic signatures in Alzheimer’s disease that are not available at the gene level. While we cannot generalize these data to infer differential isoform expression across all Alzheimer’s disease cases and controls due to the small sample size, we believe it is worth investigating in a larger cohort. In the long-term, we believe that long-read RNAseq will be the best tool to assess RNA expression patterns in complex human diseases and to identify new molecular targets for treatment and diagnosis.

## Methods

### Sample collection, RNA extraction, and quality control

Frozen post-mortem human frontal cortex brain samples were collected at the University of Kentucky Alzheimer’s Disease Research Center autopsy cohort^75^, snap-frozen in liquid nitrogen at autopsy and stored at -80°C. Postmortem interval (from death to autopsy) was <5hrs in all samples. All samples came from Caucasian individuals. Approximately 25mg of grey matter from the frontal cortex was chipped on dry ice into prechilled 1.5ml Low bind tubes (Eppendorf, catalogue number: 022431021), kept frozen throughout the process and stored at -80oC. RNA was extracted using the Lexogen SPLIT RNA extraction kit (Lexogen, catalog number: 008.48) using protocol version 008UG005V0320 (**Supplementary File 1**).

Briefly, ∼25mg of tissue was removed from -80oC storage and kept on dry ice until processing began. 400 μL chilled isolation buffer (4°C), (Lexogen SPLIT RNA kit) was added to each tube and the tissue homogenized using a plastic pestle (Kontes Pellet Pestle, VWR, catalogue number: KT749521-1500). Samples remained on ice to maintain RNA integrity while other samples were homogenized. Samples were then decanted into room-temperature phase lock gel tubes, 400 ul of chilled phenol (4°C) added and the tube inverted five times by hand. 150 ul acidic buffer (AB, Lexogen) was added to each sample, the tube inverted five times by hand before 200ul of chloroform was added and inverted for 15 seconds. After a two-minute incubation at room temperature, samples were centrifuged for 2 minutes at 12,000g at 18-20oC and the upper phase (approx. 600ul) was decanted in a new 2ml tubes. Total RNA was precipitated by the addition of 1.75x the volume of isopropanol to sample and then loaded onto a silica column by centrifugation (12,000g, 18oC for 20s) with the flow through being discarded. The column was then washed twice with 500 ul of isopropanol and then three times with 500 ul wash buffer (Lexogen) with the column being centrifuged (12,000g, 18oC for 20s) and the flow through being discarded each time. The column was transferred to a new low bind tube, and the RNA eluted by the addition of 30 ul of elution buffer (incubated for one minute and then centrifuged at 12,000g, 18oC for 60s) and the eluted RNA immediately placed on ice to prevent degradation.

RNA quality was determined initially by nanodrop (A260/A280 and A260/A230 ratios) and then via Agilent Fragment Analyzer 5200 using the RNA (15nt) DNF-471 kit (Agilent). All samples achieved Nanodrop ratios of >1.8 and Fragment Analyzer RIN >9.0 prior to sequencing and indicated high quality RNA. (**Supplementary** Fig. 38-49**; Supplementary Table 1**).

### RNA Spike-ins

ERCC RNA Spike-In Controls (Thermofisher, catalogue number: 4456740) were employed to establish a consistent baseline measurement of RNA, ensuring standardization within each experiment and across multiple experiments. ERCC controls were added to the RNA at the point of starting cDNA sample preparation at a final dilution of 1:1000.

### Library preparation, sequencing, and base-calling

Isolated RNA was kept on ice until quality control testing was completed as described above. Long read cDNA library preparation commenced, utilizing the Oxford Nanopore Technologies PCR-amplified cDNA kit (Oxford Nanopore Technologies, catalogue number: SQK-PCS111). The protocol, though not available due to a legal embargo, was followed, with two notable modifications being that in the cDNA amplification PCR the expansion time was set to 6 minutes and the PCR was 14 amplification cycles. We used the Oxford Nanopore PCS111 kit reverse transcription adapters to select polyA RNAs from our total RNA extraction. Therefore, our polyA enrichment step happens at the beginning of the cDNA synthesis in this protocol. cDNA quality was determined using Agilent Fragment Analyzer 5200 and Genomic DNA (50kb) kit (Agilent DNF-467). See **Supplementary** Figures 50-61 for cDNA traces. The libraries underwent sequencing on the PromethION platform with flow cell R9.4.1. Each sample was allocated one full PromethION flow cell. Sequencing was performed continuously for approximately 60 hours. The Fast5 files obtained were basecalled using the Guppy GPU basecaller version 6.3.9 with configuration dna_r9.4.1_450bps_hac_prom.cfg.

### Read pre-processing, genomic alignment & quality control

Nanopore long-read sequencing reads were pre-processed using pychopper^76^ version 2.7.2 with the PCS111 sequencing kit setting. Pychopper filters out any reads not containing primers on both ends and rescues fused reads containing primers in the middle. Pychopper then orients the reads to their genomic strand and trims the adapters and primers off the reads.

The pre-processed reads were then aligned to the GRCh38 human reference genome (without alternate contigs) with added ERCC sequences using minimap2^77^ version 2.22-r1101 with parameters “-ax splice -uf”. Full details and scripts are available on our GitHub (**Code Availability**). Aligned reads with a mapping quality (MAPQ) score < 10 were excluded using samtools^78^ version 1.6. The resulting bam alignment files were sorted by genomic coordinate and indexed before downstream analysis. Quality control reports and statistics were generated using PycoQC^79^ version 2.5.2. Information about mapping rate, read length, and other sequencing statistics can be found in **Supplementary Table 1** and **Supplementary** Fig. 1-4.

### Analysis using CHM13 reference

We processed the RNAseq data from the 12 dorsolateral pre-frontal cortex samples from this study using the same computational pipeline, but used the CHM13 reference genome rather than GRCh38. We also set bambu to quantification only mode rather than quantification and discovery. The reference fasta and gff3 files were retrieved from the T2T-CHM13 GitHub (https://github.com/marbl/CHM13). Here is the link to the reference genome sequence (https://s3-us-west-2.amazonaws.com/human-pangenomics/T2T/CHM13/assemblies/analysis_set/chm13v2.0.fa.gz) and the GFF3 annotation (https://s3-us-west-2.amazonaws.com/human-pangenomics/T2T/CHM13/assemblies/annotation/chm13.draft_v2.0.gene_annotation.gff3). We did this as a preliminary analysis to assess whether the extra 99 predicted protein coding genes from CHM13 reported in Nurk et. al^26^ are expressed in human frontal cortex.

### Transcript Discovery and Quantification

Filtered aligned reads were utilized for transcript quantification and discovery using bambu^11^ version 3.0.5. We ran bambu using Ensembl^2^ version 107 Gene transfer format (GTF) annotation file with added annotations for the ERCC spike-in RNAs and the GRCh38 human reference genome sequence with added ERCC sequences. Sample BAM files were individually pre-processed with bambu, and the resulting 12 files were provided as input all at once to perform transcript discovery and quantification using bambu The Novel Discovery Rate (NDR) was determined based on the recommendation by the bambu machine learning model. The model recommended a novel discovery rate threshold of 0.288. Bambu outputs three transcript level counts matrices, including total counts (all counts including reads that were partially assigned to multiple transcripts), unique counts (only counts from reads that were assigned to a single transcript), and full-length reads (only counts from reads containing all exon-exon boundaries from its respective transcript). Except where specified otherwise, expression values reported in this article come from the total counts matrix.

We used full-length reads for quantification in the mitochondria since the newly discovered spliced mitochondrial transcripts caused issues in quantification. Due to polycistronic mitochondrial transcription, many non-spliced reads were partially assigned to spliced mitochondrial transcripts, resulting in a gross overestimation of spliced mitochondrial transcript expression values. We were able to fix this issue by using full-length counts—only counting reads that match the exon-exon boundaries of newly discovered spliced mitochondrial transcripts.

We only included newly discovered (i.e., unannotated) transcripts with a median counts per millions (CPM) > 1 in downstream analysis (i.e., high-confidence new transcripts). New transcripts from mitochondrial genes were the exception, being filtered using a median full-length reads > 40 threshold. Due to the polycistronic nature of mitochondrial transcription, reads from overlapping transcripts were being erroneously assigned to new spliced mitochondrial transcripts, resulting in large misestimations of expression. To address this error, we only used full-length reads to filter out new spliced mitochondrial transcripts with low expression. We also used full-length counts to compare spliced mitochondrial transcript expression levels to expression from annotated protein coding mitochondrial transcripts.

Data from transcriptomic analysis can be visualized in the web application we created using R version 4.2.1 and Rshiny version 1.7.4 . Link to web application: https://ebbertlab.com/brain_rna_isoform_seq.html

### Subsampling discovery analysis

Nanopore long-read sequencing data was randomly subsampled into 20% increments, generating the following subsamples for each sample: 20%, 40%, 60%, and 80%. The 12 subsampled samples for each increment were ran through our long-read RNAseq discovery and quantification pipeline described under **<u>Read pre-</u> <u>processing, genomic alignment & quality control</u>** and **<u>Transcript Discovery and Quantification.</u>** We compared the number of discovered transcripts between the subsamples and the full samples to assess the effect of read depth on the number of transcripts discovered using bambu.

### Transcript discovery using publicly available GTEx data with Bambu

We obtained the long-read RNAseq data from 90 GTEx samples across 15 human tissues and cell-lines sequenced with the Oxford Nanopore Technoligies PCR amplified cDNA protocol (PCS109) generated by Glinos et al.^18^ . We then processed these data through our long-read RNAseq discovery and quantification pipeline described under **<u>Read pre-processing, genomic alignment & quality control</u>** and **<u>Transcript</u> <u>Discovery and Quantification.</u>** We used the same ENSEMBL version 88 annotations originally used in Glinos et al. and compared the results with the original Glinos et al. publication results and results from our data to assess the effect of the isoform discovery tool (i.e., bambu^11^ vs. FLAIR^23^) on the number of newly discovered transcripts. We also compared the number of newly discovered transcripts when running GTEx data through our computational pipeline with the ENSEMBL version 88 annotation and the ENSEMBL 107 annotation to assess the effect of different annotations in the number of transcripts discovered. Lastly, we compared the overlap between new transcripts from known genes discovered in our study using 12 brain samples with the original Glinos et al. publication results and the results we obtained from running the GTEx data through our computational pipeline using the ENSEMBL version 107 annotations.

### Validation of new transcripts using publicly available GTEx data

We obtained publicly available GTEx nanopore long-read RNAseq data from 6 brain samples (Brodmann area 9). One of the samples was excluded because it had less than 50,000 total reads, therefore, five samples were used for all downstream analysis. These data had been previously analyzed in Glinos et. al^18^. Fastq files were pre-processed using pychopper^76^ version 2.7.2 with the PCS109 sequencing kit setting. The pre-processed reads were then aligned to the GRCh38 human reference genome (without alternate contigs) with added ERCC sequences using minimap2^77^ version 2.22-r1101 with parameters “-ax splice -uf”. Full details and scripts are available on our GitHub (**Code Availability**). Aligned reads with a mapping quality (MAPQ) score < 10 were excluded using samtools^78^ version 1.6. The resulting bam alignment files were sorted by genomic coordinate and indexed before downstream analysis. Bam files were utilized for transcript quantification only (no discovery) using bambu^11^ version 3.0.5. We ran bambu using Ensembl^2^ version 107 Gene transfer format (GTF) annotation file with added annotations for the ERCC spike-in RNAs and all the new transcripts discovered in this study. We provided the GRCh38 human reference genome sequence with added ERCC sequences to run bambu. The transcript level unique counts matrix outputted by bambu was utilized for cross-validation of newly discovered transcripts in this study.

### Validation of new transcripts using publicly available ROSMAP data

We obtained publicly available ROSMAP Illumina 150bp paired-end RNAseq data from 251 brain samples (Brodmann area 9/46). These data had been previously analysed in Mostafavi et. al^21^. Fastq files were pre-processed and quality controlled using trim galore version 0.6.6. We generated the reference transcriptome using the GTF annotation file containing all transcripts from ENSEMBL 107, the ERCC spike-in RNAs, and all the new transcripts discovered in this study. We used this annotation in combination with the GRCh38 reference genome and the tool gffread version 0.12.7 to generate our reference transcriptome for alignment. The pre-processed reads were then aligned to this reference transcriptome using STAR^80^ version 2.7.10b. Full details and scripts are available on our GitHub (**Code Availability**). Aligned reads with a mapping quality (MAPQ) score < 255 were excluded using samtools^78^ version 1.6. We aligned to the transcriptome and filtered reads with MAPQ < 255 to ensure that we would only have reads that uniquely aligned to a transcript for downstream analysis and cross-validation. We quantified the number of uniquely aligned reads using salmon^81^ version 0.13.1. The counts matrix containing uniquely aligned read counts outputted by salmon was utilized for cross-validation of newly discovered transcripts in this study.

### Splice site motif analysis

We utilized the online meme suite tool^82^ (https://meme-suite.org/meme/tools/meme) to create canonical 5’ and 3’ splice site motifs and estimate the percentage of exons containing these motifs. For known genes we only included exons from multi-exonic transcripts that were expressed with a median CPM > 1 in our samples. If two exons shared a start site or an end site, one of them was excluded from the analysis. For new high-confidence transcripts, we filtered out any exon start or end sites contained in the Ensembl annotation. If two or more exons shared a start site or an end site, we only used one of those sites for downstream analyses. For the 5’ splice site analysis we included the last three nucleotides from the exon and the first six nucleotides from the intron. For the 3’ splice site analysis we included the last 10 nucleotides from the intron and the first three nucleotides from the exon. The coordinates for 5’ and 3’ splice site motifs were chosen based on previous studies^83,84^. The percentage of exons containing the canonical 5’ splice site motif was calculated using the proportion of 5’ splice site sequences containing GT as the two last nucleotides in the intron. The percentage of exons containing the canonical 3’ splice site motif was calculated by taking the proportion of 3’ splice site sequences containing AG as the first two nucleotides in the intron. Fasta files containing 5’ splice site sequences from each category of transcript (1. known transcript from known gene body, 2. new transcript from known gene, 3. new transcript from new gene body, 4. new transcript from mitochondrial gene body) were individually submitted to the online meme suite tool to generate splice site motifs. The same process was repeated for 3’ splice site sequences. Due to the small number of transcripts, it was not possible to generate reliable splice site motif memes for new transcripts from mitochondrial transcripts, we instead just used the 5’ GT sequence and 3’ AG sequence to represent them in **Fig. 2g**.

### Comparison between annotations

Annotations from new high-confidence transcripts discovered in this study were compared to annotations from previous studies using gffcompare^85^ version 0.11.2. Transcripts were considered overlapping when gffcompare found a complete match of the exon-exon boundaries (i.e., intron chain) between two transcripts. The annotation from Glinos et al.^18^ was retrieved from: https://storage.googleapis.com/gtex_analysis_v9/long_read_data/flair_filter_transcripts.gtf.gz. The annotation from Leung et al.^19^ was retrieved from: https://zenodo.org/record/7611814/preview/Cupcake_collapse.zip#tree_item12/HumanCTX.collapsed.gff.

### Differential gene expression analysis

Although bambu outputs a gene level counts matrix, this matrix includes intronic reads. To create a gene level counts matrix without intronic read, we summed the transcript counts within each gene using a custom Python script (see code on GitHub; Python version 3.10.8). This gene level counts matrix without intronic reads was used for all gene-level analysis in this article. We only performed differential gene expression analysis on genes with a median CPM > 1 (20,448 genes included in the analysis). The counts matrix for genes with CPM > 1 was loaded into R version 4.2.2. We performed differential gene expression analysis with DESeq2^86^ version 1.38.3 using default parameters. Differential gene expression analysis was performed between AD samples and cognitively unimpaired controls. We set the threshold for differential expression at |log2 fold change| > 1 and FDR corrected p-value (q-value) < 0.05. Detailed descriptions of statistical analysis results can be found on **Supplementary Table 9**. DESeq2 utilizes the Wald test for statistical comparisons.

### Differential isoform expression analysis

For differential isoform expression analysis, we used the transcript counts matrix output by bambu. We only performed differential isoform expression analysis on transcripts with a median CPM > 1 coming from genes expressing two or more transcripts with CPM > 1 (19,423 transcripts from 7,042 genes included in the analysis). This filtered counts matrix was loaded into R version 4.2.2. We performed differential isoform expression analysis with DESeq2 version 1.38.3 using default parameters. Differential isoform expression analysis was performed using the same methods as the gene-level analysis, comparing AD samples and cognitively unimpaired controls, including the same significance thresholds (|log2 fold change| > 1 and FDR corrected p-value < 0.05). Detailed descriptions of statistical analysis results can be found on **Supplementary Table 10**. DESeq2 utilizes the Wald test for statistical comparisons.

### Figures and tables

Figures and tables were generated using custom R (version 4.2.2) scripts and custom Python (version 3.10.8). We used the following R libraries: tidyverse (version 1.3.2), EnhancedVolcano (version 1.18.0), DESeq2 (version 1.38.3), and ggtranscript^87^ (version 0.99.3). We used the following Python libraries: numpy (version 1.24.1), pandas (version 1.5.2), regex (version 2022.10.31), matplotlib (version 3.6.2), seaborn (version 0.12.2), matplotlib_venn (version 0.11.7), wordcloud (version 1.8.2.2), plotly (version 5.11.0), notebook (version 6.5.2). See **Code availability** for access to the custom scripts used to generate figures and tables.

### PCR primer design

We used the extended annotations output by bambu to create a reference transcriptome for primer design. This extended annotation contained information for all transcripts contained in Ensembl version 107 with the addition of all newly discovered transcripts by bambu (without applying a median CPM filter). This annotation was converted into a transcriptome sequence fasta file using gffread (version 0.12.7) and the GRCh38 human reference genome. We used the online NCBI primer design tool (https://www.ncbi.nlm.nih.gov/tools/primer-blast/) to design primers. We utilized default settings for the tool, however, we provided transcriptome described above as the custom database to check for primer pair specificity. We only moved forward with validation when we could generate a primer pair specific to a single new high-confidence transcript. Detailed information about the primers generated can be found in **Supplementary Table 4.**

### PCR and gel electrophoresis Validations

New isoform and gene validations were conducted using PCR and gel electrophoresis. For this purpose, 2ug of RNA was transcribed into cDNA using the High-Capacity cDNA Reverse Transcription kit (AB Applied Biosystems, catalogue number: 4368814) following the published protocol. The resulting cDNA was quantified using a nanodrop, and its quality was assessed using the Agilent Fragment analyzer 5200 with the DNA (50kb) kit (Agilent, DNF-467). Next, 500ng of the cDNA was combined with primers specific to the newly identified isoforms and genes (**Supplementary Table 4**). The amplification was performed using Invitrogen Platinum II Taq Hot start DNA Polymerase (Invitrogen 14966-005) in the Applied Biosystem ProFlex PCR system. The specific primer sequences, annealing temperatures, and the number of PCR cycles are detailed in **Supplementary Table 4**. After the PCR amplification, the resulting products were analyzed on a 1% agarose TAE gel containing 0.5ug/ml ethidium bromide. The gel was run for 30 minutes at 125v, and the amplified cDNA was visualized using a UV light source. Gels from PCR validation for each transcript can be found in **Supplementary** Fig. 5-26**,33,34.**

### Real-time quantitative PCR (RT-qPCR) Validations

The real-time quantitative PCR (RT-qPCR) assays were performed using the QuantStudieo 5™ Real-Time PCR System (Applied Biosystems). Amplifications were carried out in 25 μL reaction solutions containing 12.5 μL 2x PerfeCTa SYBR green SuperMix (2X) (Quantabio Cat# 95054-500), 1.0 μL first-stranded cDNA, 1 ul of each specific primer (10mM, see **Supplementary Table 5**) and 9.0 μL ultra-pure, nuclease free water. RT-qPCR conditions involved an initial hold stage: 50*o*C for 2 mins followed by 95 °C for 3 min with a ramp of 1.6*o*C/s followed by PCR stage of 95 °C for 15 s and 60 °C for 60 s for a total of 50 cycles.). MIQE guidelines by Bustin et al.^25^ suggests Ct < 40 as a cutoff for RT-qPCR validation, but we used a more stringent cutoff of Ct < 35 to be conservative. This means that we only considered a new RNA isoform to be validated by RT-qPCR if the mean Ct value for our samples was below 35. We only attempted to validate new RNA isoforms through RT-qPCR if they first failed to be validated through standard PCR and gel electrophoresis. We did this because RT-qPCR is a more sensitive method, allowing us to validate RNA isoforms that are less abundant or that are harder to amplify through PCR.

In additions, we performed quantification of new and known RNA isoforms from the following genes: *SLC26A1*, *MT-RNR2*, and *MAOB* (**Supplementary Table 6,7**). We followed recommendations by Penna et al.^88^ and used the CYC1 as the gene for Ct value normalization in our human postmortem brain samples. To allow for comparison between different isoforms from the same gene, we used *2*^-Δ*Ct*^ as the expression estimate instead of the more common *2*^-ΔΔ*Ct*^ expression estimate. This is because the *2*^-ΔΔ*Ct*^ expression estimate is optimized for comparisons between samples within the same gene/isoform but does not work well for comparison between genes/isoforms. On the other hand, the *2*^-Δ*Ct*^ expression estimate allows for comparison between different genes/isoforms. RNA isoform relative abundance for RT-qPCR and long-read RNAseq was calculated as follows:

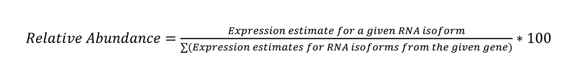

### Proteomics analysis

We utilized publicly available tandem mass spectrometry data from round two of the Religious Orders Study and Rush Memory and Aging Project (ROSMAP) brain proteomics study, previously analyzed by Johnson et al.^89^ and Higginbotham et al^90^. We also utilized publicly available deep tandem mass spectrometry data from six human cell lines, processed with six different proteases and three tandem mass spectrometry fragmentation methods, previously analyzed by Sinitcyn et al^91^. This cell line dataset represents the largest human proteome with the highest sequence coverage ever reported. We started the analysis by creating a protein database containing the predicted protein sequence from all three reading frames for the 700 new high-confidence RNA isoforms we discovered, totaling 2100 protein sequences. We translated each high-confidence RNA isoform in three reading frames using pypGATK^92^ version 0.0.23. We also included the protein sequences for known protein coding transcripts that came from genes represented in the 700 new high-confidence RNA isoforms and had a median CPM > 1 in our RNAseq data. We used this reference protein fasta file to process the brain and cell-line proteomics data separately using FragPipe^93–99^ version 20.0—a Java-based graphical user interface that facilitates the analysis of mass spectrometry-based proteomics data by providing a suite of computational tools. Detailed parameters used for running FragPipe can be found on GitHub and Zenodo, see **Code Availability** and **Data Availability**.

Mass spectrometry suffers from a similar issue as short-read RNAseq, only being able to detect relatively short peptides that do not cover the entire length of most proteins. This makes it challenging to accurately detect RNA isoforms from the same gene. To avoid false-discoveries, we took measures to ensure that we would only consider an RNA isoform to be validated at the protein level if it had peptide hits that are unique to it (i.e., not contained in other known human proteins). We started by taking the FragPipe output and only keeping peptide hits that mapped to only one of the proteins in the database. We then ran the sequence from those peptides against the database we provided to FragPipe to confirm that they were truly unique. Surprisingly, a small percentage of peptide hits that FragPipe reported as unique were contained in two or more proteins in our database, these hits were excluded from downstream analysis. We then summed the number of unique peptide spectral counts for every protein coming from a new high-confidence RNA isoform. We filtered out any proteins with less than six spectral counts. We took the peptide hits for proteins that had more than 5 spectral counts and used online protein-protein NCBI blast tool (blastp)^100^ to search it against the human RefSeq protein database. We used loose thresholds for our blast search to ensure that even short peptides matches would be reported. A detailed description of the blast search parameters can be found on Zenodo. Spectral counts coming from peptides that had a blast match with 100% query coverage and 100% identity to a known human protein were removed from downstream analysis. We took the remaining spectral counts after the blast search filter and summed them by protein id. Proteins from high-confidence RNA isoforms that had more than five spectral counts after blast search filter were considered validated at the protein level. This process was repeated to separately analyze the brain mass spectrometry data and the cell-line mass spectrometry data.

### Reproducibility

Read pre-processing, alignment, filtering, transcriptome quantification and discovery, and quality control steps for Nanopore and Illumina data were implemented using custom NextFlow pipelines. NextFlow enables scalable and reproducible scientific workflows using software containers^101^. Singularity containers were used for most of the analysis in this paper, except for website creation and proteomics analysis due to feasibility issues. Singularity containers enable the creation and employment of containers that package up pieces of software in a way that is portable and reproducible^102^. Instructions on how to access the singularity containers can be found in the GitHub repository for this project. When not forbidden by legal embargo, manufacturer protocols were included as supplementary material. Any changes to standard manufacturer protocols were detailed in **Methods**. All code used for analysis in this article is publicly available on GitHub, see **Code Availability.** All raw data, output from long-read RNAseq and proteomics pipelines, references, and annotations are publicly available, see **Data availability.** Long-read RNAseq results from this article can be easily visualized through this web application: https://ebbertlab.com/brain_rna_isoform_seq.html

## Data availability

Raw long-read RNAseq data generated and utilized in this manuscript are publicly available in Synapse: https://www.synapse.org/#!Synapse:syn52047893.

Raw long-read RNAseq data generated and utilized in this manuscript are also publicly available in NIH SRA (accession number: SRP456327): https://trace.ncbi.nlm.nih.gov/Traces/?view=study&acc=SRP456327

Output from long-read RNAseq and proteomics pipelines, reference files, and annotations are publicly available here: https://doi.org/10.5281/zenodo.8180677

Long-read RNAseq results from this article can be easily visualized through this web application: https://ebbertlab.com/brain_rna_isoform_seq.html

Raw cell-line deep proteomics data used utilized in this article are publicly available here: https://proteomecentral.proteomexchange.org/cgi/GetDataset?ID=PXD024364

Raw brain proteomics data from round 2 of the ROSMAP TMT study are publicly available here: https://www.synapse.org/#!Synapse:syn17015098

GTEx long-read RNAseq data used for validation of our study results is available here: https://anvil.terra.bio/#workspaces/anvil-datastorage/AnVIL_GTEx_V9_hg38

ROSMAP short-read RNAseq data used for validation of our study results is available here: https://www.synapse.org/#!Synapse:syn21589959

## Code availability

All code used in the manuscript is available at: https://github.com/UK-SBCoA-EbbertLab/brain_cDNA_discovery

## Contributions

BAH, JB, and ME developed and designed the study and wrote the manuscript. BAH, MP, BW, KD, MW, EF, and AS performed all analyses. MP developed the RShiny app and KD embedded the RShiny app into ebbertlab.com. JB, KN, LG, GF, PD, SG, EG, RW, and SME helped generate sequencing and supporting data. NS, EF, and AS generated and advised on proteomics analyses. PN provided the invaluable brain samples and pathology. JF, MR, and JM provided important intellectual contributions.

## Competing interests

The authors report no competing interests.

## Funding

This work was supported by the National Institutes of Health [R35GM138636, R01AG068331 to M.E., and 5R50CA243890 to S.G.], the BrightFocus Foundation [A2020161S to M.E.], Alzheimer’s Association [2019-AARG-644082 to M.E.], PhRMA Foundation [RSGTMT17 to M.E.]; Ed and Ethel Moore Alzheimer’s Disease Research Program of Florida Department of Health [8AZ10 and 9AZ08 to M.E., and 6AZ06 to J.F.]; and the Muscular Dystrophy Association (M.E.).

## Acknowledgments

We appreciate the contributions of the Sanders-Brown Center on Aging at the University of Kentucky. We are deeply grateful to the research participants and their families who make this research possible. We thank Sonya L. Anderson from the University of Kentucky brain bank for preparing the brain samples used in this study. We would like to thank the University of Kentucky Center for Computational Sciences and Information Technology Services Research Computing for their support and use of the Morgan Compute Cluster and associated research computing resources. We would like to thank Singularity Sylabs for providing support and extra cloud storage for our software containers. We are grateful for the support from the Goeke lab members who quickly and thoroughly answered our numerous questions about bambu on GitHub. We would like to thank Dr. Thiago Wendt Viola, Dr. Rodrigo Grassi-Oliveira, and Dr. Consuelo Walss-Bass for guidance and help in the early stages of the proteomics analysis. We thank the reviewers for their sincere and meaningful contributions to improve the quality of the manuscript. The results published here are in in part based on data obtained from the AD Knowledge Portal. Short-read RNAseq data used for cross-validation of results in this study were provided by the Rush Alzheimer’s Disease Center, Rush University Medical Center, Chicago. Rush Alzheimer’s Disease Center data collection was supported through funding by NIA grants P30AG10161 (ROS), R01AG15819 (ROSMAP; genomics and RNAseq), R01AG17917 (MAP), RC2AG0365 (RNAseq).

## Extended Data

**Extended Data Figure 1:**
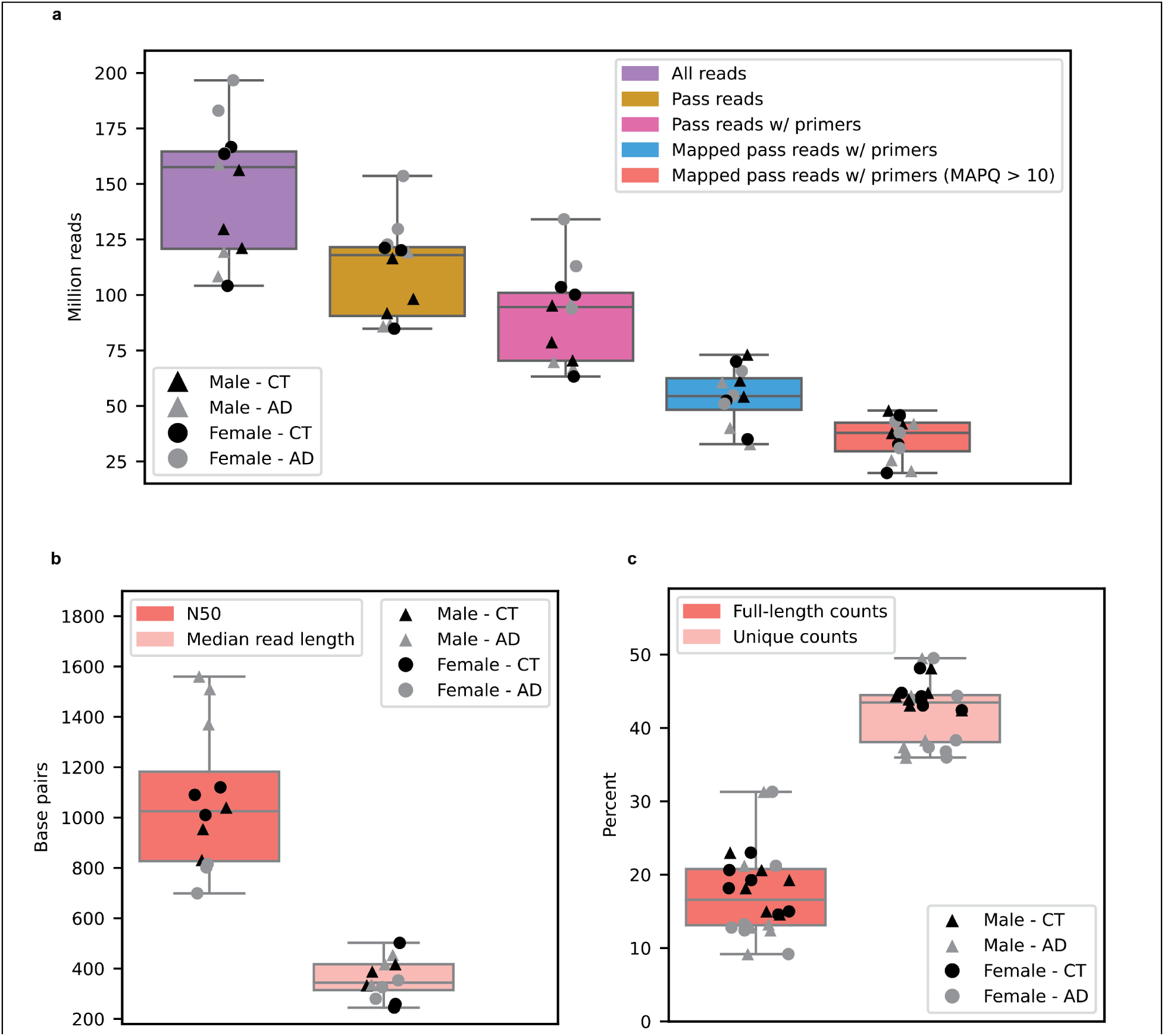
Basic sequencing metrics. **a,** Number of reads per sample after each step of the analysis. All downstream analysis were done with Mapped pass reads with both primers an MAPQ > 10. **b,** N50 and median read length for Mapped pass reads with both primers and MAPQ > 10. **c,** Percentage of reads that are full-length or unique as determined by bambu. Full-length counts = reads containing all exon-exon boundaries (i.e., intron chain) from its respective transcript. Unique counts = reads that were assigned to a single transcript.

**Extended Data Figure 2:**
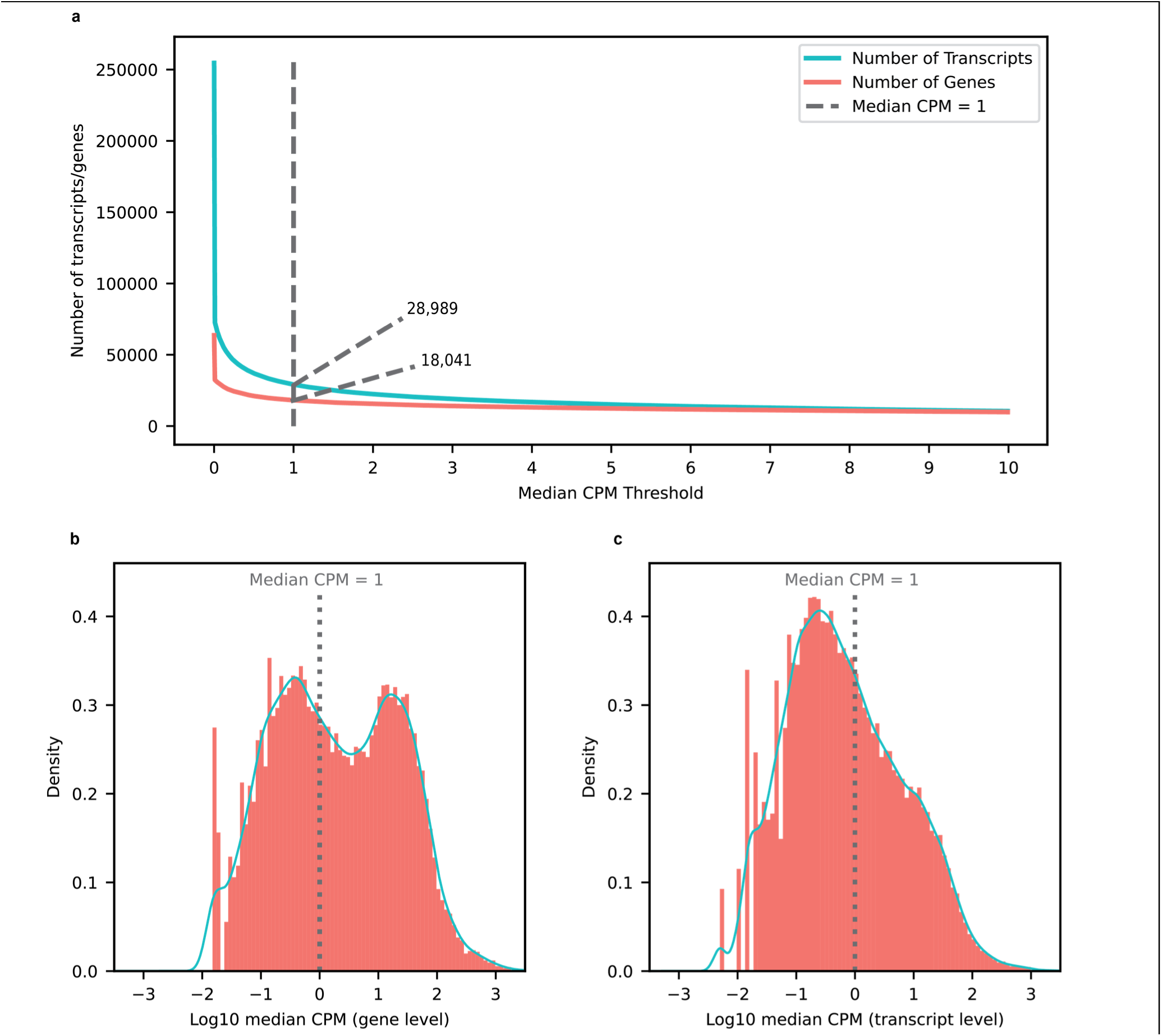
Expression distribution and diversity for genes and transcripts. **a,** Number of genes and transcripts represented across median CPM threshold. Cutoff shown as the dotted line set at median CPM = 1. **b,** Distribution of log10 median CPM values for gene bodies, dotted line shows cutoff point of median CPM = 1. **c,** Distribution of log10 median CPM values for gene bodies, dotted line shows cutoff point of median CPM = 1.

**Extended Data Figure 3:**
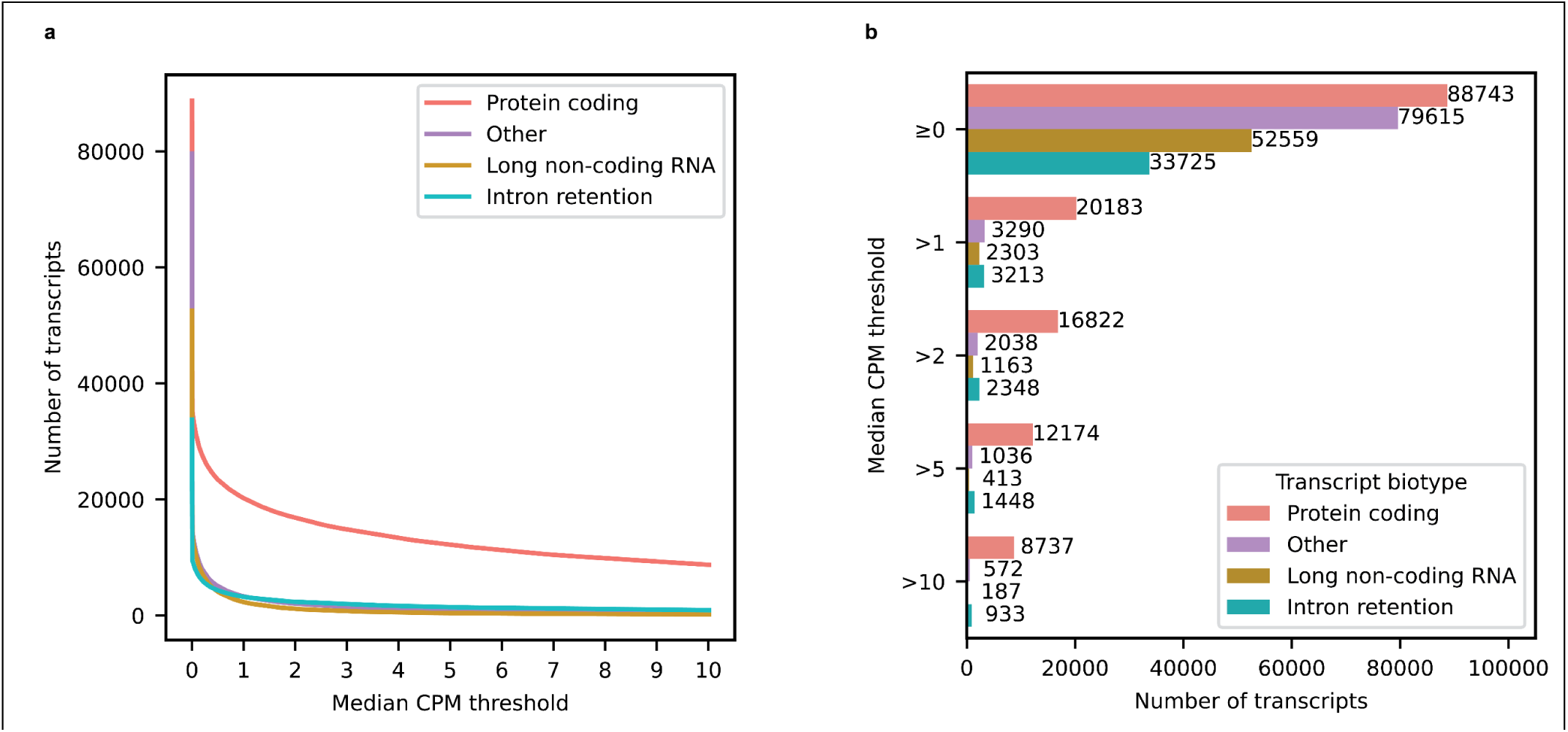
Expression of different transcript biotypes on aged human frontal cortex tissue using long-read RNAseq data. **a,** Lineplot showing the number of transcripts from different biotypes expressed above different median CPM threshold in long-read RNAseq data from aged human dorsolateral frontal cortex postmortem tissue. b, Barplot showing the number of transcripts from different biotypes expressed at or above different median CPM threshold in long-read RNAseq data from aged human dorsolateral frontal cortex postmortem tissue.

**Extended Data Figure 4:**
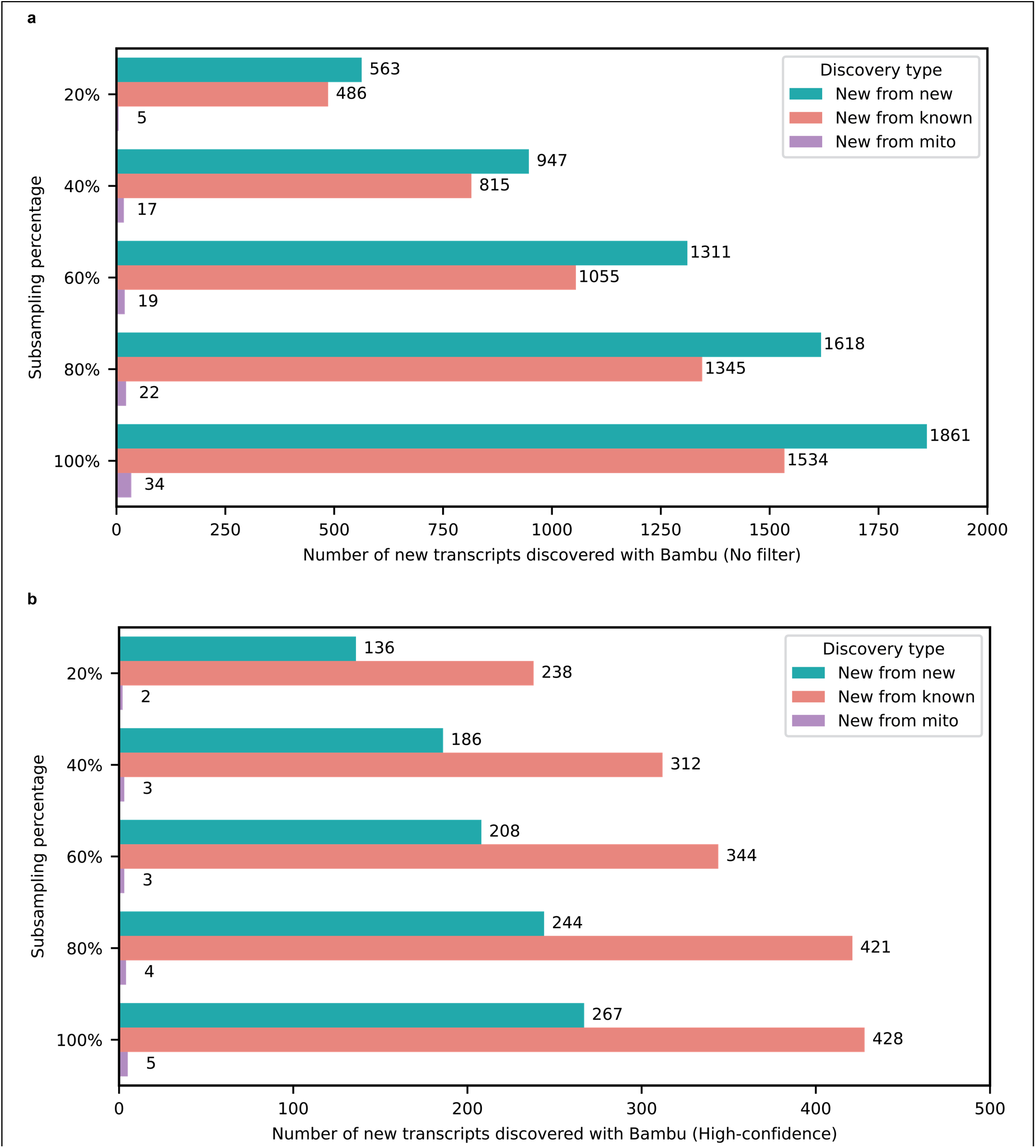
Number of newly discovered transcripts across subsampling range. **a,** Barplot showing the subsampling percentage on the Y-axis and number of new transcripts discovered with Bambu without filtering by expression estimates (no filter) on the X-axis. **b,** Barplot showing the subsampling percentage on the Y-axis and number of new transcripts discovered with Bambu when filtering by expression estimates X-axis (high-confidence; median CPM > 1). Nuclear encoded transcripts were filtered by median CPM > 1 and mitochondrially encoded transcripts were filtered by median full-length counts > 40. We used a different filter for mitochondrial transcripts due to issues in read assignment due to the polycistronic nature of mitochondrial transcription.

**Extended Data Figure 5:**
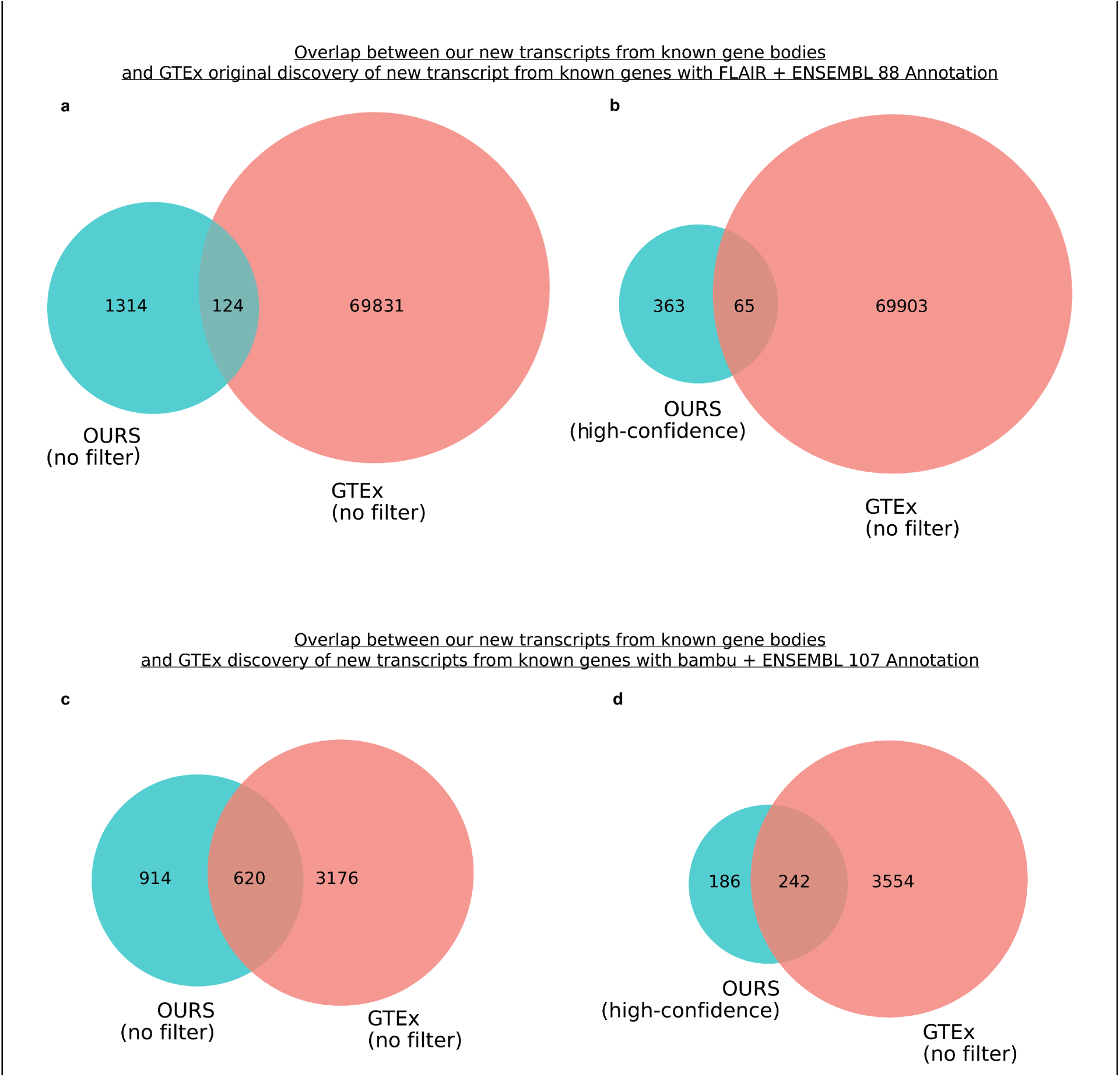
Difference in transcript discovery overlap based on annotation and computational tool used. **a,** Venn diagram showing the overlap between all our new transcripts from known gene bodies and new transcripts from known gene bodies in original GTEx long-read RNAseq article published by Glinos et al.^18^ using FLAIR for transcript discovery and ENSEMBL 88 annotation. **b,** Same as **a** but showing comparison only for new high-confidence transcripts from known gene bodies in our data. We used 70,000 as the number of new transcripts from known gene bodies in GTEx since they report just over 70,000 novel transcripts for annotated genes in their abstract. **c,** Venn diagram showing the overlap between all our new transcripts from known gene bodies and new transcripts from known gene bodies found when running GTEx long-read RNAseq data from article published by Glinos et al.^18^ using bambu for transcript discovery and ENSEMBL 107 annotation. **d,** Same as **a** but showing comparison only for new high-confidence transcripts from known gene bodies in our data. Venn diagrams are not to scale to improve readability.

**Extended Data Figure 6.**
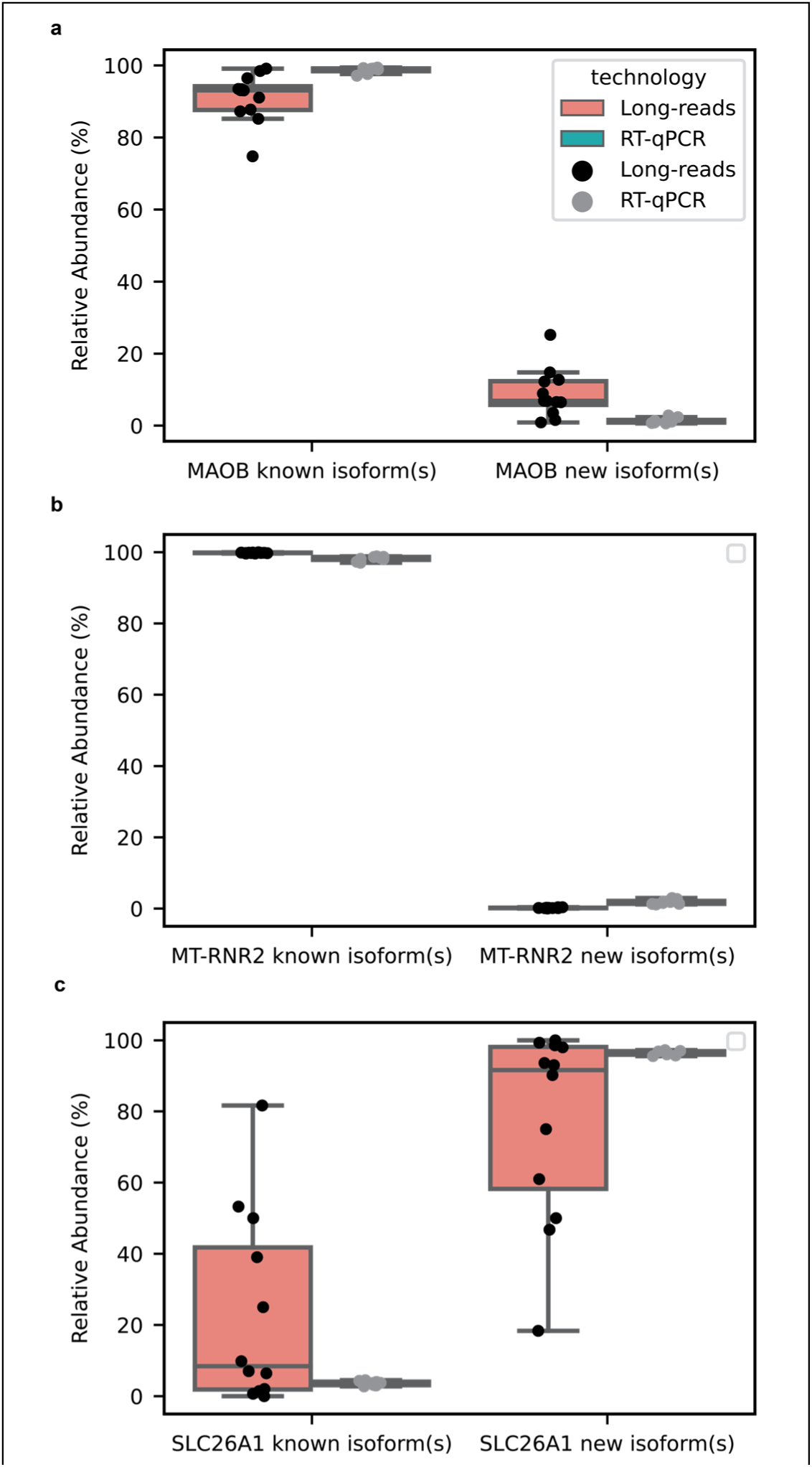
RT-qPCR validations for new RNA isoforms from MAOB, SLC26A1, *MT-RNR2* RNA isoforms match long-read sequencing data. **a,** Comparison of relative abundance between long-read sequencing and RT-qPCR for RNA isoforms in *MAOB*. **b,** Same as **a**, but for *MT-RNR2* c, Same as **a**, but for SLC26A1. Relative abundance was calculated as

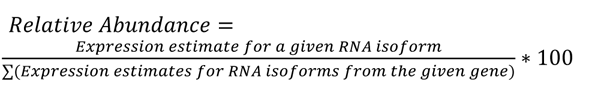 We used CPM (Counts Per Million) as the expression estimate for long-read sequencing and 2^(-ΔCt) for RT-qPCR. We used 2-ΔCt as the expression estimate instead of the more common 2-ΔΔCt. This is because the 2-ΔΔCt is optimized for comparisons between samples within the same gene/isoform, but does not work well for comparison between genes/isoforms. On the other hand, the 2-ΔCt expression estimate allows for comparison between different genes/isoforms. The housekeeping gene for RT-qPCR was CYC1.

**Extended Data Figure 7:**
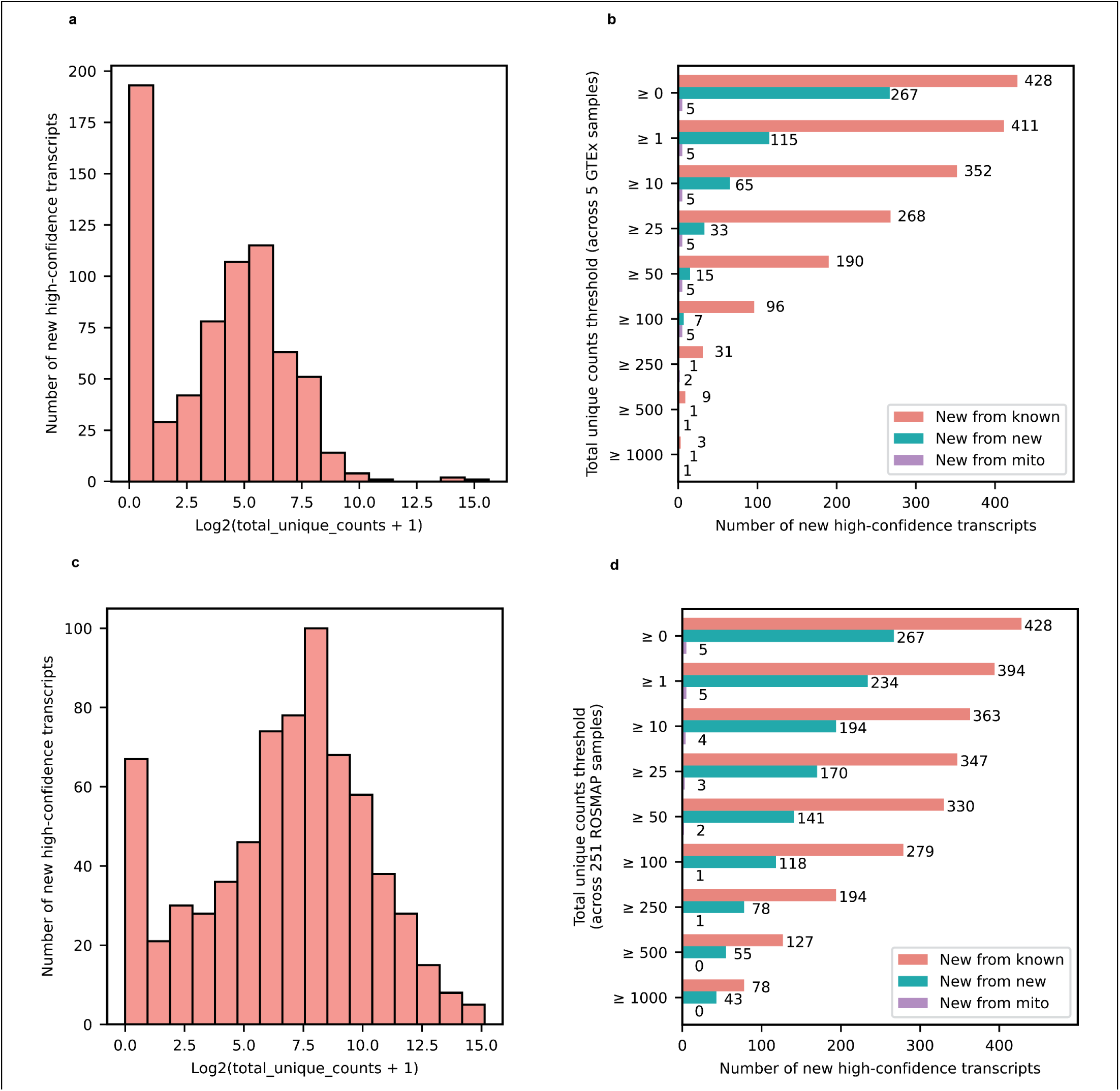
External validation of new high-confidence transcripts using publicly availabla data from 5 GTEx brain samples (Brodmann area 9) sequenced with long-read RNAseq and 251 ROSMAP brain samples (Brodmann area 9/46) sequenced with Illumina 150bp paired-end RNAseq reads. **a,** Histogram showing total unique counts for new high-onfidence transcripts across five GTEx long-read RNAseq data from brain samples. Total unique counts are shown in a log2(total unique counts + 1) scale to avoid streching generated by outliers. **b,** Barplot showing the number of new high-confidence transcripts that meet different total unique counts thresholds in cross-validation using five GTEx long-read RNAseq data from brain samples. The “≥ 0” Y-axis label shows the total number of high-confidence transcripts before any filtering. Legend colors: New from known denotes new transcripts from known gene bodies, New from new denotes new transcripts from newly discovered gene bodies, and new from mito denotes new mitochondrially encoded spliced transcripts. **c,** Same as a but for 251 ROSMAP brain samples sequenced with 150bp paired-end Illumina RNAseq. **d,** Same as **b** but for 251 ROSMAP brain samples sequenced with 150bp paired-end Illumina RNAseq. We observed that 98.8% of the new high-confidence transcripts from known gene bodies had at least one uniquely mapped read in either GTEx or ROSMAP data and 69.6% had at least 100 uniquely mapped reads in either dataset.

**Extended Data Figure 8:**
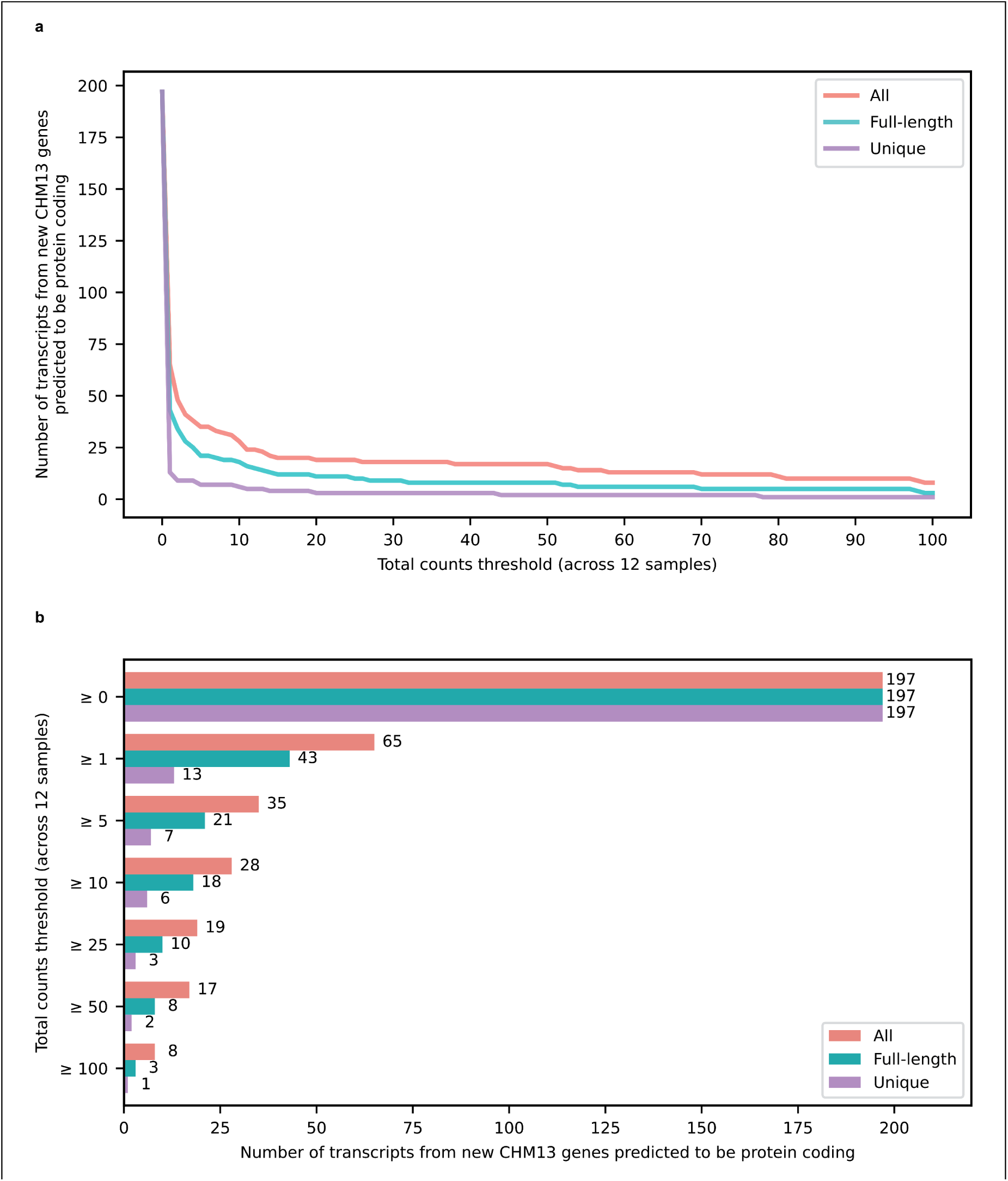
Expression of 197 transcripts from extra 99 predicted protein coding genes in CHM13 reported by Nurk et al. **a,** Lineplot with number of transcripts from extra 99 protein coding genes that are expressed across the total counts threshold for our 12 brain samples. The red line indicates all counts (including partial assignments), mint green line indicates full-length reads and purple line indicates unique reads. **b,** Barplot showing the number of transcripts from extra 99 protein coding genes expressed at or above different total counts thresholds. The top y-axis label shows all the 197 annotated RNA isoforms from the extra 99 predicted protein coding genes in CHM13 reported by Nurk et al.

**Extended Data Figure 9:**
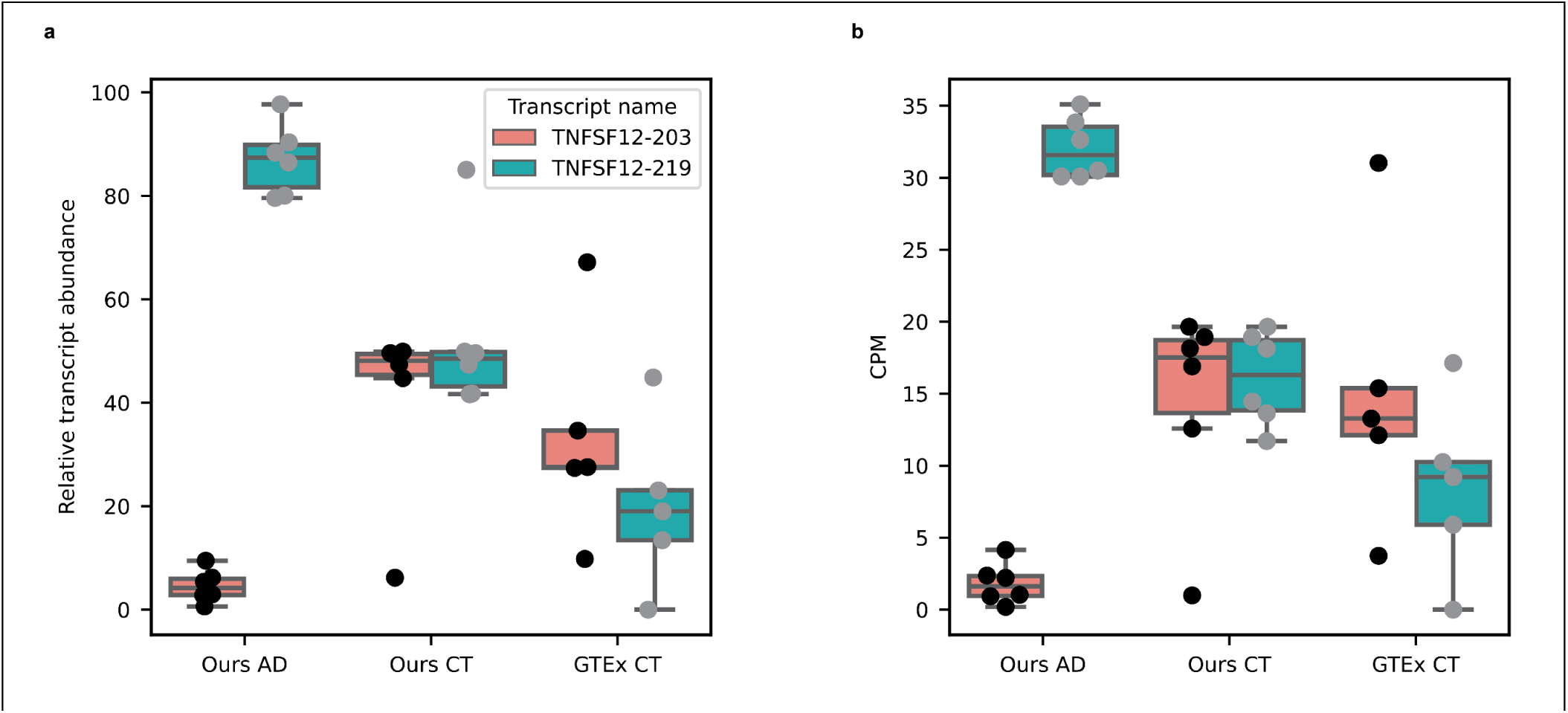
Attempt at validation of TNFSF12 RNA isoform expression pattern in healthy controls. **a,** Boxplot showing the relative transcript abudance (percentage) for TNFSF12 RNA isoforms that are differentially expressed between Alzheimer’s disease cases and controls in this study. On the X-axis, the “OURS AD” and “OURS CT” labels represents the six Alzheimer’s disease and six control brain samples sequenced in this study. The “GTEx CT” label represents the 5 GTEx brain samples (Brodmann area 9) sequences with PCR amplified long-read nanopore RNAseq. **b,** Boxplot showing the CPM for TNFSF12 RNA isoforms that are differentially expressed between Alzheimer’s disease cases and controls in this study. X-axis labels follow the same pattern as **a.**

**Extended Data Figure 10:**
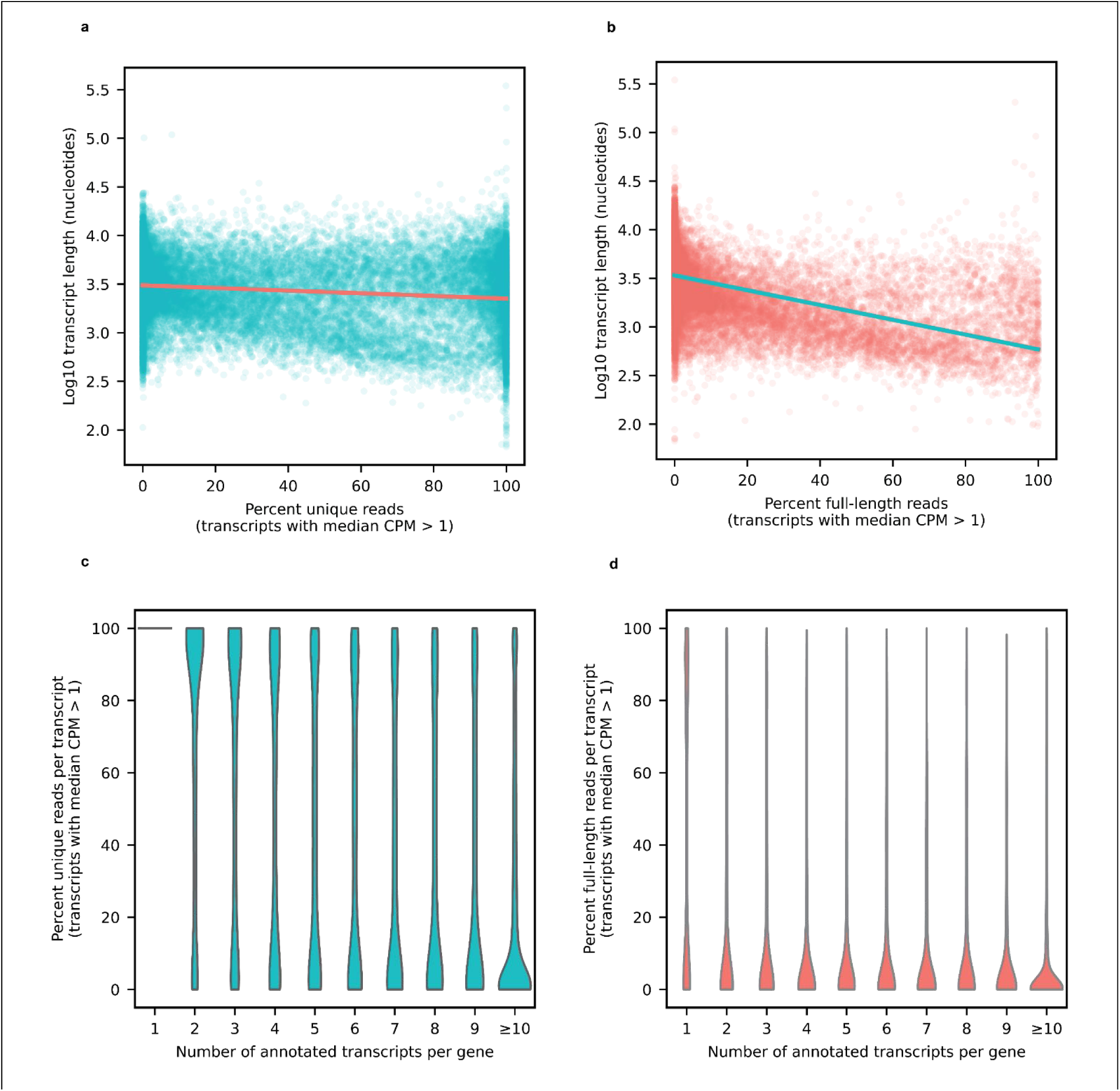
Percentage of unique and full-length reads per transcript. **a,** Scatterplot showing the percentage of uniquely aligned reads for each transcript with a median CPM > 1 on the X-axis and the Log10 transcript length on the Y axis. **b,** Scatterplot showing the percentage of full-length reads for each transcript with a median CPM > 1 on the X-axis and the Log10 transcript length on the Y axis. **c**, Violin plot showing the percentage of uniquely aligned reads for each transcript with median CPM > 1 on the Y-axis and the number of annotated transcript per gene on the X-axis. **d,** Violin plot showing the percentage of full-length reads for each transcript with median CPM > 1 on the Y-axis and the number of annotated transcript per gene on the X-axis.

## Notes

https://ebbertlab.com/brain_rna_isoform_seq.html

https://zenodo.org/records/8180677

https://www.synapse.org/#!Synapse:syn52047893/wiki/622953

### Competing Interest Statement

The authors have declared no competing interest.

### Summary of Updates

This manuscript has been revised to include more validation of unnanotaed/new RNA isoforms. These validations include using external datasets (in-silico) as well as orthogonal methods such as RT-qPCR. Manuscript has also been revised to present a more thorough description of limitations.

## References

1. Park, E., Pan, Z., Zhang, Z., Lin, L. & Xing, Y. The Expanding Landscape of Alternative Splicing Variation in Human Populations. Am. J. Hum. Genet. 102, 11–26 (2018).

2. Martin, F. J. et al. Ensembl 2023. Nucleic Acids Res. 51, D933–D941 (2023).

3. Yang, X. et al. Widespread Expansion of Protein Interaction Capabilities by Alternative Splicing. Cell 164, 805–817 (2016).

4. Oberwinkler, J., Lis, A., Giehl, K. M., Flockerzi, V. & Philipp, S. E. Alternative splicing switches the divalent cation selectivity of TRPM3 channels. J. Biol. Chem. 280, 22540–22548 (2005).

5. Végran, F. et al. Overexpression of caspase-3s splice variant in locally advanced breast carcinoma is associated with poor response to neoadjuvant chemotherapy. Clin. Cancer Res. Off. J. Am. Assoc. Cancer Res. 12, 5794–5800 (2006).

6. Warren, C. F. A., Wong-Brown, M. W. & Bowden, N. A. BCL-2 family isoforms in apoptosis and cancer. Cell Death Dis. 10, 177 (2019).

7. Trapnell, C. et al. Transcript assembly and quantification by RNA-Seq reveals unannotated transcripts and isoform switching during cell differentiation. Nat. Biotechnol. 28, 511–515 (2010).

8. Tilgner, H. et al. Comprehensive transcriptome analysis using synthetic long-read sequencing reveals molecular co-association of distant splicing events. Nat. Biotechnol. 33, 736–742 (2015).

9. Ringeling, F. R. et al. Partitioning RNAs by length improves transcriptome reconstruction from short-read RNA-seq data. Nat. Biotechnol. 40, 741–750 (2022).

10. Benchmarking long-read RNA-sequencing analysis tools using in silico mixtures | Nature Methods. https://www.nature.com/articles/s41592-023-02026-3.

11. Chen, Y. et al. Context-aware transcript quantification from long-read RNA-seq data with Bambu. Nat. Methods 1–9 (2023) doi:10.1038/s41592-023-01908-w.

12. Dou, Z. et al. Aberrant Bcl-x splicing in cancer: from molecular mechanism to therapeutic modulation. J. Exp. Clin. Cancer Res. 40, 194 (2021).

13. Tilgner, H. et al. Microfluidic isoform sequencing shows widespread splicing coordination in the human transcriptome. Genome Res. 28, 231–242 (2018).

14. Course, M. M. et al. Aberrant splicing of PSEN2, but not PSEN1, in individuals with sporadic Alzheimer’s disease. Brain J. Neurol. 146, 507–518 (2023).

15. Okubo, M. et al. RNA-seq analysis, targeted long-read sequencing and in silico prediction to unravel pathogenic intronic events and complicated splicing abnormalities in dystrophinopathy. Hum. Genet. 142, 59–71 (2023).

16. Liu, M. et al. Long-read sequencing reveals oncogenic mechanism of HPV-human fusion transcripts in cervical cancer. Transl. Res. J. Lab. Clin. Med. 253, 80–94 (2023).

17. Schwenk, V. et al. Transcript capture and ultradeep long-read RNA sequencing (CAPLRseq) to diagnose HNPCC/Lynch syndrome. J. Med. Genet. 60, 747–759 (2023).

18. Glinos, D. A. et al. Transcriptome variation in human tissues revealed by long-read sequencing. Nature 608, 353– 359 (2022).

19. Leung, S. K. et al. Full-length transcript sequencing of human and mouse cerebral cortex identifies widespread isoform diversity and alternative splicing. Cell Rep. 37, 110022 (2021).

20. Mattick, J. S. et al. Long non-coding RNAs: definitions, functions, challenges and recommendations. Nat. Rev. Mol. Cell Biol. 24, 430–447 (2023).

21. Mostafavi, S. et al. A molecular network of the aging human brain provides insights into the pathology and cognitive decline of Alzheimer’s disease. Nat. Neurosci. 21, 811–819 (2018).

22. Systematic assessment of long-read RNA-seq methods for transcript identification and quantification - PubMed. https://pubmed.ncbi.nlm.nih.gov/37546854/.

23. Tang, A. D. et al. Full-length transcript characterization of SF3B1 mutation in chronic lymphocytic leukemia reveals downregulation of retained introns. Nat. Commun. 11, 1438 (2020).

24. Tseng, E. cDNA_Cupcake. (2023).

25. Bustin, S. A. et al. The MIQE Guidelines: Minimum Information for Publication of Quantitative Real-Time PCR Experiments. Clin. Chem. 55, 611–622 (2009).

26. Nurk, S. et al. The complete sequence of a human genome. Science 376, 44–53 (2022).

27. Wagner, J. et al. Curated variation benchmarks for challenging medically relevant autosomal genes. Nat. Biotechnol. 40, 672–680 (2022).

28. Singh, T. et al. Rare coding variants in ten genes confer substantial risk for schizophrenia. Nature 604, 509–516 (2022).

29. Palmer, D. S. et al. Exome sequencing in bipolar disorder identifies AKAP11 as a risk gene shared with schizophrenia. Nat. Genet. 54, 541–547 (2022).

30. Billingsley, K. J., Bandres-Ciga, S., Saez-Atienzar, S. & Singleton, A. B. Genetic risk factors in Parkinson’s disease. Cell Tissue Res. 373, 9–20 (2018).

31. Perrone, F., Cacace, R., van der Zee, J. & Van Broeckhoven, C. Emerging genetic complexity and rare genetic variants in neurodegenerative brain diseases. Genome Med. 13, 59 (2021).

32. Shadrina, M., Bondarenko, E. A. & Slominsky, P. A. Genetics Factors in Major Depression Disease. Front. Psychiatry 9, 334 (2018).

33. Satterstrom, F. K. et al. Large-Scale Exome Sequencing Study Implicates Both Developmental and Functional Changes in the Neurobiology of Autism. Cell 180, 568–584.e23 (2020).

34. Stein, M. B. et al. Genome-wide association analyses of post-traumatic stress disorder and its symptom subdomains in the Million Veteran Program. Nat. Genet. 53, 174–184 (2021).

35. Maihofer, A. X. et al. Enhancing Discovery of Genetic Variants for Posttraumatic Stress Disorder Through Integration of Quantitative Phenotypes and Trauma Exposure Information. Biol. Psychiatry 91, 626–636 (2022).

36. Hatoum, A. S. et al. Multivariate genome-wide association meta-analysis of over 1 million subjects identifies loci underlying multiple substance use disorders. Nat. Ment. Health 1, 210–223 (2023).

37. Bellenguez, C. et al. New insights into the genetic etiology of Alzheimer’s disease and related dementias. Nat. Genet. 54, 412–436 (2022).

38. Gee, H. Y. et al. Mutations in SLC26A1 Cause Nephrolithiasis. Am. J. Hum. Genet. 98, 1228–1234 (2016).

39. Pfau, A. et al. SLC26A1 is a major determinant of sulfate homeostasis in humans. J. Clin. Invest. 133, e161849 (2023).

40. Parvari, R. et al. A recessive contiguous gene deletion of chromosome 2p16 associated with cystinuria and a mitochondrial disease. Am. J. Hum. Genet. 69, 869–875 (2001).

41. Shaheen, R. et al. Mutation in WDR4 impairs tRNA m(7)G46 methylation and causes a distinct form of microcephalic primordial dwarfism. Genome Biol. 16, 210 (2015).

42. Braun, D. A. et al. Mutations in WDR4 as a new cause of Galloway-Mowat syndrome. Am. J. Med. Genet. A. 176, 2460–2465 (2018).

43. Osborn, D. P. S. et al. Autosomal recessive cardiomyopathy and sudden cardiac death associated with variants in MYL3. Genet. Med. 23, 787–792 (2021).

44. Gilbody, S., Lewis, S. & Lightfoot, T. Methylenetetrahydrofolate reductase (MTHFR) genetic polymorphisms and psychiatric disorders: a HuGE review. Am. J. Epidemiol. 165, 1–13 (2007).

45. Lee, H. J. et al. Association study of polymorphisms in synaptic vesicle-associated genes, SYN2 and CPLX2, with schizophrenia. *Behav. Brain Funct.* **1**, 15 (2005).

46. Tan, Y.-Y., Jenner, P. & Chen, S.-D. Monoamine Oxidase-B Inhibitors for the Treatment of Parkinson’s Disease: Past, Present, and Future. J. Park. Dis. 12, 477–493 (2022).

47. Guerreiro, R. et al. TREM2 Variants in Alzheimer’s Disease. N. Engl. J. Med. 368, 117–127 (2013).

48. Kiianitsa, K. et al. Novel TREM2 splicing isoform that lacks the V-set immunoglobulin domain is abundant in the human brain. J. Leukoc. Biol. 110, 829–837 (2021).

49. Shaw, B. C. et al. An alternatively spliced TREM2 isoform lacking the ligand binding domain is expressed in human brain. J. Alzheimers Dis. JAD 87, 1647–1657 (2022).

50. Tsegay, P. S. et al. Incorporation of 5’,8-cyclo-2’deoxyadenosines by DNA repair polymerases via base excision repair. *DNA Repair* **109**, 103258 (2022).

51. Kaufman, B. A. & Van Houten, B. POLB: A new role of DNA polymerase beta in mitochondrial base excision repair. DNA Repair 60, A1–A5 (2017).

52. Butchbach, M. E. R. Genomic Variability in the Survival Motor Neuron Genes (SMN1 and SMN2): Implications for Spinal Muscular Atrophy Phenotype and Therapeutics Development. Int. J. Mol. Sci. 22, 7896 (2021).

53. Melnik, B. C. & Plewig, G. Impaired Notch signalling: the unifying mechanism explaining the pathogenesis of hidradenitis suppurativa (acne inversa). Br. J. Dermatol. 168, 876–878 (2013).

54. Dimmock, D. P. et al. Clinical and molecular features of mitochondrial DNA depletion due to mutations in deoxyguanosine kinase. Hum. Mutat. 29, 330–331 (2008).

55. Guo, B. et al. Humanin peptide suppresses apoptosis by interfering with Bax activation. Nature 423, 456–461 (2003).

56. Herai, R. H., Negraes, P. D. & Muotri, A. R. Evidence of nuclei-encoded spliceosome mediating splicing of mitochondrial RNA. Hum. Mol. Genet. 26, 2472–2479 (2017).

57. Rahman, S. Mitochondrial disease and epilepsy. Dev. Med. Child Neurol. 54, 397–406 (2012).

58. Delatycki, M. B. & Bidichandani, S. I. Friedreich ataxia- pathogenesis and implications for therapies. Neurobiol. Dis. 132, 104606 (2019).

59. Lin, M. T. & Beal, M. F. Mitochondrial dysfunction and oxidative stress in neurodegenerative diseases. Nature 443, 787–795 (2006).

60. Amorim, J. A. et al. Mitochondrial and metabolic dysfunction in ageing and age-related diseases. Nat. Rev. Endocrinol. 18, 243–258 (2022).

61. Sen, P. et al. Spurious intragenic transcription is a feature of mammalian cellular senescence and tissue aging. *Nat*. Aging 3, 402–417 (2023).

62. VanDongen, A. M. J. & VanDongen, H. M. A. Effects of mRNA untranslated regions on translational efficiency of NMDA receptor subunits. Neurosignals 13, 194–206 (2004).

63. Leppek, K., Das, R. & Barna, M. Functional 5′ UTR mRNA structures in eukaryotic translation regulation and how to find them. Nat. Rev. Mol. Cell Biol. 19, 158–174 (2018).

64. Goedert, M., Wischik, C. M., Crowther, R. A., Walker, J. E. & Klug, A. Cloning and sequencing of the cDNA encoding a core protein of the paired helical filament of Alzheimer disease: identification as the microtubule-associated protein tau. Proc. Natl. Acad. Sci. U. S. A. 85, 4051–4055 (1988).

65. Goedert, M., Spillantini, M. G., Potier, M. C., Ulrich, J. & Crowther, R. A. Cloning and sequencing of the cDNA encoding an isoform of microtubule-associated protein tau containing four tandem repeats: differential expression of tau protein mRNAs in human brain. EMBO J. 8, 393–399 (1989).

66. Andreadis, A., Brown, W. M. & Kosik, K. S. Structure and novel exons of the human tau gene. Biochemistry 31, 10626–10633 (1992).

67. Ratajczak, W., Atkinson, S. D. & Kelly, C. The TWEAK/Fn14/CD163 axis-implications for metabolic disease. Rev. Endocr. Metab. Disord. 23, 449–462 (2022).

68. Boström, G. et al. Different Inflammatory Signatures in Alzheimer’s Disease and Frontotemporal Dementia Cerebrospinal Fluid. J. Alzheimers Dis. 81, 629–640.

69. Masters, C. L. et al. Amyloid plaque core protein in Alzheimer disease and Down syndrome. Proc. Natl. Acad. Sci. U. S. A. 82, 4245–4249 (1985).

70. Loureiro, L. O. et al. A recurrent SHANK3 frameshift variant in Autism Spectrum Disorder. Npj Genomic Med. 6, 1–12 (2021).

71. Uchino, S. & Waga, C. SHANK3 as an autism spectrum disorder-associated gene. Brain Dev. 35, 106–110 (2013).

72. Kelemen, O. et al. Function of alternative splicing. Gene 514, 1–30 (2013).

73. Keren, H., Lev-Maor, G. & Ast, G. Alternative splicing and evolution: diversification, exon definition and function. Nat. Rev. Genet. 11, 345–355 (2010).

74. Kim, E., Magen, A. & Ast, G. Different levels of alternative splicing among eukaryotes. Nucleic Acids Res. 35, 125– 131 (2007).

75. Schmitt, F. A., et al. University of Kentucky Sanders-Brown Healthy Brain Aging Volunteers: Donor Characteristics, Procedures and Neuropathology. Curr. Alzheimer Res. 9, 724–733 (2012).

76. epi2me-labs/pychopper: cDNA read preprocessing. https://github.com/epi2me-labs/pychopper.

77. Li, H. Minimap2: pairwise alignment for nucleotide sequences. Bioinformatics 34, 3094–3100 (2018).

78. Li, H. et al. The Sequence Alignment/Map format and SAMtools. Bioinforma. Oxf. Engl. 25, 2078–2079 (2009).

79. Leger, A. & Leonardi, T. pycoQC, interactive quality control for Oxford Nanopore Sequencing. J. Open Source Softw. 4, 1236 (2019).

80. Dobin, A. et al. STAR: ultrafast universal RNA-seq aligner. Bioinformatics 29, 15–21 (2013).

81. Patro, R., Duggal, G., Love, M. I., Irizarry, R. A. & Kingsford, C. Salmon: fast and bias-aware quantification of transcript expression using dual-phase inference. Nat. Methods 14, 417–419 (2017).

82. Bailey, T. L. et al. MEME SUITE: tools for motif discovery and searching. Nucleic Acids Res. 37, W202–208 (2009).

83. ROCA, X., SACHIDANANDAM, R. & KRAINER, A. R. Determinants of the inherent strength of human 5′ splice sites. RNA 11, 683–698 (2005).

84. Carranza, F., Shenasa, H. & Hertel, K. J. Splice site proximity influences alternative exon definition. RNA Biol. 19, 829–840.

85. Pertea, G. & Pertea, M. GFF Utilities: GffRead and GffCompare. F1000Research 9, (2020).

86. Love, M. I., Huber, W. & Anders, S. Moderated estimation of fold change and dispersion for RNA-seq data with DESeq2. Genome Biol. 15, 550 (2014).

87. Gustavsson, E. K., Zhang, D., Reynolds, R. H., Garcia-Ruiz, S. & Ryten, M. ggtranscript: an R package for the visualization and interpretation of transcript isoforms using ggplot2. Bioinformatics 38, 3844–3846 (2022).

88. Penna, I. et al. Selection of Candidate Housekeeping Genes for Normalization in Human Postmortem Brain Samples. Int. J. Mol. Sci. 12, 5461–5470 (2011).

89. Johnson, E. C. B. et al. Large-scale deep multi-layer analysis of Alzheimer’s disease brain reveals strong proteomic disease-related changes not observed at the RNA level. Nat. Neurosci. 25, 213–225 (2022).

90. Higginbotham, L. et al. Unbiased Classification of the Human Brain Proteome Resolves Distinct Clinical and Pathophysiological Subtypes of Cognitive Impairment. 2022.07.22.501017 Preprint at 10.1101/2022.07.22.501017 (2022).

91. Sinitcyn, P. et al. Global detection of human variants and isoforms by deep proteome sequencing. Nat. Biotechnol. (2023) doi:10.1038/s41587-023-01714-x.

92. ProteoGenomics Analysis Toolkit. (2023).

93. Nesvilab/FragPipe. (2023).

94. Chang, H.-Y. et al. Crystal-C: A computational tool for refinement of open search results. J. Proteome Res. 19, 2511–2515 (2020).

95. Kong, A. T., Leprevost, F. V., Avtonomov, D. M., Mellacheruvu, D. & Nesvizhskii, A. I. MSFragger: ultrafast and comprehensive peptide identification in mass spectrometry–based proteomics. Nat. Methods 14, 513–520 (2017).

96. da Veiga Leprevost, F., et al. Philosopher: a versatile toolkit for shotgun proteomics data analysis. Nat. Methods 17, 869–870 (2020).

97. Yu, F., Haynes, S. E. & Nesvizhskii, A. I. IonQuant Enables Accurate and Sensitive Label-Free Quantification With FDR-Controlled Match-Between-Runs. Mol. Cell. Proteomics MCP 20, 100077 (2021).

98. Teo, G. C., Polasky, D. A., Yu, F. & Nesvizhskii, A. I. Fast Deisotoping Algorithm and Its Implementation in the MSFragger Search Engine. J. Proteome Res. 20, 498–505 (2021).

99. Tsou, C.-C. et al. DIA-Umpire: comprehensive computational framework for data-independent acquisition proteomics. Nat. Methods 12, 258–264, 7 p following 264 (2015).

100. Steinegger, M. & Söding, J. MMseqs2 enables sensitive protein sequence searching for the analysis of massive data sets. Nat. Biotechnol. 35, 1026–1028 (2017).

101. Nextflow enables reproducible computational workflows | Nature Biotechnology. https://www.nature.com/articles/nbt.3820.

102. Kurtzer, G. M., Sochat, V. & Bauer, M. W. Singularity: Scientific containers for mobility of compute. PLoS ONE 12, e0177459 (2017).

