## Supplementary Fig. for "Using deep long-read RNAseq in Alzheimer’s disease brain to assess medical relevance of RNA isoform diversity"

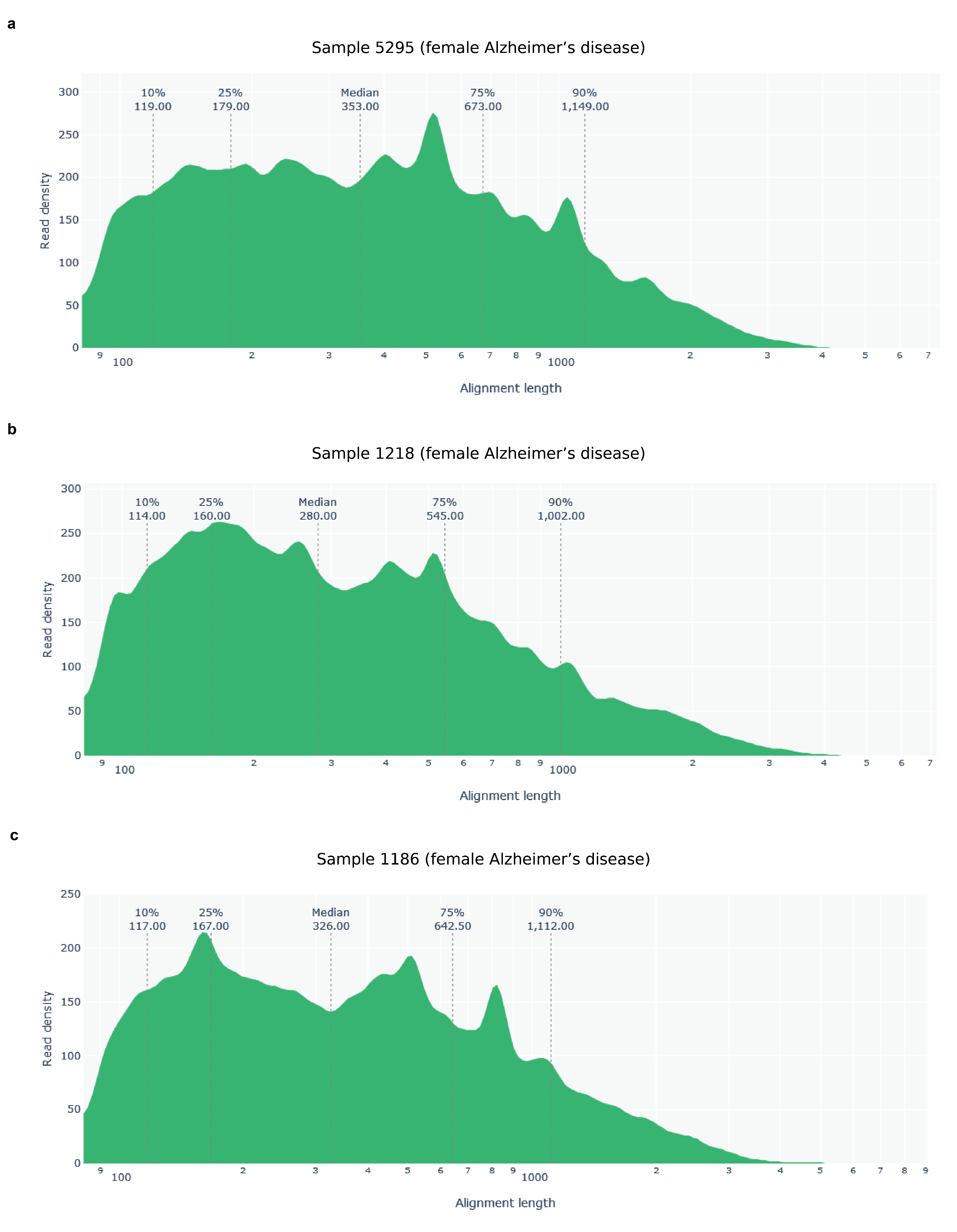

**Supplementary Figure 1: Aligned read length distribution for the three female control brain samples.**

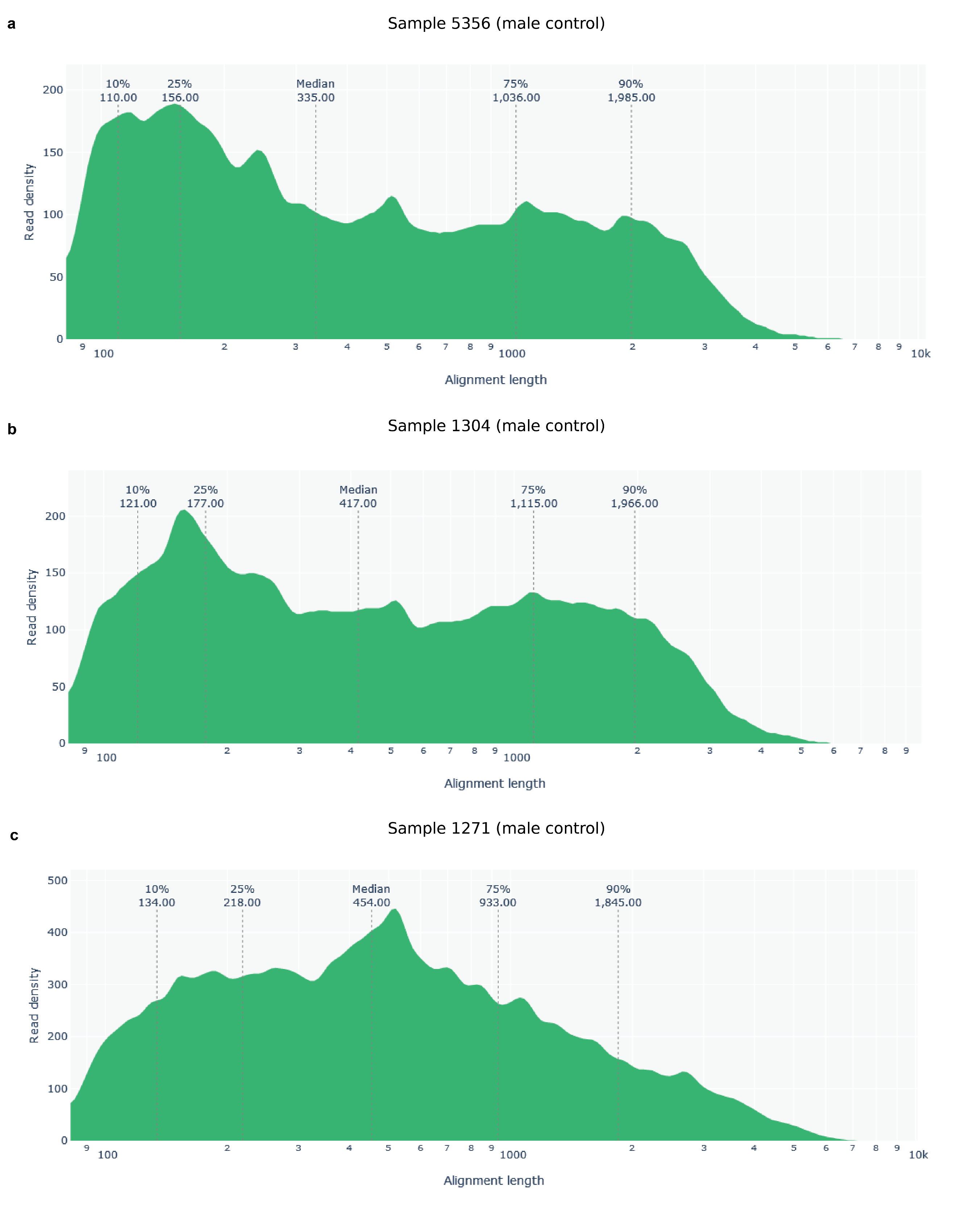

**Supplementary Figure 2: Aligned read length distribution for the three male control brain samples.**

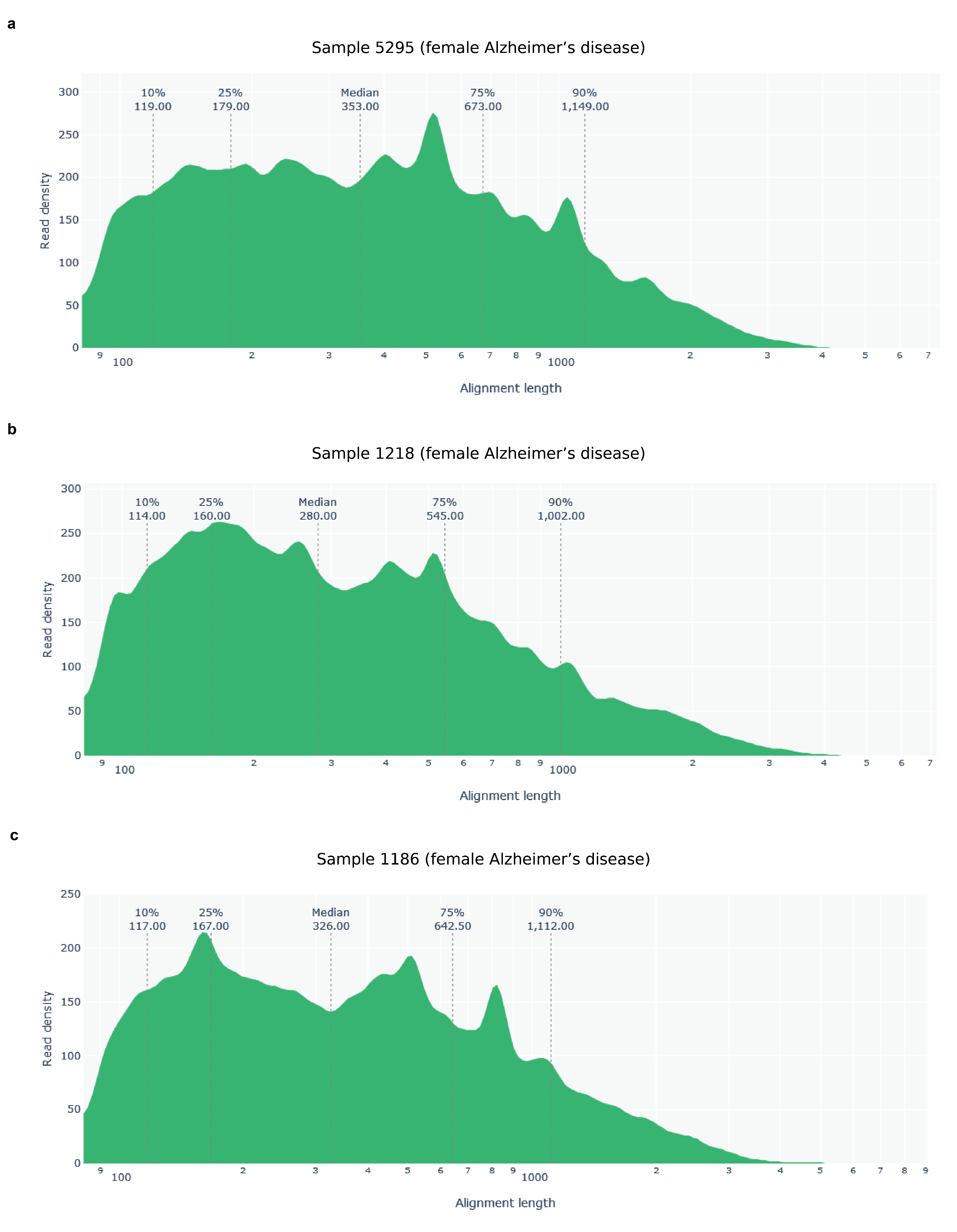

**Supplementary Figure 3: Aligned read length distribution for the three female Alzheimer’s disease brain samples.**

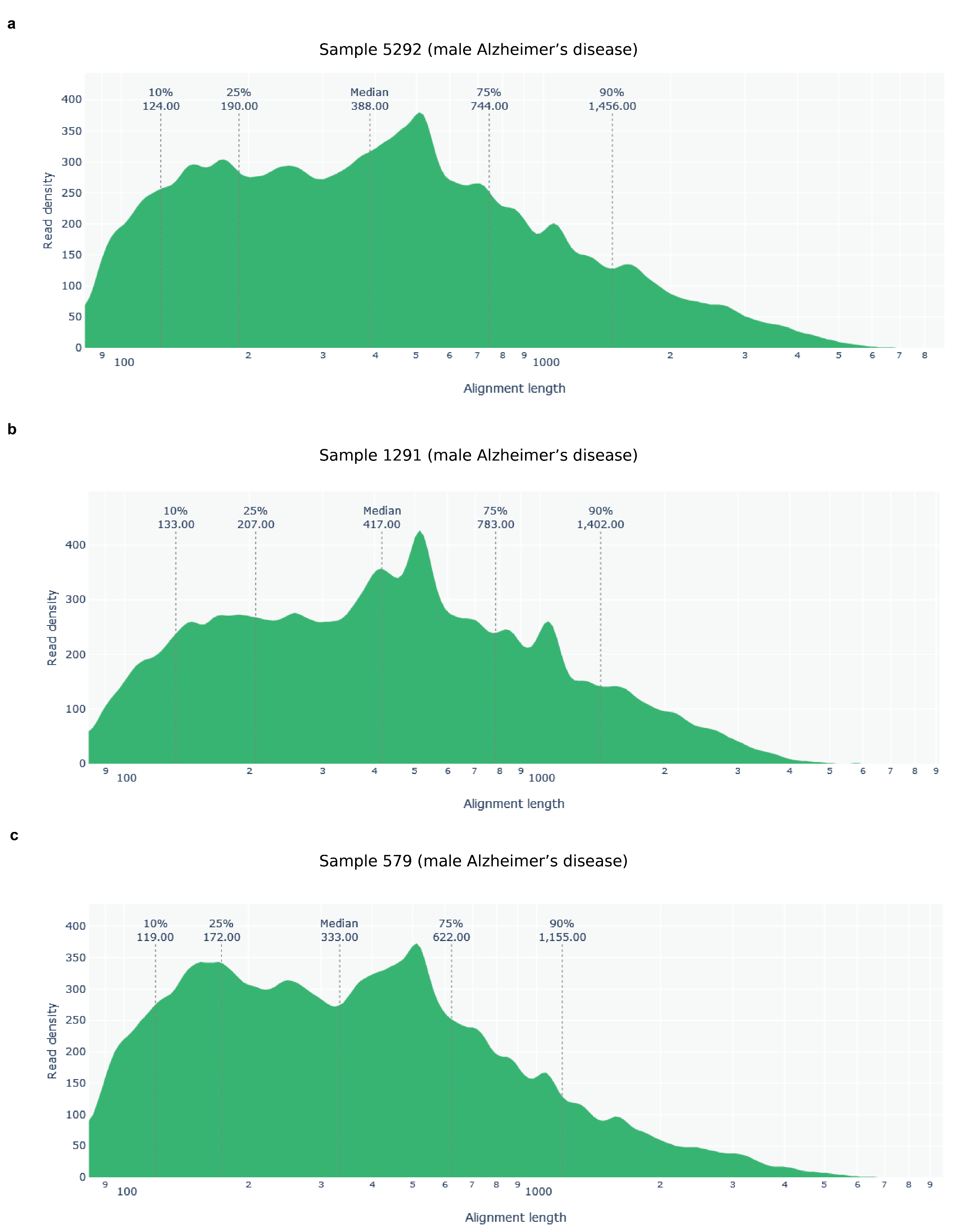

**Supplementary Figure 4: Aligned read length distribution for the three male Alzheimer’s disease brain samples.**

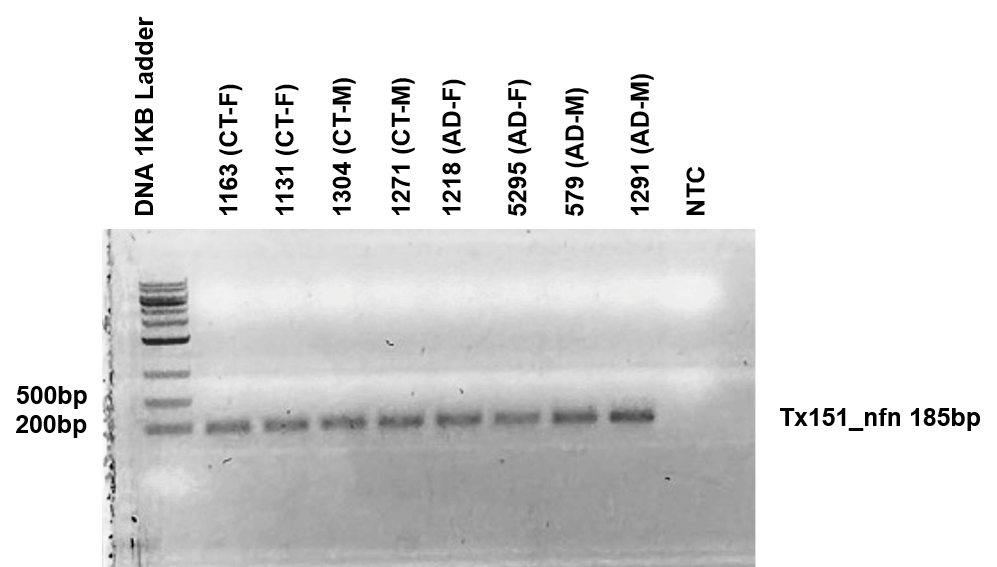

**Supplementary Figure 5: PCR validation for new high confidence transcript Tx151 (New gene body*).***

Number above the lanes is the sample id for 8 out of the 12 samples used in this study. The other 4 samples were not included because we ran out of tissue left for generating new cDNA. Labels on the right side of the figure indicate transcript id for the new high-confidence RNA isoform being validated and the expected product length from the PCR primers used to amplify the RNA isoform. Tx151 is a new RNA isoform from a newly discovered gene body and was succesfully validated in this gel.

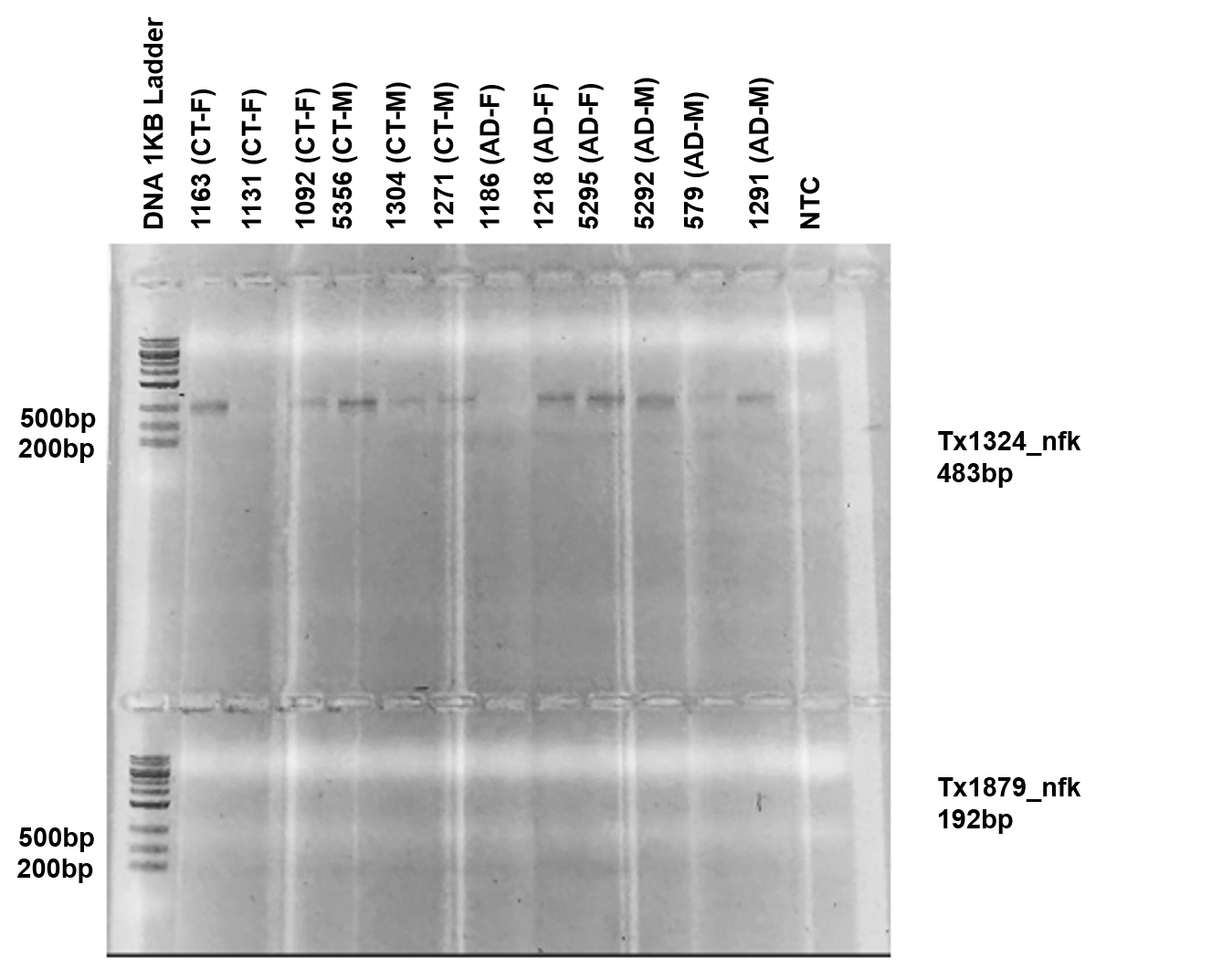

**Supplementary Figure 6: PCR validation for new high confidence transcripts Tx1324 (*NELFA)* and Tx1879 (*MAOB*).**

Number above the lanes is the sample id for 8 out of the 12 samples used in this study. The other 4 samples were not included because we ran out of tissue left for generating new cDNA. Labels on the right side of the figure indicate transcript id for the new high-confidence RNA isoform being validated and the expected product length from the PCR primers used to amplify the RNA isoform. Tx1324 is a new RNA isoform from the *NELFA* gene and was succesfully validated in this gel. Tx1879 is a new RNA isoform for the *MAOB* gene and was not validated in this gel, but was validated with a different primer pair in **Supplementary Figure 16**.

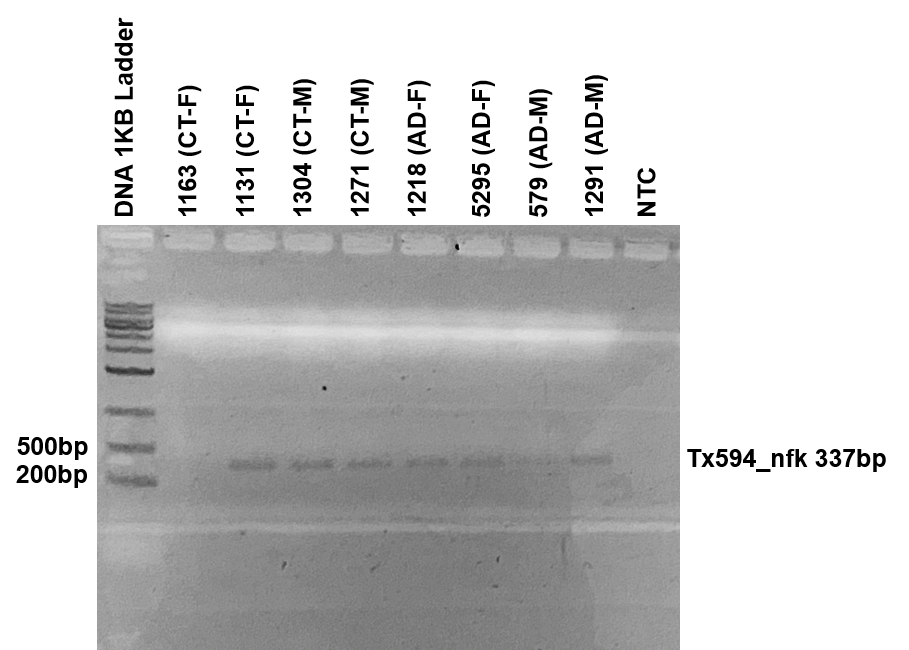

**Supplementary Figure 7: PCR validation for new high confidence transcript Tx594 (*MTHFS).***

Number above the lanes is the sample id for 8 out of the 12 samples used in this study. The other 4 samples were not included because we ran out of tissue left for generating new cDNA. Labels on the right side of the figure indicate transcript id for the new high-confidence RNA isoform being validated and the expected product length from the PCR primers used to amplify the RNA isoform. Tx594 is a new RNA isoform from the MTHFS gene and was succesfully validated in this gel.

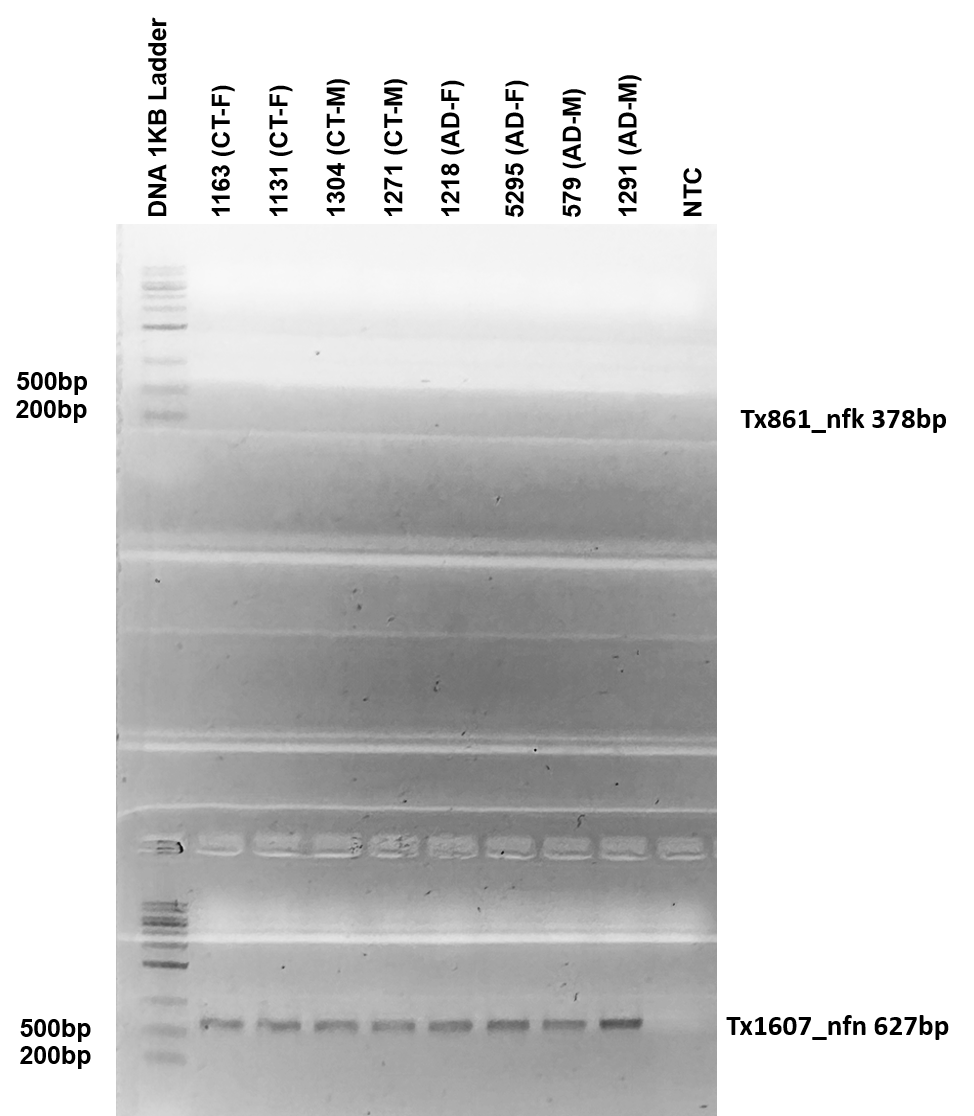

**Supplementary Figure 8: PCR validation for new high confidence transcripts Tx861 (*PSENEN)* and Tx1607 (New gene body).**

Number above the lanes is the sample id for 8 out of the 12 samples used in this study. The other 4 samples were not included because we ran out of tissue left for generating new cDNA. Labels on the right side of the figure indicate transcript id for the new high-confidence RNA isoform being validated and the expected product length from the PCR primers used to amplify the RNA isoform. Tx861 is a new RNA isoform from the *PSENEN* gene and was not validated in this gel, however it was validated through RT-qPCR (**Supplementary Table 5**). Tx2603 is a new RNA isoform ffrom a newly discovered gene body and was succesfully validated in this gel.

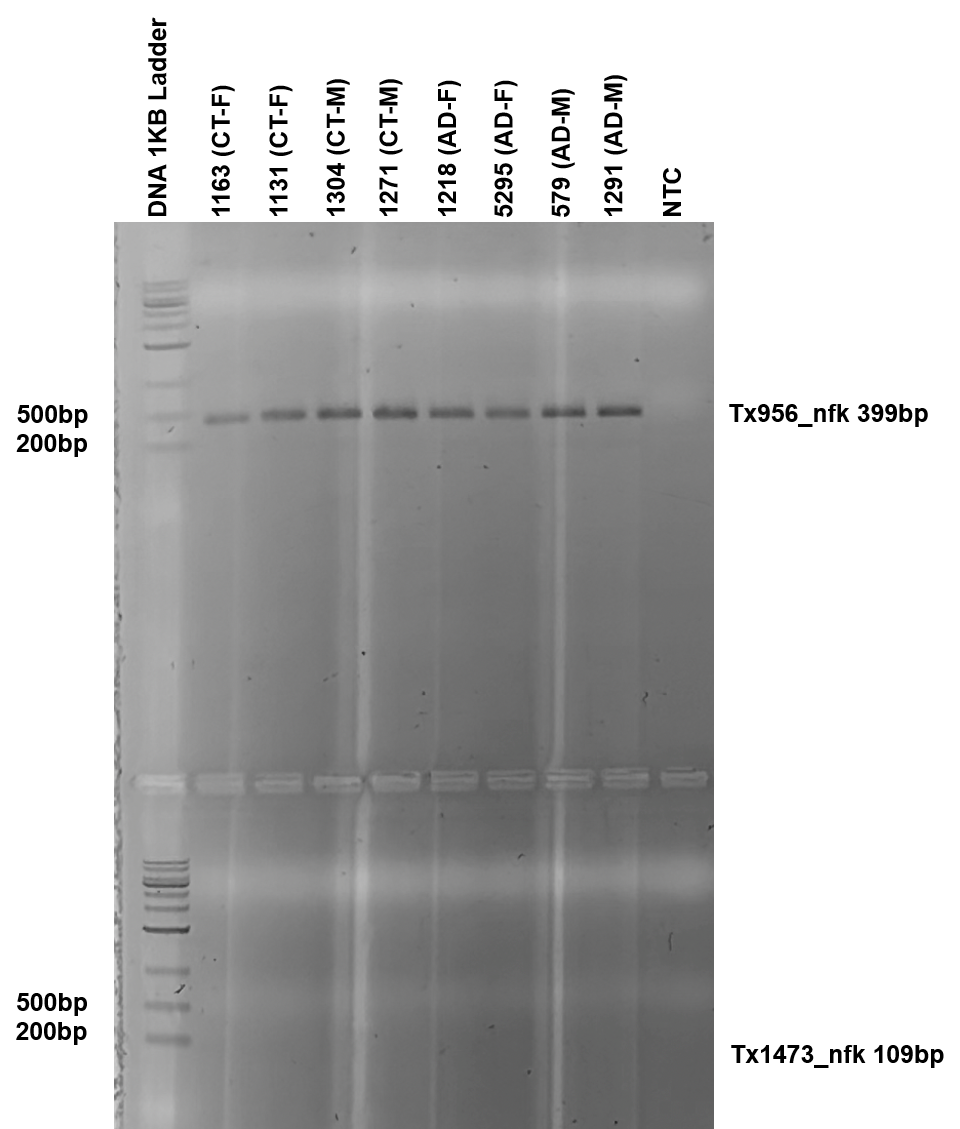

**Supplementary Figure 9: PCR validation for new high confidence transcripts Tx956 (*CAMKMT)* and Tx1473 (*CPLX2*).**

Number above the lanes is the sample id for 8 out of the 12 samples used in this study. The other 4 samples were not included because we ran out of tissue left for generating new cDNA. Labels on the right side of the figure indicate transcript id for the new high-confidence RNA isoform being validated and the expected product length from the PCR primers used to amplify the RNA isoform. Tx956 is a new RNA isoform from the *CAMKMT* gene and was succesfully validated in this gel. Tx1473 is a new RNA isoform from the *CPLX2* gene and was not validated in this gel and was also not validated through RT-qPCR (**Supplementary Table 5**).

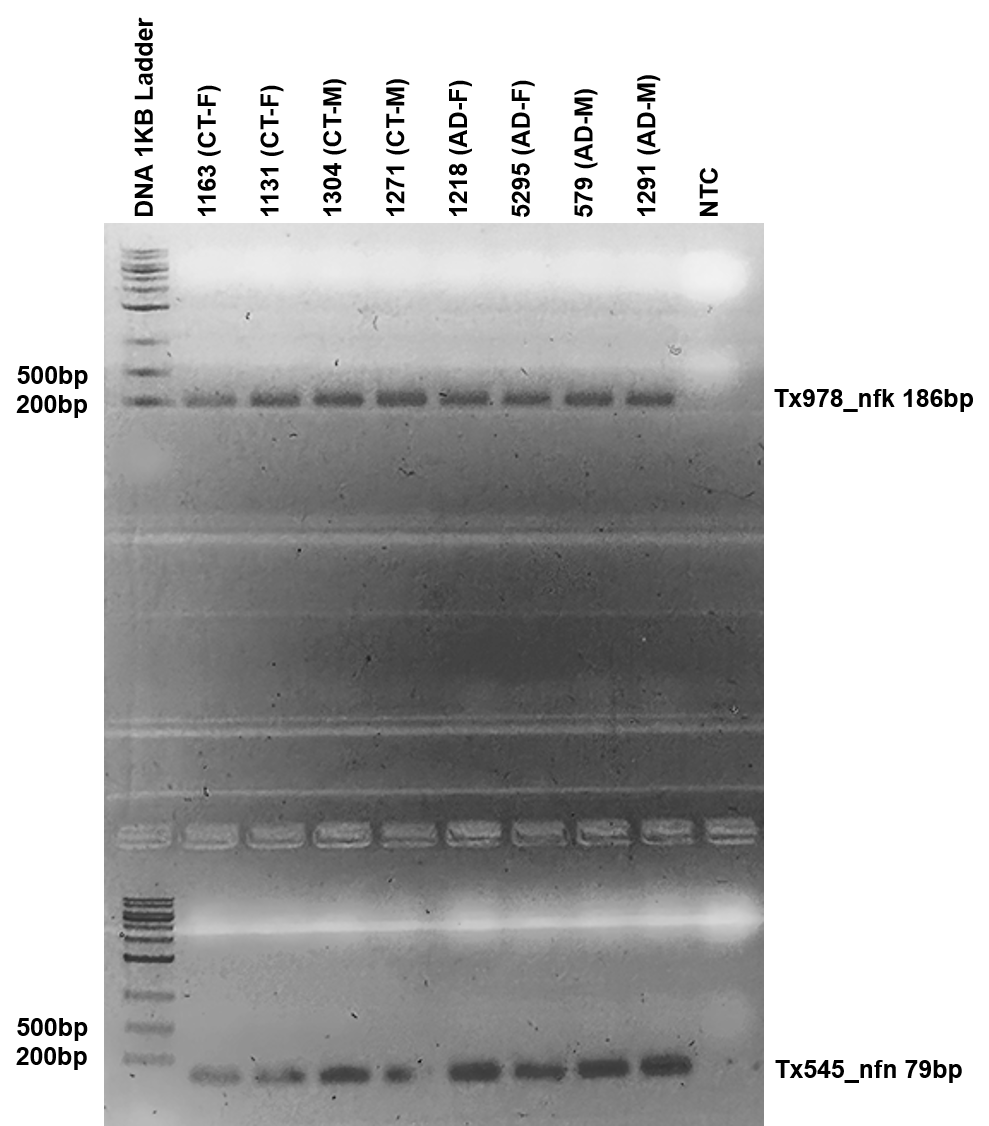

**Supplementary Figure 10: PCR validation for new high confidence transcripts Tx978 (*DGUOK)* and Tx545 (New gene body).**

Number above the lanes is the sample id for 8 out of the 12 samples used in this study. The other 4 samples were not included because we ran out of tissue left for generating new cDNA. Labels on the right side of the figure indicate transcript id for the new high-confidence RNA isoform being validated and the expected product length from the PCR primers used to amplify the RNA isoform. Tx978 is a new RNA isoform from the *DGUOK* gene and was succesfully validated in this gel. Tx545 is a new RNA isoform from a newly discovered gene body and was succesfully validated in this gel.

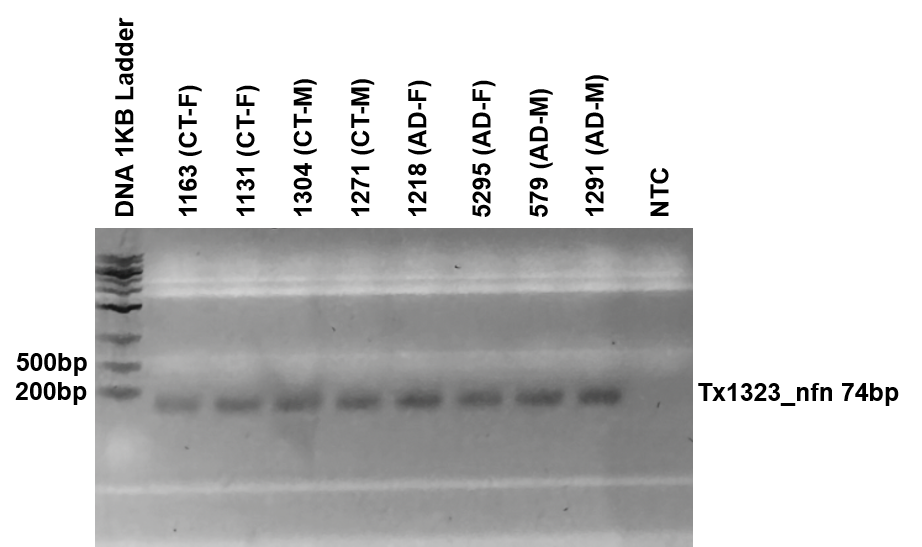

**Supplementary Figure 11: PCR validation for new high confidence transcript Tx1323 (New gene body*).***

Number above the lanes is the sample id for 8 out of the 12 samples used in this study. The other 4 samples were not included because we ran out of tissue left for generating new cDNA. Labels on the right side of the figure indicate transcript id for the new high-confidence RNA isoform being validated and the expected product length from the PCR primers used to amplify the RNA isoform. Tx1323 is a new RNA isoform from a newly discovered gene body and was succesfully validated in this gel.

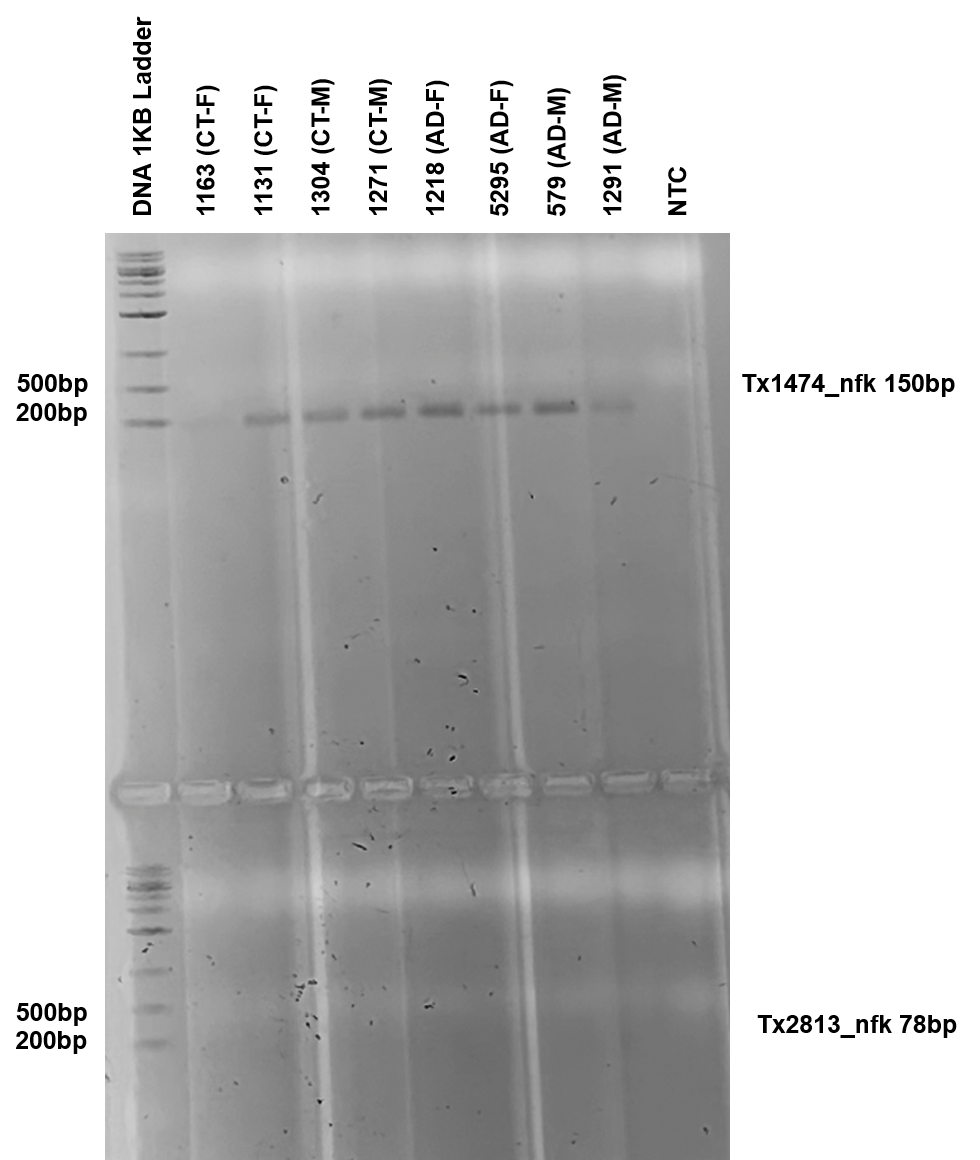

**Supplementary Figure 12: PCR validation for new high confidence transcripts Tx1474 (*CPLX2)* and Tx2813 (*PSENEN*).**

Number above the lanes is the sample id for 8 out of the 12 samples used in this study. The other 4 samples were not included because we ran out of tissue left for generating new cDNA. Labels on the right side of the figure indicate transcript id for the new high-confidence RNA isoform being validated and the expected product length from the PCR primers used to amplify the RNA isoform. Tx1474 is a new RNA isoform from the *CPLX2* gene and was succesfully validated in this gel. Tx2813 is a new RNA isoform from the *PSENEN* gene and was succesfully validated in this gel, however it was validated through RT-qPCR (**Supplementary Table 5**).

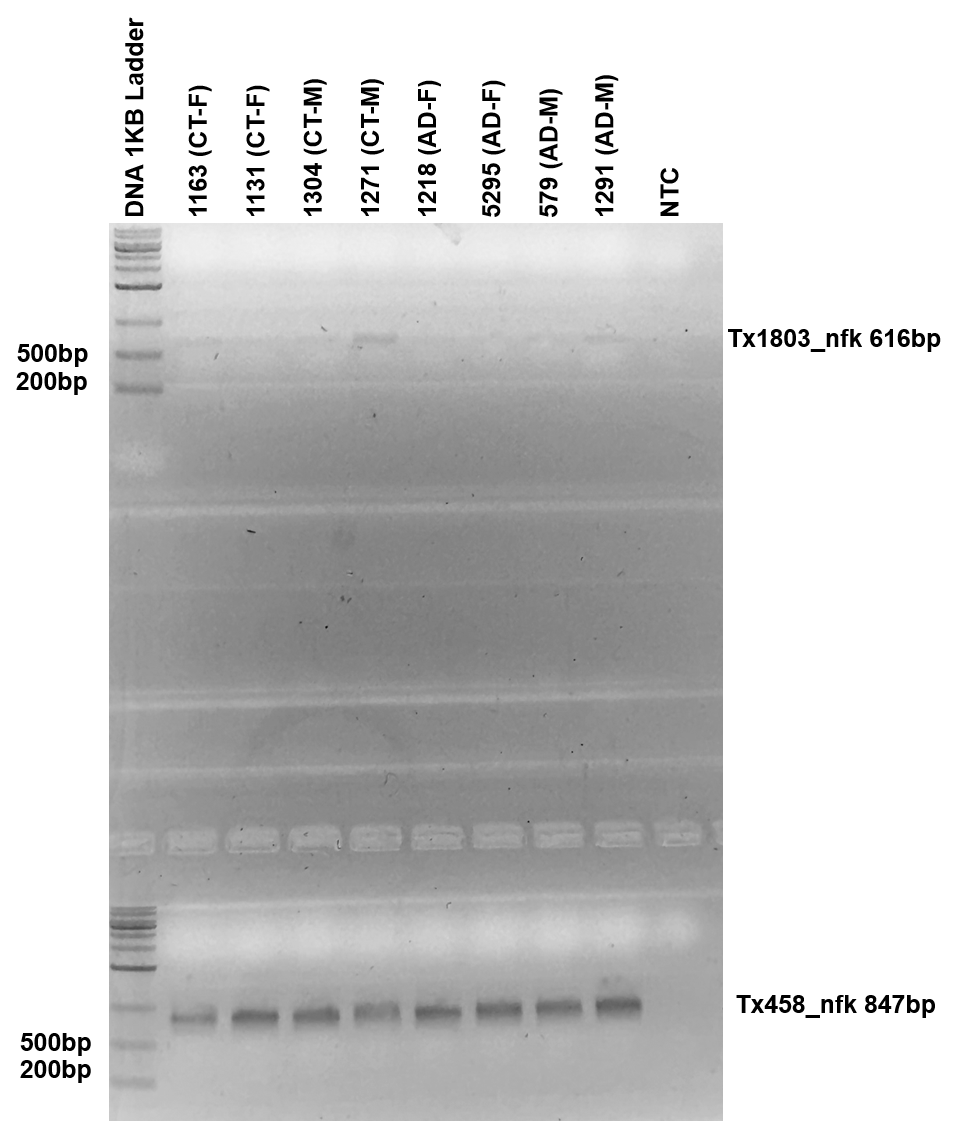

**Supplementary Figure 13: PCR validation for new high confidence transcripts Tx1803 (*TMOD1)* and Tx458 (*NAA16*).**

Number above the lanes is the sample id for 8 out of the 12 samples used in this study. The other 4 samples were not included because we ran out of tissue left for generating new cDNA. Labels on the right side of the figure indicate transcript id for the new high-confidence RNA isoform being validated and the expected product length from the PCR primers used to amplify the RNA isoform. Tx1803 is a new RNA isoform from the *TMOD1* gene and was not validated in this gel, however it was validated through RT-qPCR (**Supplementary Table 5**). Tx458 is a new RNA isoform from the *NAA16* gene and was succesfully validated in this gel.

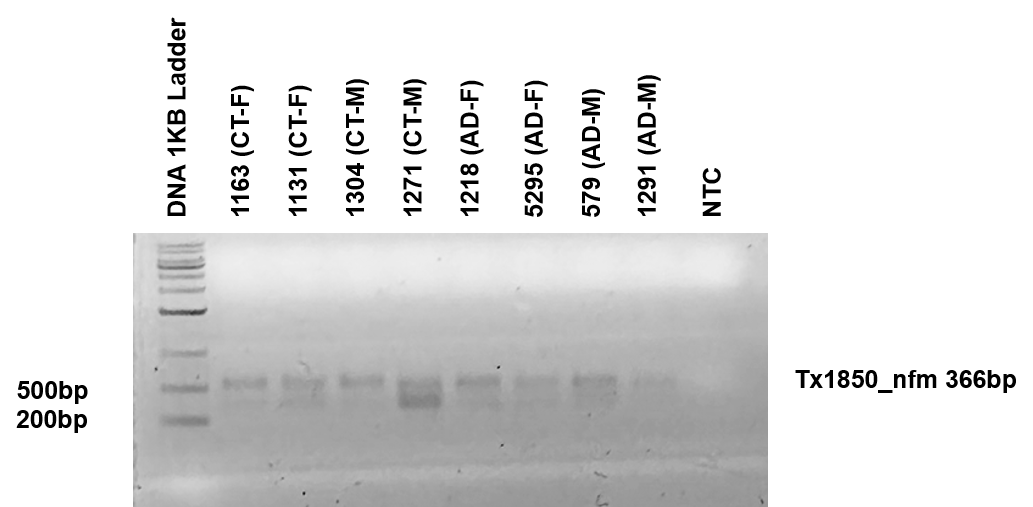

**Supplementary Figure 14: PCR validation for new high confidence transcript Tx1850 (New spliced mitochondrial isoform).**

Number above the lanes is the sample id for 8 out of the 12 samples used in this study. The other 4 samples were not included because we ran out of tissue left for generating new cDNA. Labels on the right side of the figure indicate transcript id for the new high-confidence RNA isoform being validated and the expected product length from the PCR primers used to amplify the RNA isoform. Tx1850 is a new spliced RNA isoform from mitochondria and was not succesfully validated due to double bands shown in gel that indicate promiscous primer binding. Because of promiscous primer binding we did not attempt to validate it through RT-qPCR.

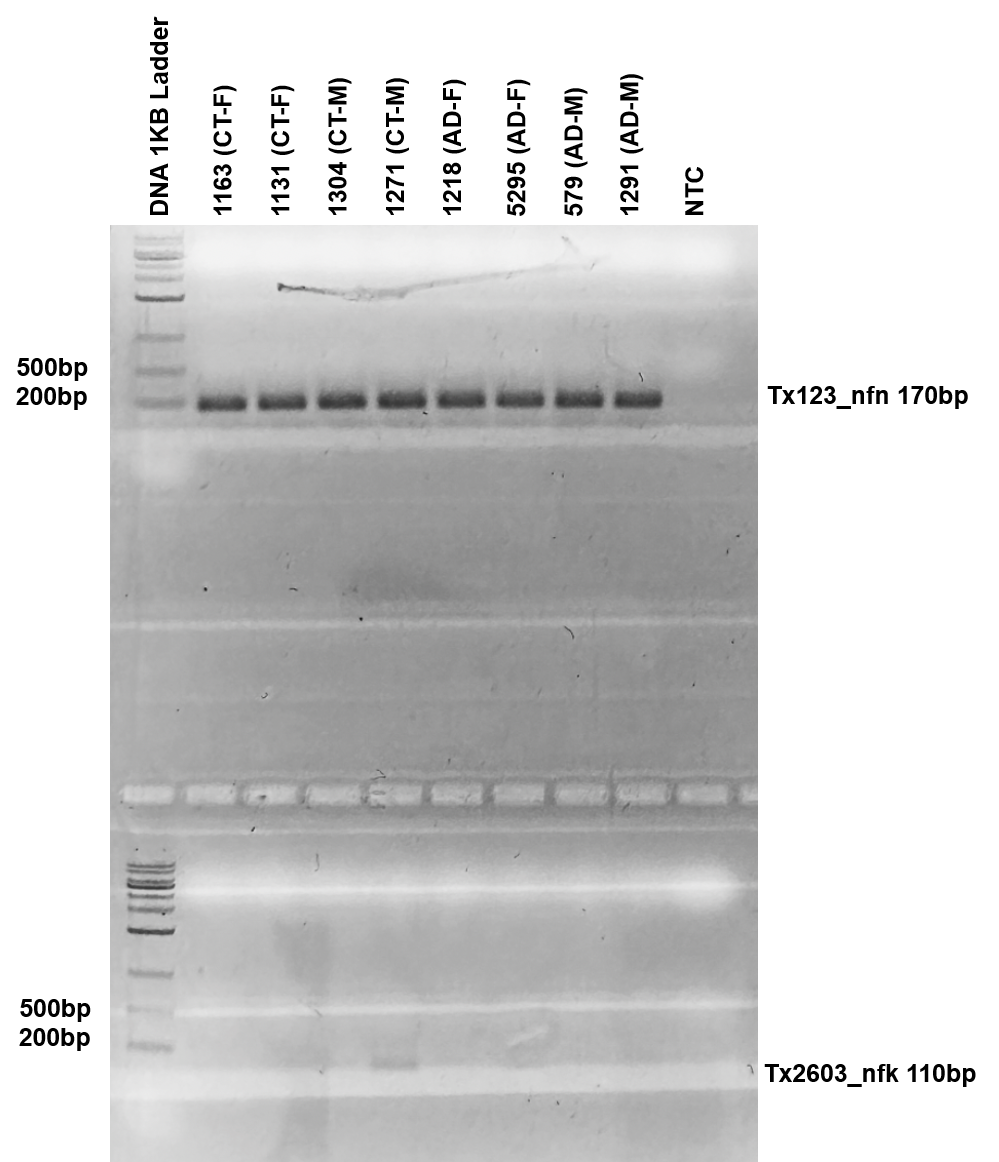

**Supplementary Figure 15: PCR validation for new high confidence transcripts Tx123 (New gene body*)* and Tx2603 (*WDR4*).**

Number above the lanes is the sample id for 8 out of the 12 samples used in this study. The other 4 samples were not included because we ran out of tissue left for generating new cDNA. Labels on the right side of the figure indicate transcript id for the new high-confidence RNA isoform being validated and the expected product length from the PCR primers used to amplify the RNA isoform. Tx123 is a new RNA isoform from a newly discovered gene body and was succesfully validated in this gel. Tx2603 is a new RNA isoform for the *WRD4* gene and was not validated in this gel, however it was validated through RT-qPCR (**Supplementary Table 5**).

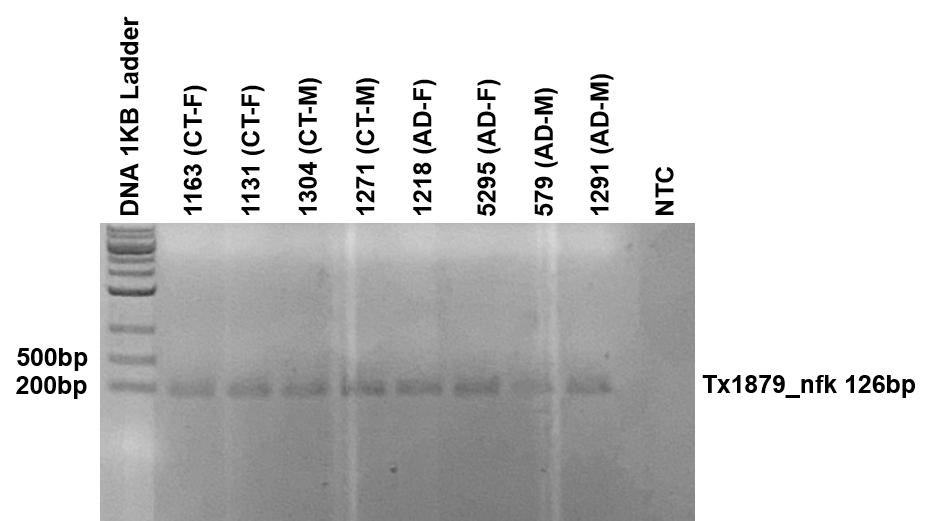

**Supplementary Figure 16: PCR validation for new high confidence transcript Tx1879 (*MAOB).***

Number above the lanes is the sample id for 8 out of the 12 samples used in this study. The other 4 samples were not included because we ran out of tissue left for generating new cDNA. Labels on the right side of the figure indicate transcript id for the new high-confidence RNA isoform being validated and the expected product length from the PCR primers used to amplify the RNA isoform. Tx1879 is a new RNA isoform from the *MAOB* gene and was succesfully validated in this gel.

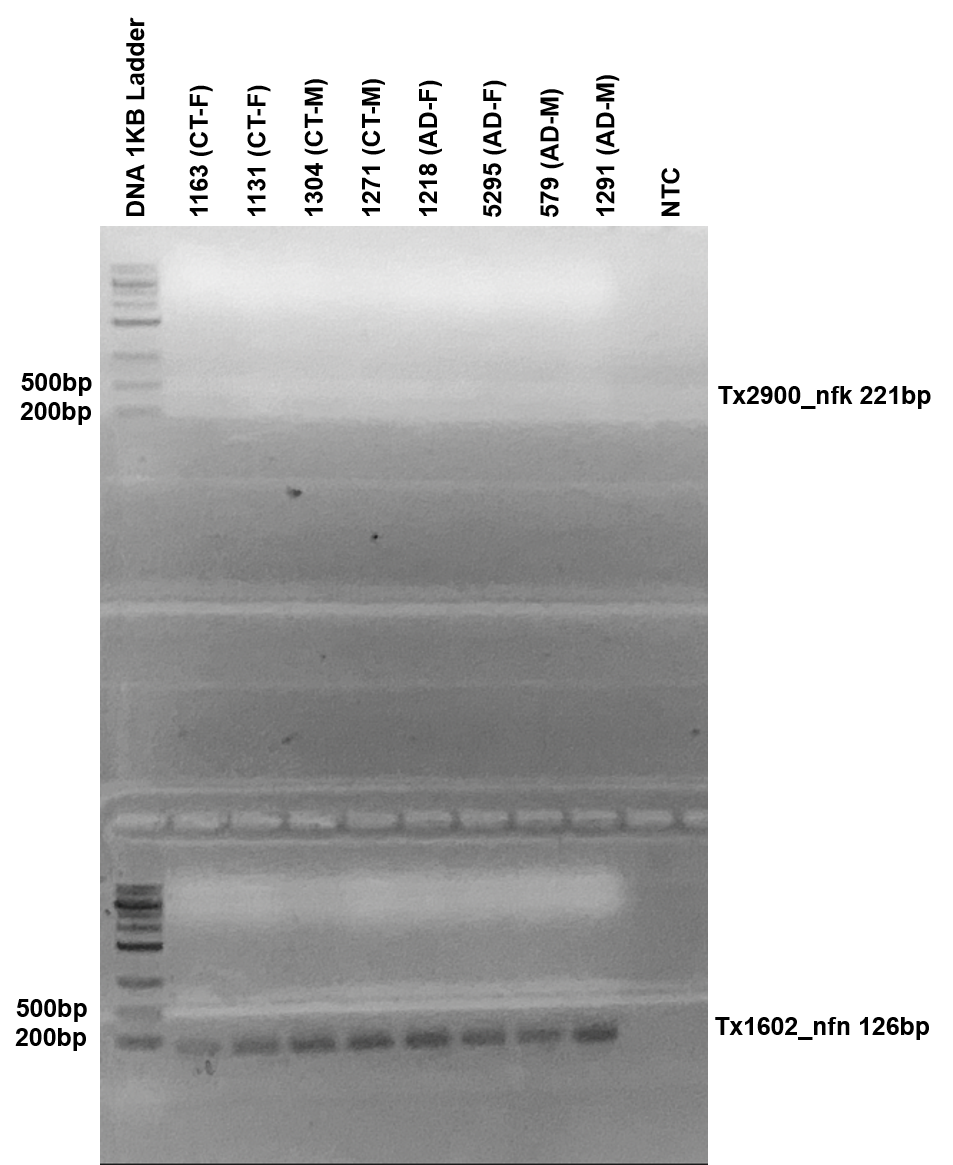

**Supplementary Figure 17: PCR validation for new high confidence transcripts Tx2900 (*ENSG00000255122)* and Tx1602 (New gene body).**

Number above the lanes is the sample id for 8 out of the 12 samples used in this study. The other 4 samples were not included because we ran out of tissue left for generating new cDNA. Labels on the right side of the figure indicate transcript id for the new high-confidence RNA isoform being validated and the expected product length from the PCR primers used to amplify the RNA isoform. Tx2900 is a new RNA isoform from the *ENSG00000255122* gene and was not validated in this gel, however it was validated through RT-qPCR (**Supplementary Table 5**). Tx1602 is a new RNA isoform from a newly discovered gene body and was succesfully validated in this gel.

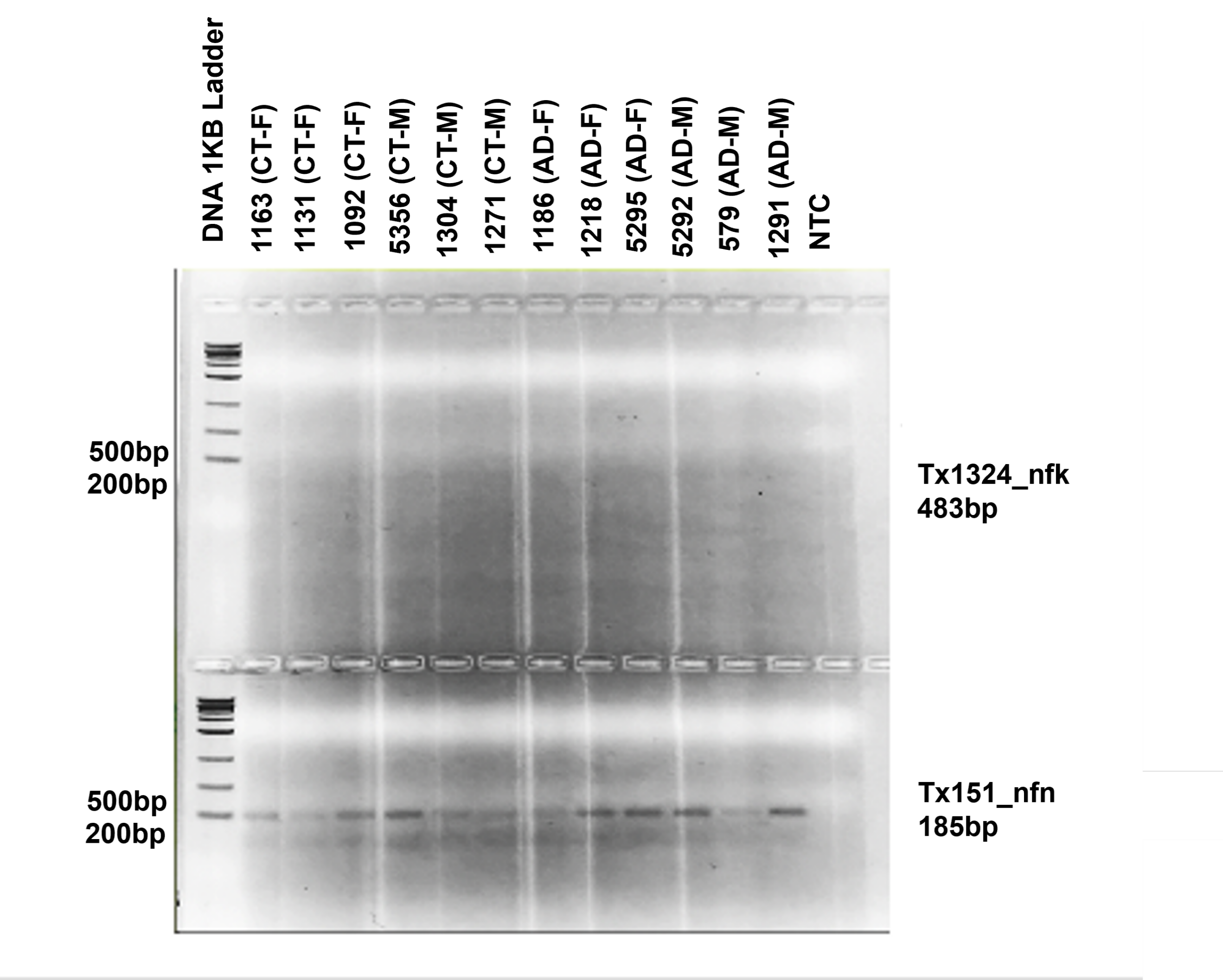

**Supplementary Figure 18: PCR validation for new high confidence transcripts Tx1324 (*NELFA)* and Tx151 (*New gene body*).**

Number above the lanes is the sample id for the 12 samples used in this study. Labels on the right side of the figure indicate transcript id for the new high-confidence RNA isoform being validated and the expected product length from the PCR primers used to amplify the RNA isoform. Tx1324 is a new RNA isoform from the *NELFA* gene and was not validated in this gel, but was validated in **Supplementary Figure 6**. Tx151 is a new RNA isoform from a newly discovered gene body and was validated in this gel (double bands), but was succesfully validated in **Supplementary Figure 5**.

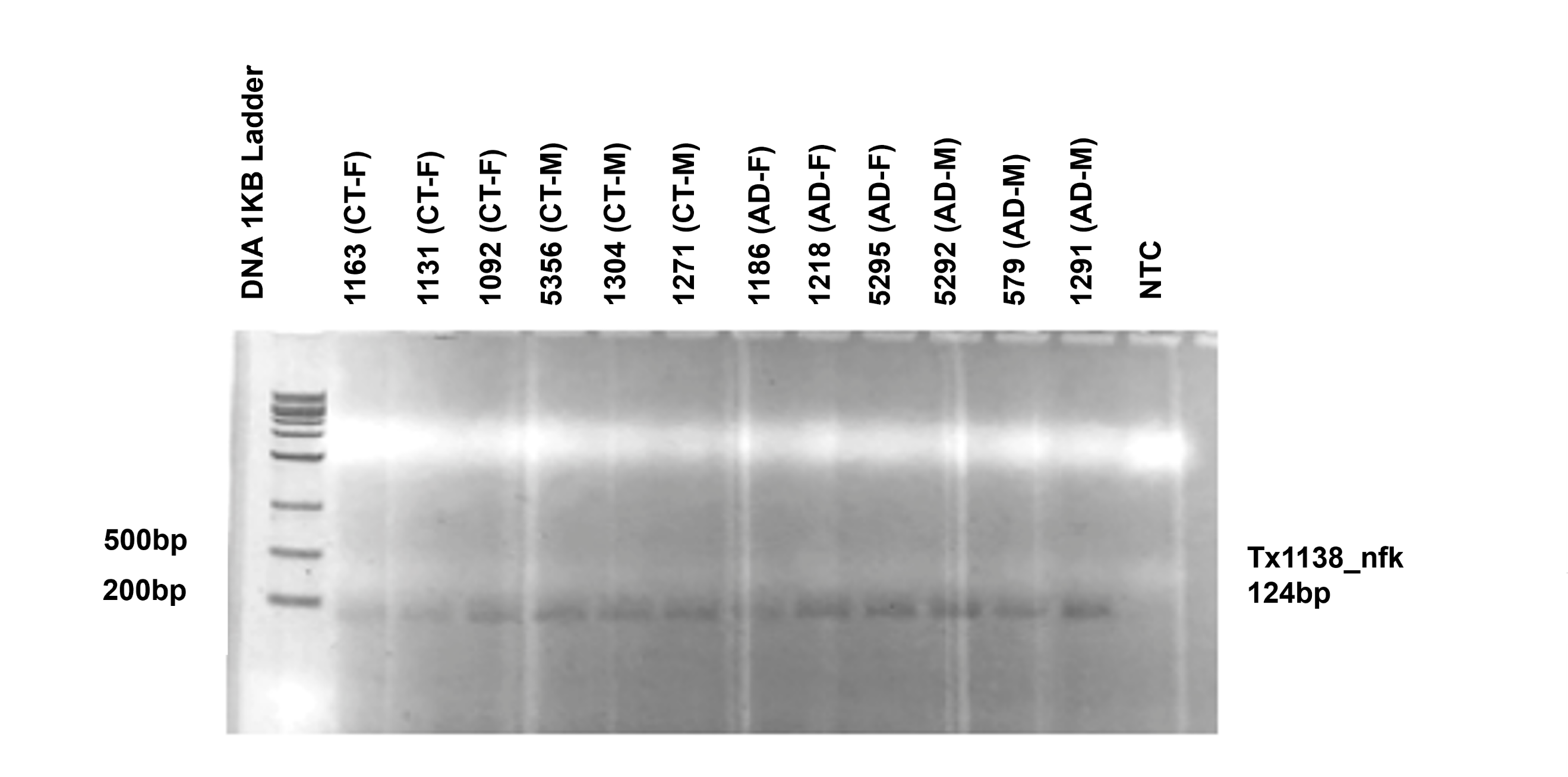

**Supplementary Figure 19: PCR validation for new high confidence transcript Tx1138 (*IFNAR1)***

Number above the lanes is the sample id for the 12 samples used in this study. Labels on the right side of the figure indicate transcript id for the new high-confidence RNA isoform being validated and the expected product length from the PCR primers used to amplify the RNA isoform. Tx1138 is a new RNA isoform from the *IFNAR1* gene and was succesfully validated.

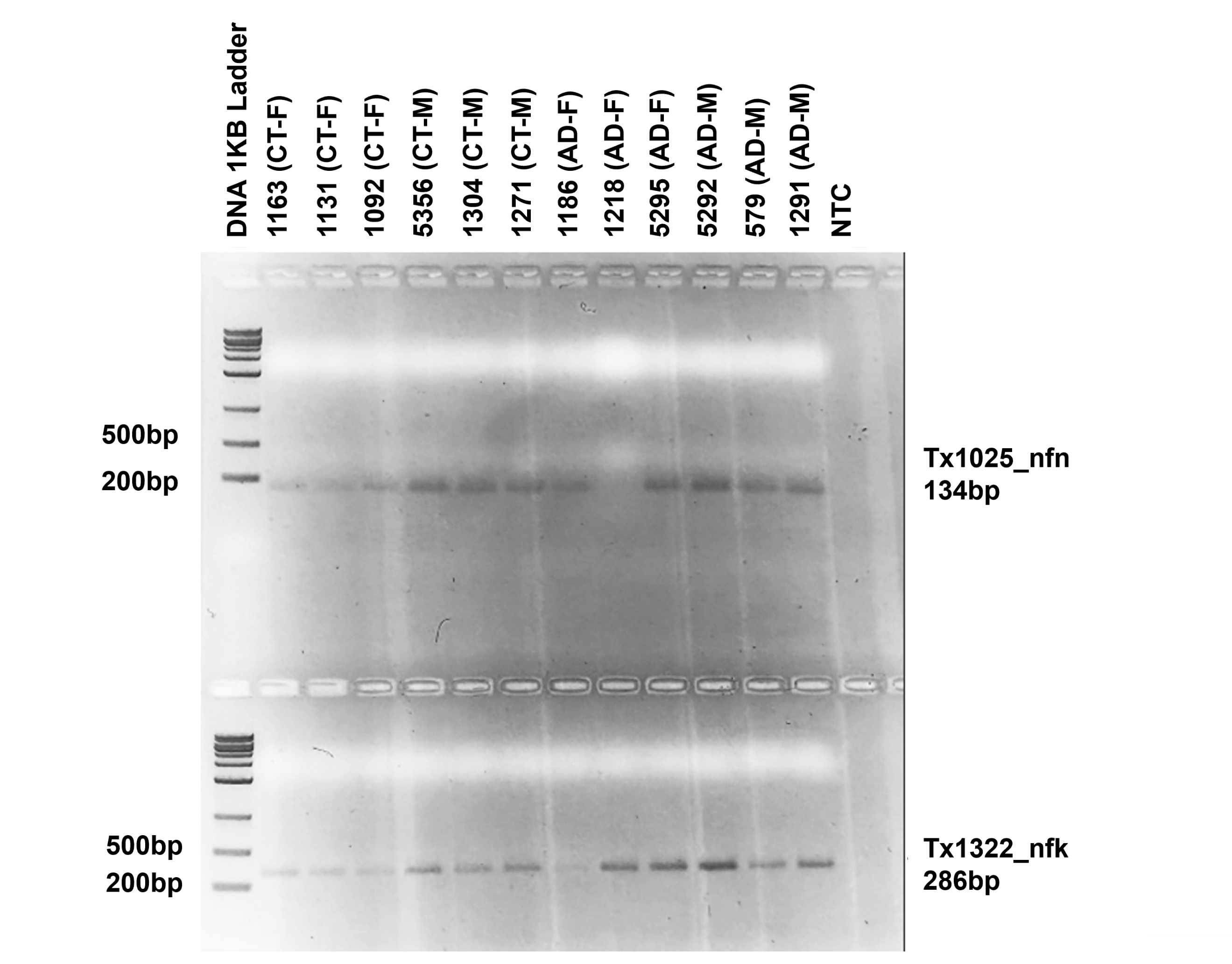

**Supplementary Figure 20: PCR validation for new high confidence transcripts Tx1025 (New gene body) and Tx1322 (*SLC26A1)*.**

Number above the lanes is the sample id for the 12 samples used in this study. Labels on the right side of the figure indicate transcript id for the new high-confidence RNA isoform being validated and the expected product length from the PCR primers used to amplify the RNA isoform. Tx1025 is a new RNA isoform from a newly discovered gene body and was succesfully validated. Tx1322 is a new RNA isoform for the *SLC26A1* gene and was succesfully validated.

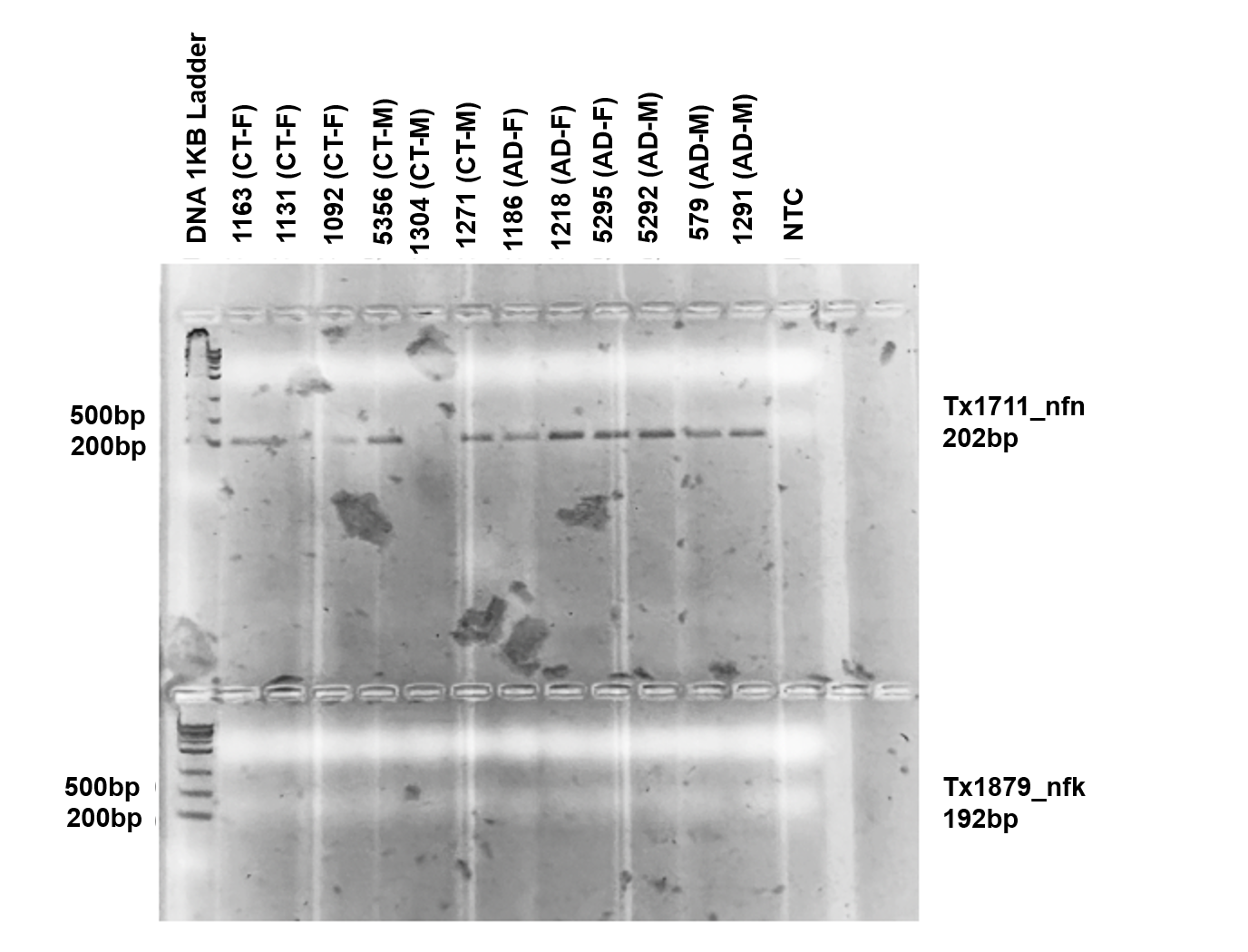

**Supplementary Figure 21: PCR validation for new high confidence transcripts Tx1711 (New gene body) and Tx1879 (*MAOB*).**

Number above the lanes is the sample id for the 12 samples used in this study. Labels on the right side of the figure indicate transcript id for the new high-confidence RNA isoform being validated and the expected product length from the PCR primers used to amplify the RNA isoform. Tx1711 is a new RNA isoform from a newly discovered gene body and was succesfully validated. Tx1879 is a new RNA isoform for the *MAOB* gene and was not validated in this gel, but was validated with a different primer pair in **Supplementary Figure 16**.

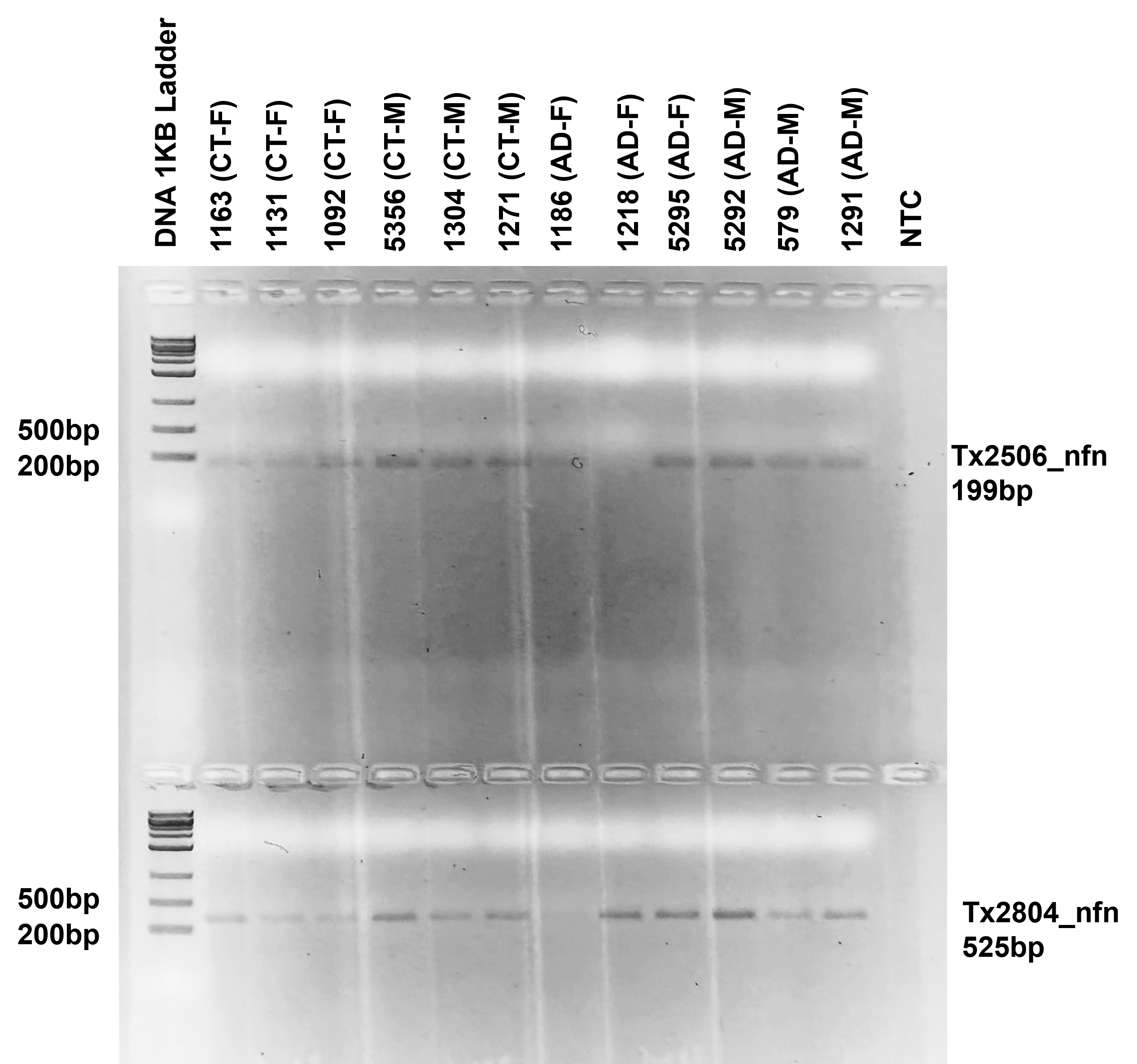

**Supplementary Figure 22: PCR validation for new high confidence transcripts Tx2506 (New gene body) and Tx2804 (New gene body).**

Number above the lanes is the sample id for the 12 samples used in this study. Labels on the right side of the figure indicate transcript id for the new high-confidence RNA isoform being validated and the expected product length from the PCR primers used to amplify the RNA isoform. Tx2506 is a new RNA isoform from a new gene body and was succesfully validated. Tx2804 is a new RNA isoform from a new gene body and was not validated in this figure. We did not consider the Tx2804 validation succesful on this gel because the band is considerably smaller than the predicted 525 nucleotide PCR product. However, we ran another gel and succesfully validated it in  **Supplementary Figure 23**. That PCR reaction was ran with a higher annealing temperature to prevent formation of primer dimers.

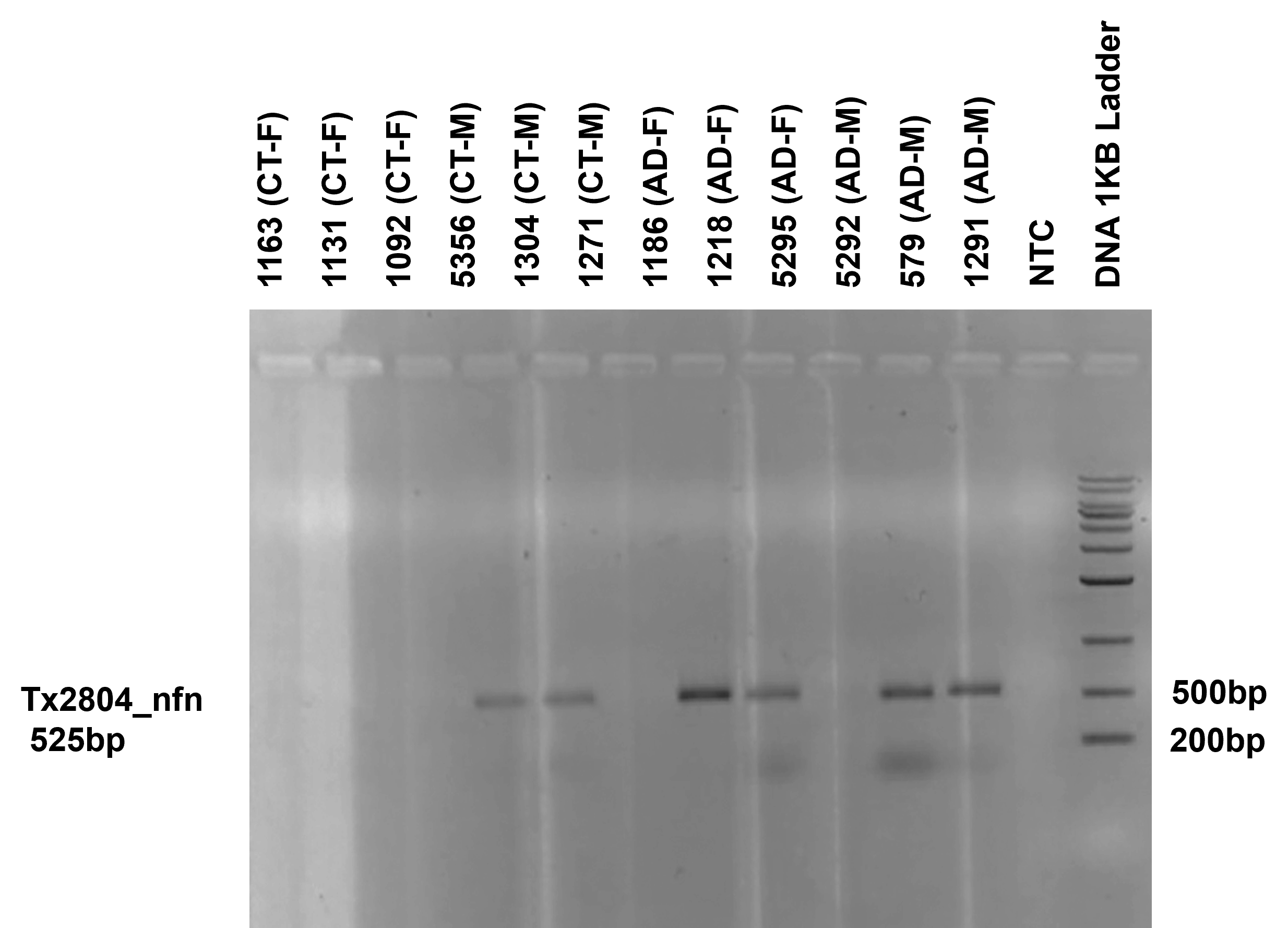

**Supplementary Figure 23: Re-run of PCR validation for new high confidence transcript Tx2804 (New gene body).**

Number above the lanes is the sample id for the 12 samples used in this study. Label on the left side of the figure indicate transcript id for the new high-confidence RNA isoform being validated and the expected product length from the PCR primers used to amplify the RNA isoform. Tx2804 is a new RNA isoform from a new gene body and was succesfully validated.

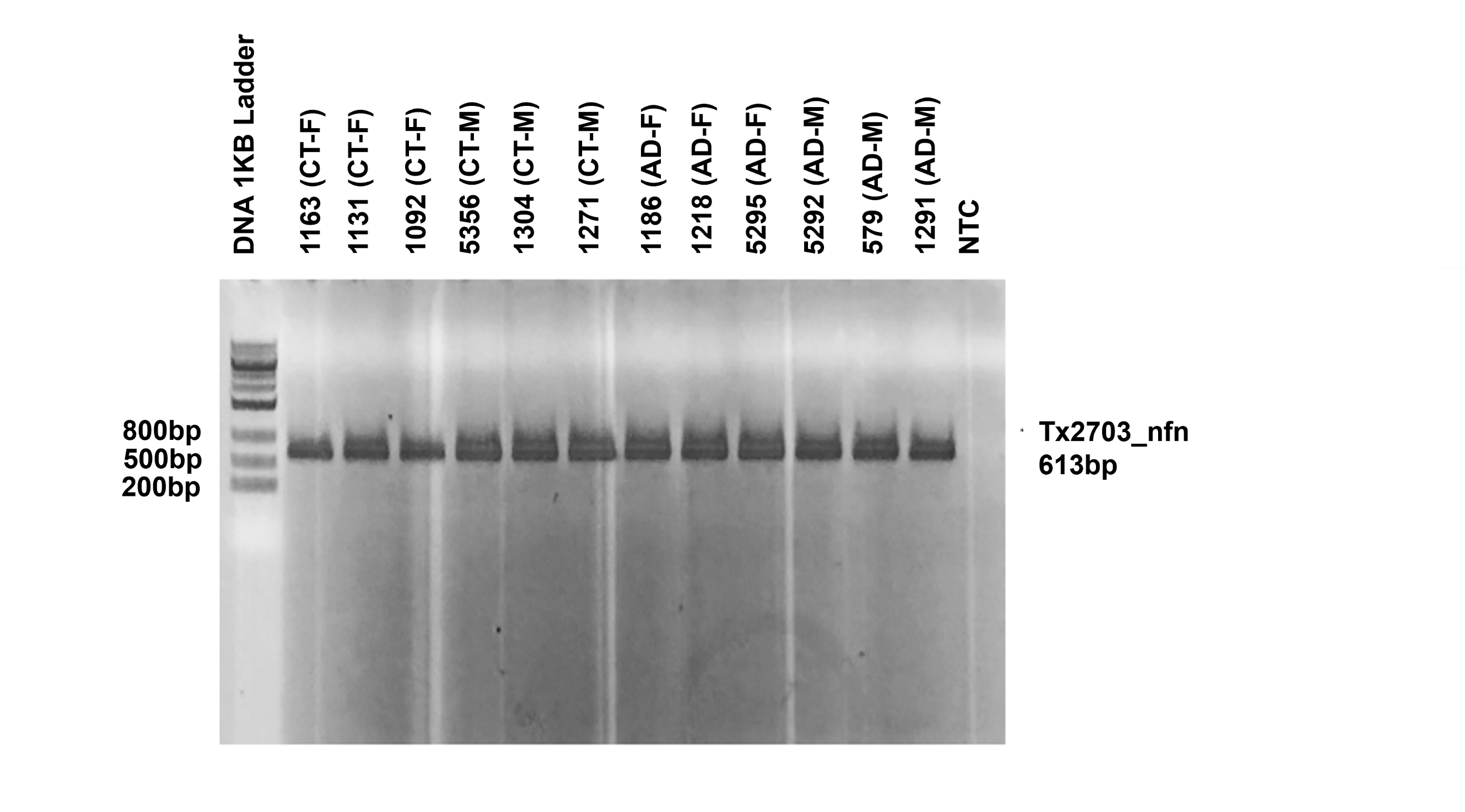

**Supplementary Figure 24: PCR validation for new high confidence transcripts Tx2703 (New gene body).**

Number above the lanes is the sample id for the 12 samples used in this study. Labels on the right side of the figure indicate transcript id for the new high-confidence RNA isoform being validated and the expected product length from the PCR primers used to amplify the RNA isoform. Tx2703 is a new RNA isoform from a newly discovered gene body and was succesfully validated.

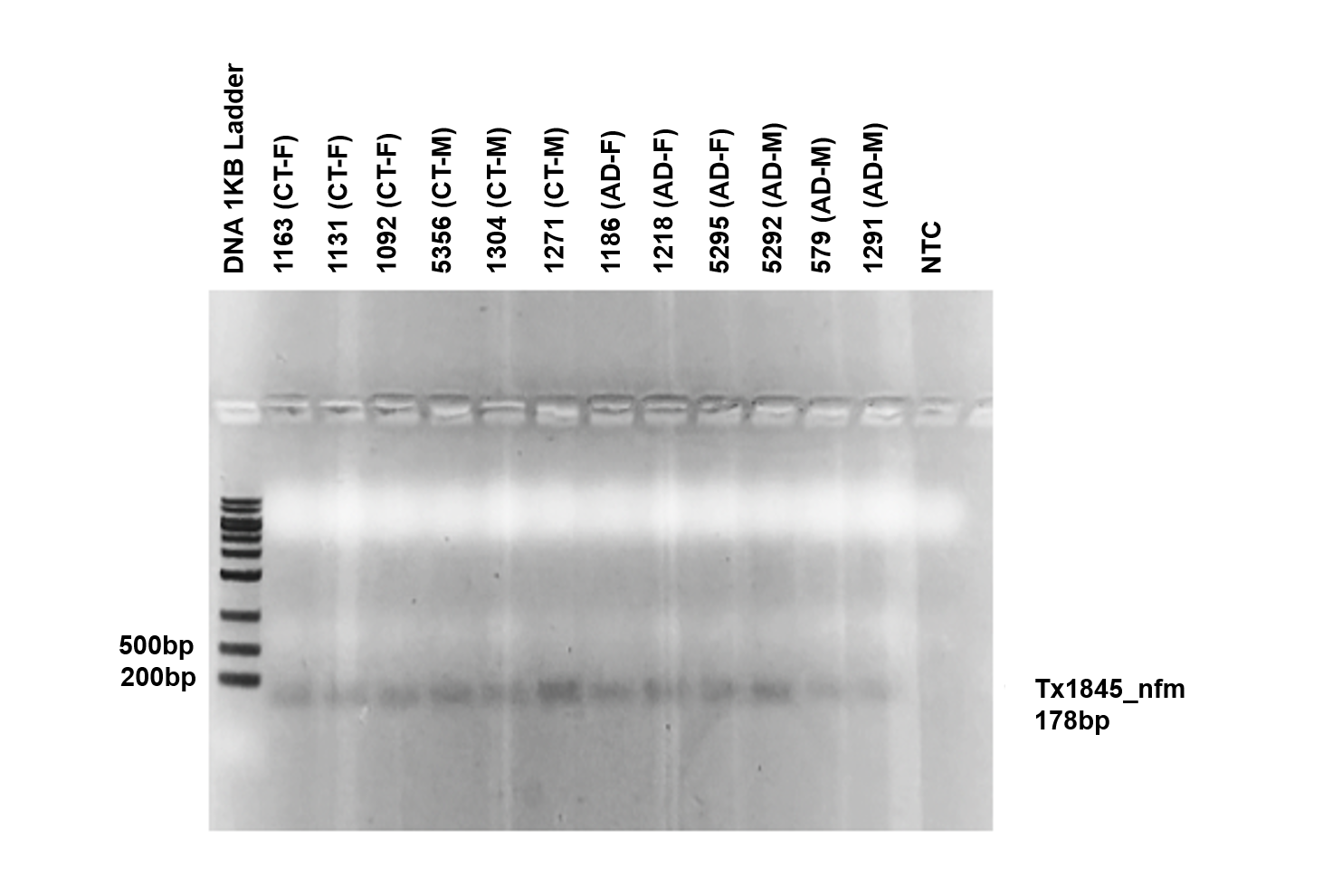

**Supplementary Figure 25: PCR validation for new high confidence transcripts Tx1845 (New spliced mitochondrial isoform).**

Number above the lanes is the sample id for the 12 samples used in this study. Labels on the right side of the figure indicate transcript id for the new high-confidence RNA isoform being validated and the expected product length from the PCR primers used to amplify the RNA isoform. Tx1845 is a new spliced RNA isoform from mitochondria and was succesfully validated.

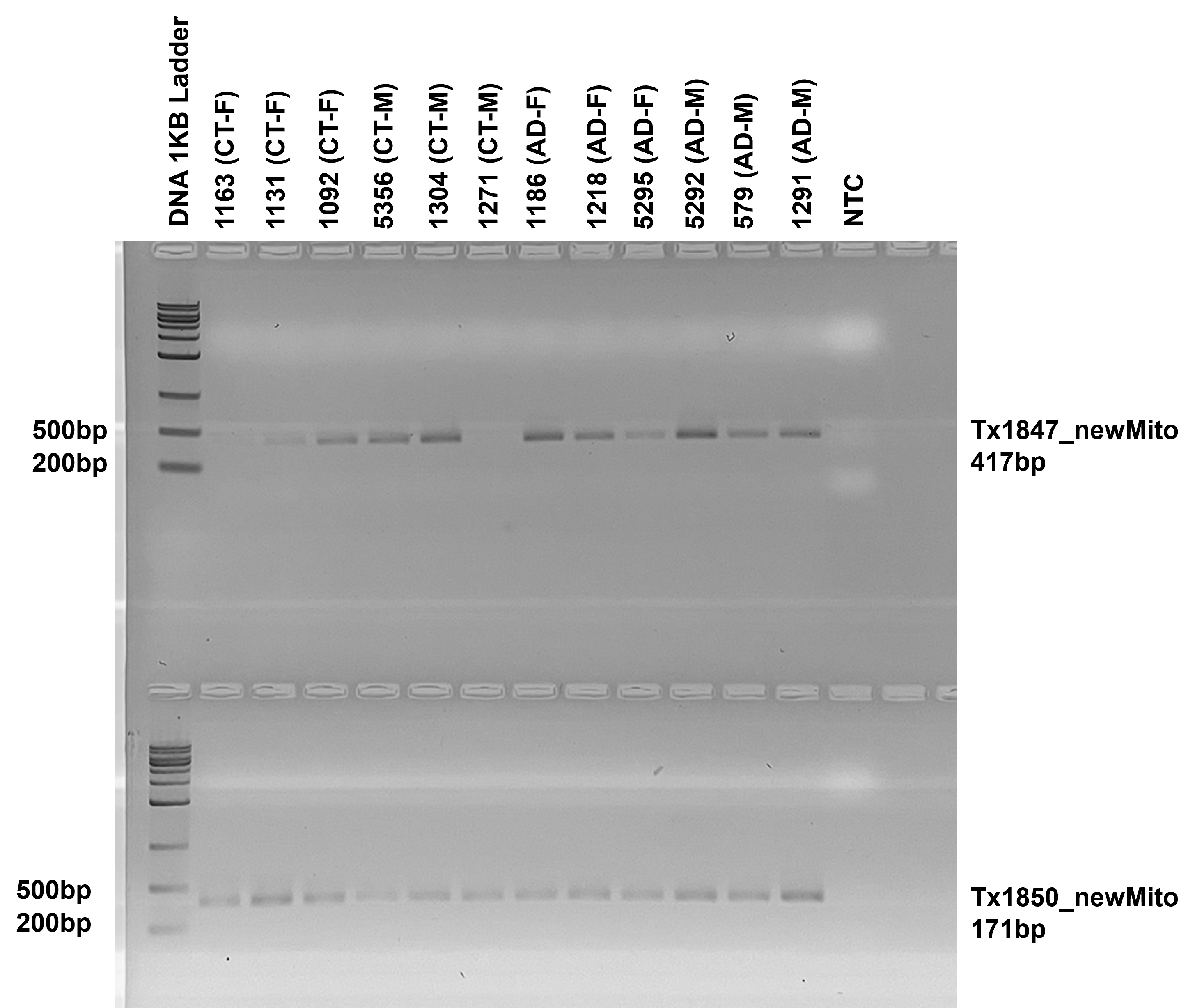

**Supplementary Figure 26: PCR validation for new high confidence transcripts Tx1847 and Tx1850 (New spliced mitochondrial isoforms).**

Number above the lanes is the sample id for the 12 samples used in this study. Labels on the right side of the figure indicate transcript id for the new high-confidence RNA isoform being validated and the expected product length from the PCR primers used to amplify the RNA isoform. Tx1847 is a new spliced RNA isoform from mitochondria and was succesfully validated. Tx1850 is a new spliced RNA isoform from the mitochondria and was not validated. We did not consider the Tx1850 validation succesful on this gel because the band is considerably larger than the predicted 171 nucleotide PCR product.

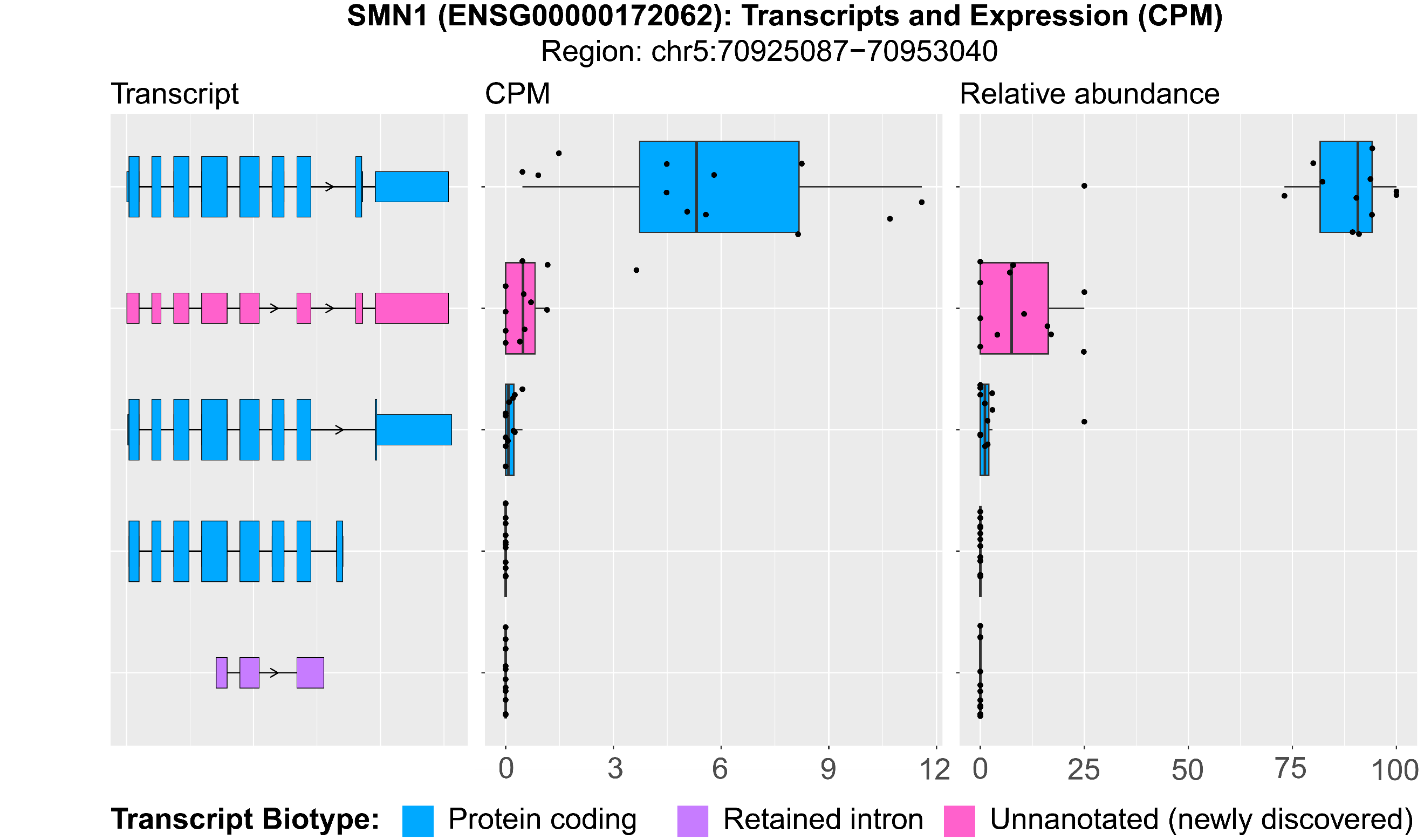

**Supplementary Figure 27: RNA isoform expression and structure for SMN1.**

RNA isoform structure, CPM normalized expression, and relative abundance for SMN1 RNA isoforms. We only show the top 5 most highly expressed RNA isoforms for this gene. The new RNA isoform is shown in pink. This new isoform did not meet our high-confidence threshold (median CPM > 1), but we report it for interest because of the gene’s direct role in neurodegenerative disease.

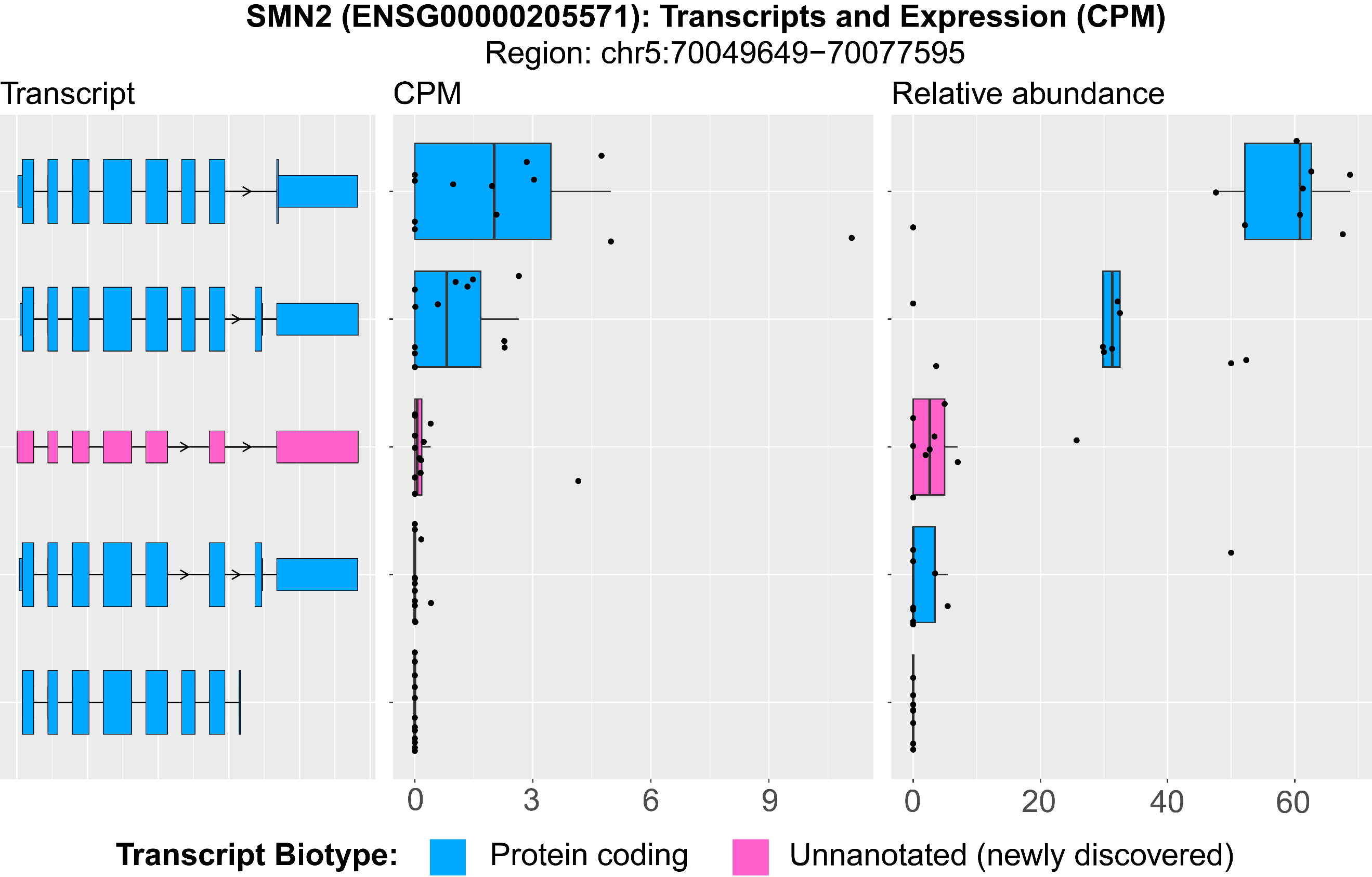

**Supplementary Figure 28: RNA isoform expression and structure for SMN2.**

RNA isoform structure, CPM normalized expression, and relative abundance for SMN2 RNA isoforms. We only show the top 5 most highly expressed RNA isoforms for this gene. The new RNA isoform is shown in pink. This new isoform did not meet our high-confidence threshold (median CPM > 1), but we report it for interest because of the gene’s direct role in neurodegenerative disease.

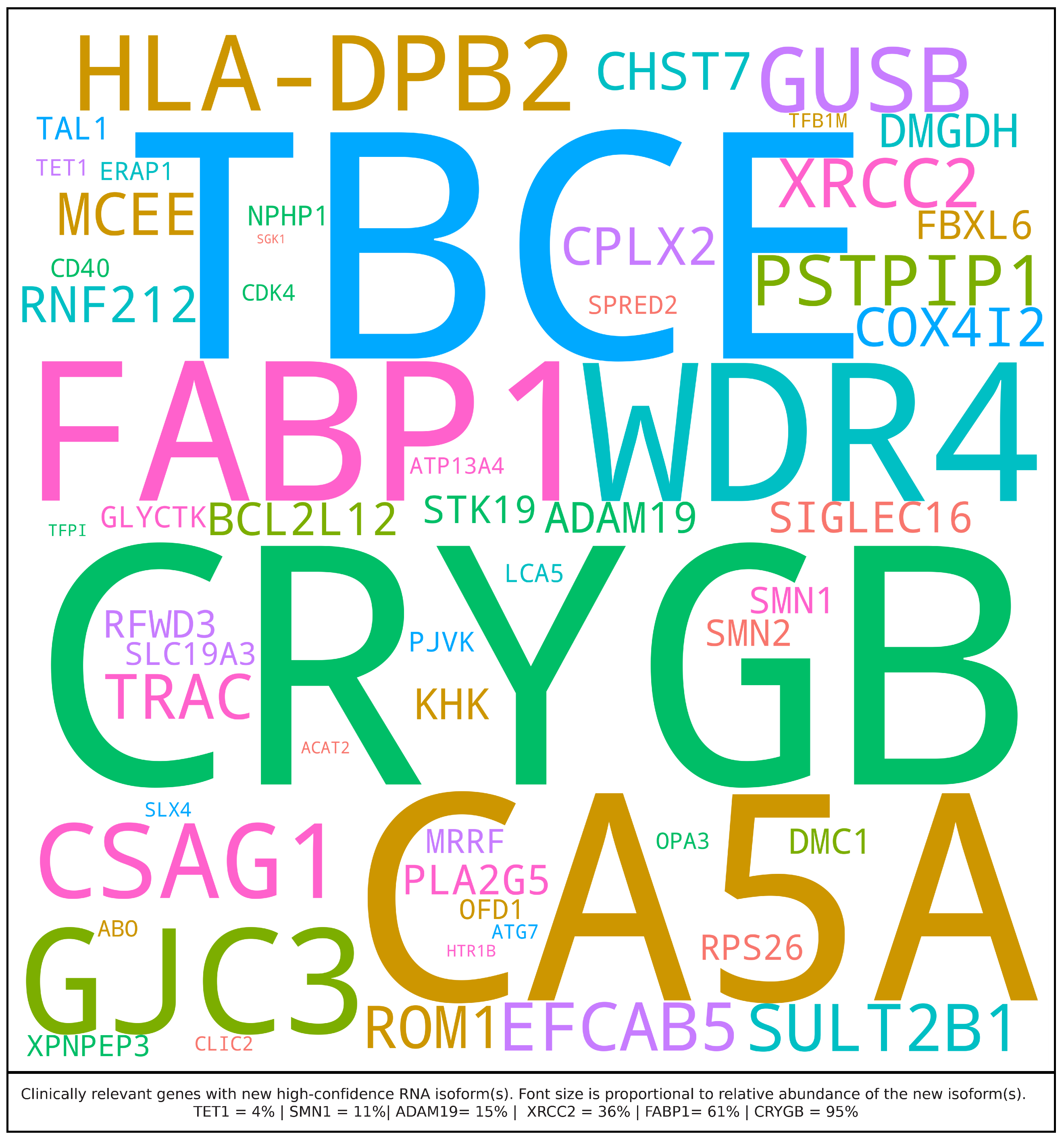

**Supplementary Figure 29: Medically relevant genes with RNA isoforms expressed at median CPM ≤ 1 in human frontal cortex.** Gene names for medically relevant genes where we discovered a new RNA isoform that was not annotated in Ensembl version 107. Only included new RNA isoforms with a median CPM ≤ 1. The size of gene name is proportional to the relative abundance of the new RNA isoform(s).

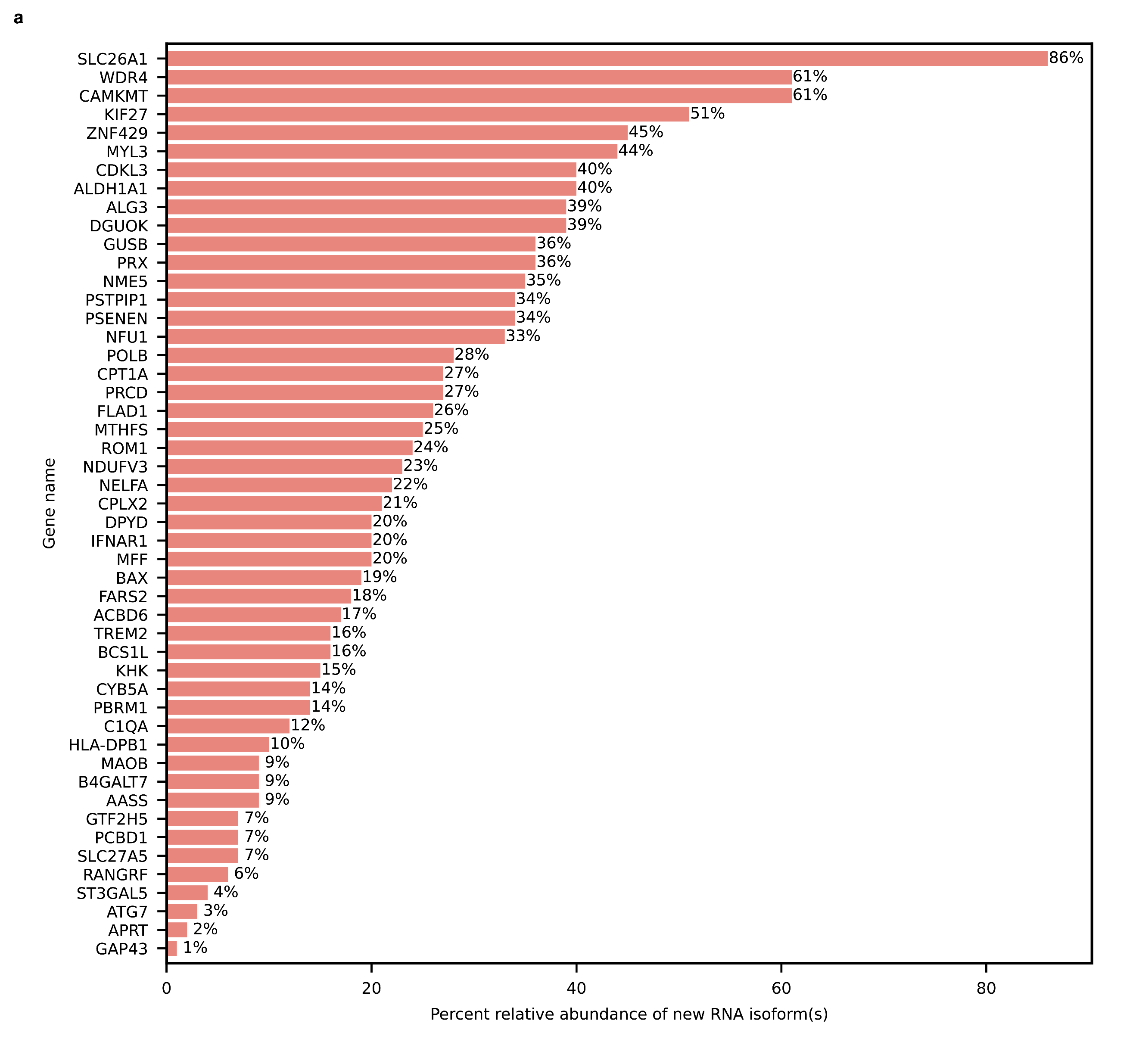

**Supplementary Figure 30: Percent relative abundance of new RNA isoforms in medically relevant genes.**

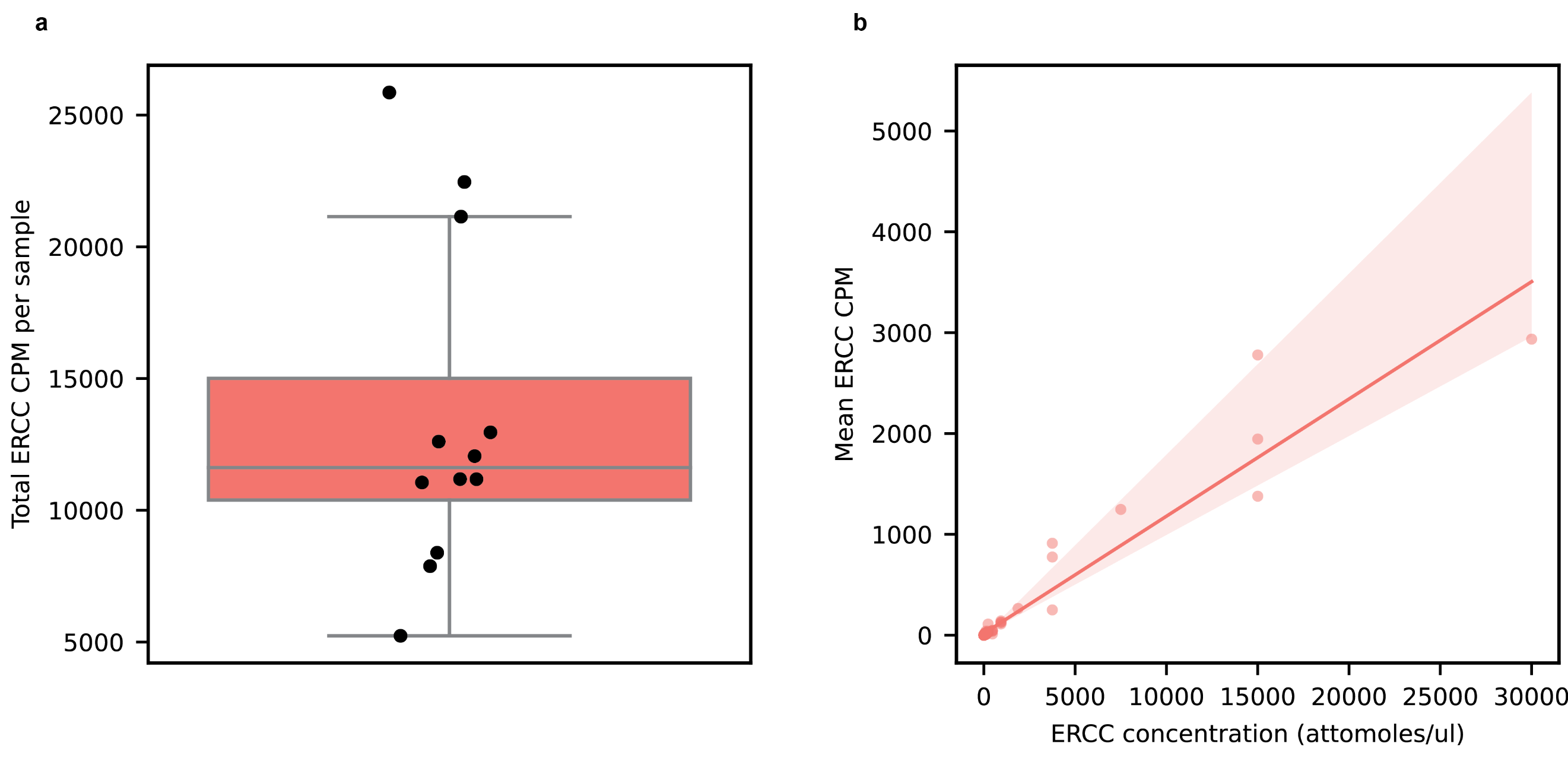

**Supplementary Figure 31: Expression distribution and diversity for genes and transcripts.**

**a,** Sum of CPM for all ERCC control spike-in RNA transcripts for each sample. **b,** Correlation between ERCC transcript concentration and mean ERCC CPM for that same transcript in all 12 samples. Spearman correlation coefficient = 0.98.

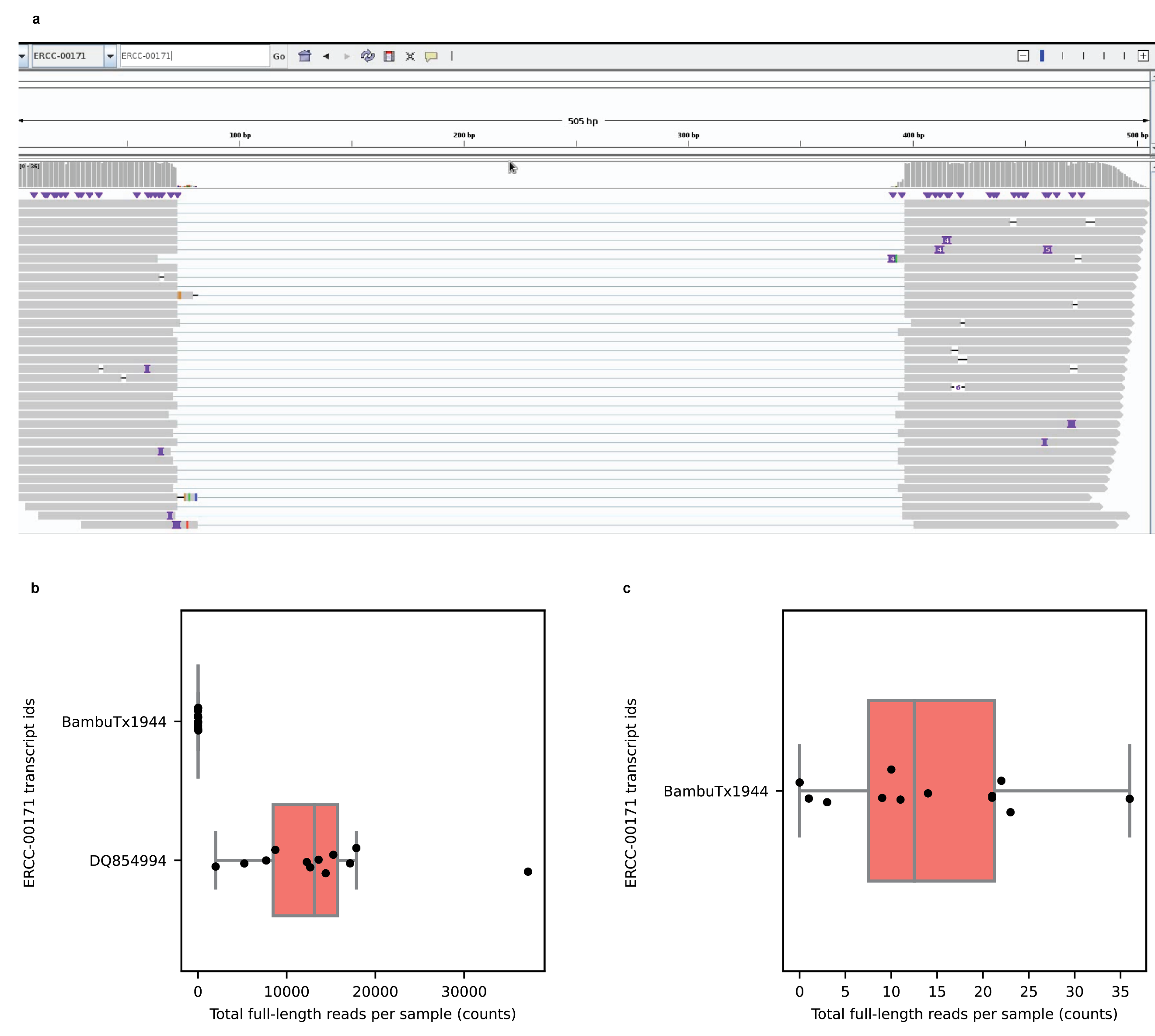

**Supplementary Figure 32: Expression distribution and diversity for genes and transcripts.**

**a,** Interactive Genome Viewer (IGV) screenshot showing the “spliced” ERCC reads for one of our samples. **b,** Comparison between total number of full-length reads for the known ERCC transcript and the new spliced ERCC transcript that shares its nucleotide sequence. Each dot on the graph represents a sample. The new spliced ERCC transcript is present at very low concentration in comparison to the non-spliced ERCC transcript that shares its nucleotide sequence. **c,** Number of full-length reads for the new spliced ERCC transcript in each of the 12 samples.

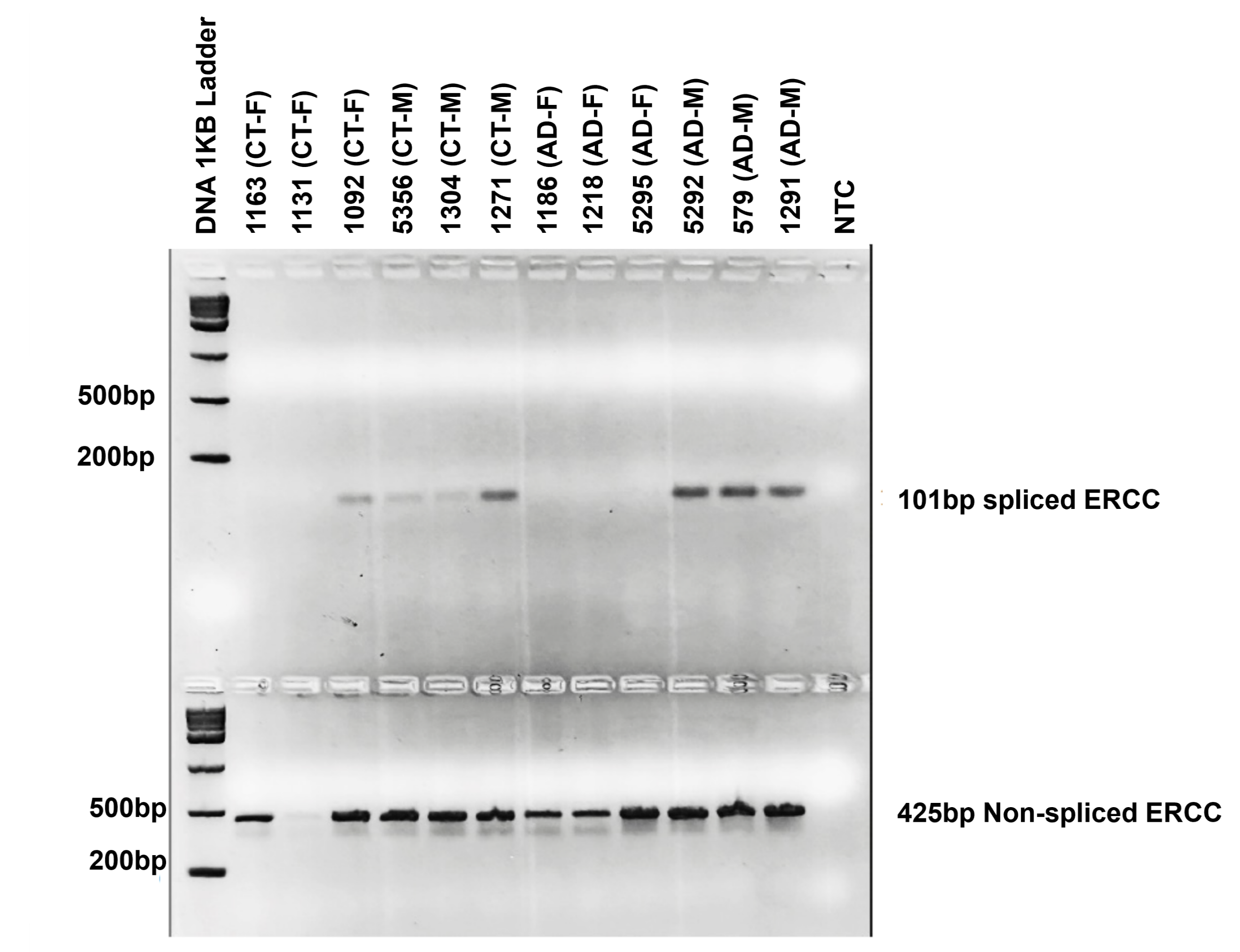

**Supplementary Figure 33: PCR validation for spliced ERCC RNA isoform in 12 samples.**

Number above the lanes is the sample id for the 12 samples used in this study. Labels on the right side of the figure indicate transcript id for the RNA isoform being validated and the expected product length from the PCR primers used to amplify the RNA isoform. ERCC spliced refers to the new spliced ERCC RNA isoform found in our data, which was succesfully validated. No splice ERCC refers to the regular ERCC sequence that is no spliced, which was also validated succesfully, as expected.

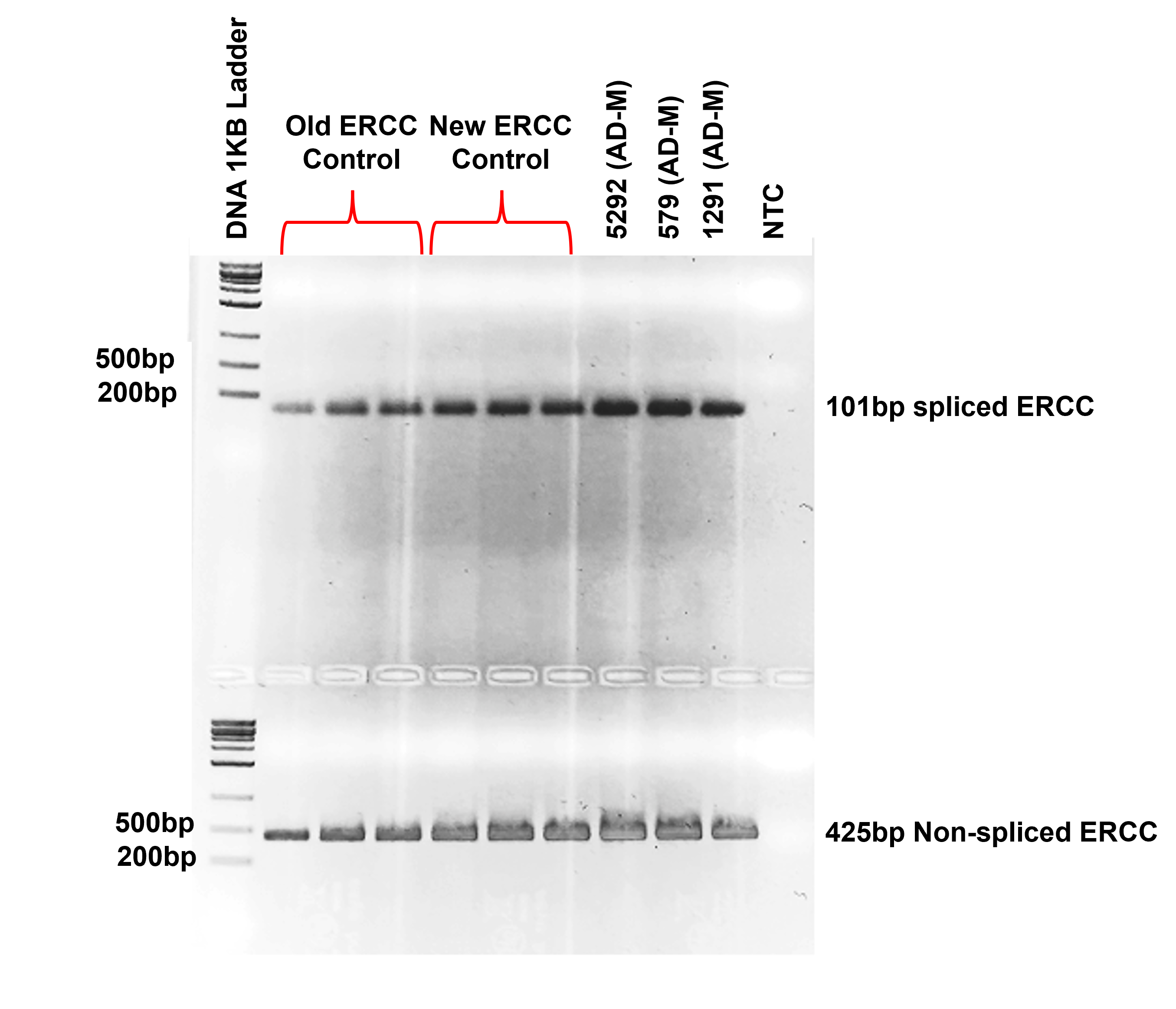

**Supplementary Figure 34: PCR validation for spliced ERCC RNA isoform in two different ERCC batches and 4 samples.**

Number above the lanes is the sample id for 4 samples used in this study, old ERCC control refers to an older ERCC batch whereas new ERCC refers to a fresh ERCC batch we tested with. Labels on the right side of the figure indicate transcript id for the RNA isoform being validated and the expected product length from the PCR primers used to amplify the RNA isoform. ERCC spliced refers to the new spliced ERCC RNA isoform found in our data, which was succesfully validated. No splice ERCC refers to the regular ERCC sequence that is no spliced, which was also validated succesfully as expected.

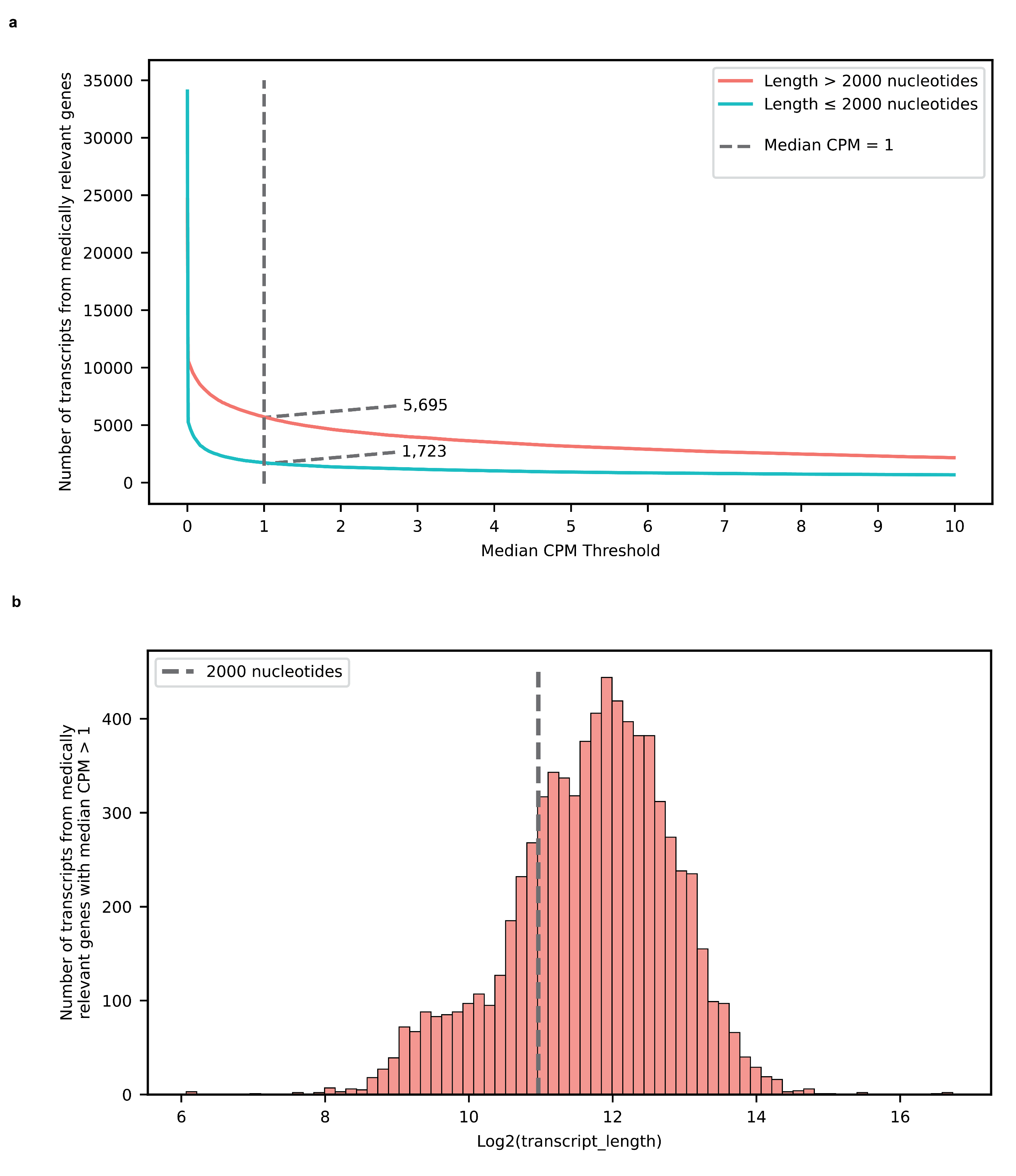

**Supplementary Figure 35: Expression and length distribution for transcripts from medically relevant genes. a,** Lineplot showing number of transcripts from medically relevant genes that are expressed across a range of median CPM thresholds in our data. The red/coral line show transcripts longer than 2,000 nucleotides and the blue/mint line shows trnascripts shorter or equal to 2,000 nucleotides. The dotted line is drawn at median CPM = 1 and the numbers by the line represent the number of transcripts expressed at median CPM > 1 for each transcript length category. **b,** Histogram showing the read length distribution for transcripts from medically relevant genes that our expressed in our data with median CPM > 1. Read length is shown in a log2 scale to avoid stretching the plot because of outliers.

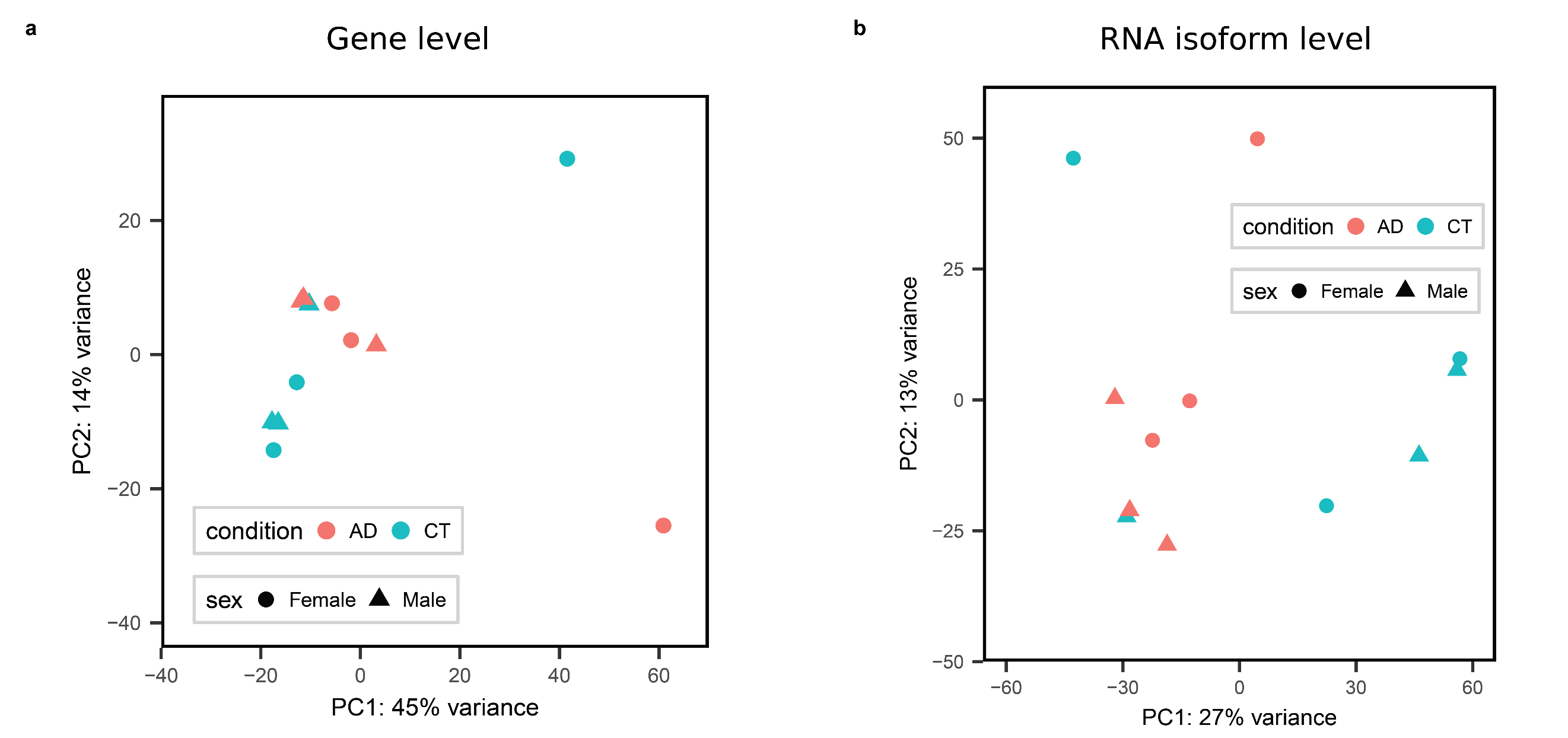

**Supplementary Figure 36: Principal component analysis plots for gene level and RNA isoform level RNAseq data. a,** Principal component analysis plots for gene level data generated using DEseq2 on unfiltered gene level counts. **b,** Principal component analysis plots for RNA isoform level data generated using DEseq2 on unfiltered gene level counts.

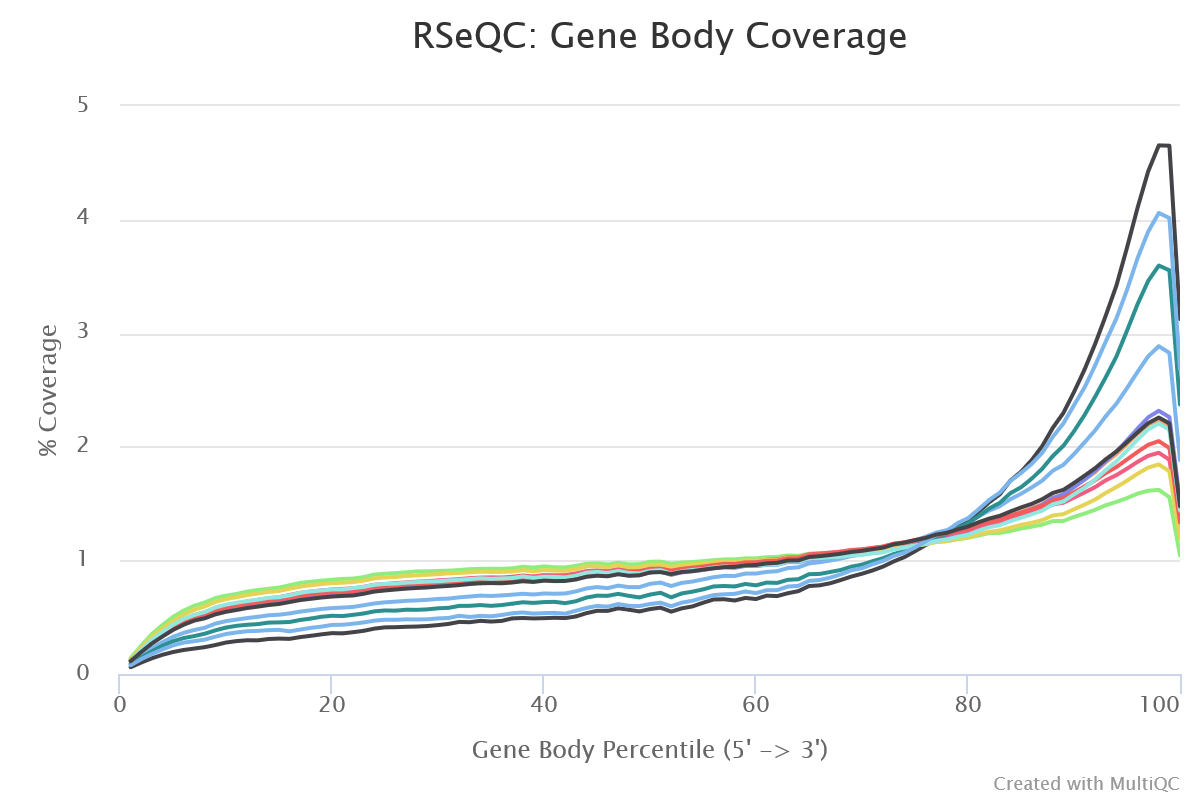

**Supplementary Figure 37: 3’ bias in long-read RNAseq samples from this study.** Y-axis shows percent coverage of housekeeping transcripts and X-axis shows gene body percentile, where 0 is the 5’ end and 100 is the 3’ end. All samples have 3’ bias, but some have a much stronger bias than others. Plot was generated with RseQC^1^ using the “geneBody_coverage.py” module with the housekeeping BED file provided in: <https://sourceforge.net/projects/rseqc/files/BED/Human_Homo_sapiens/hg38.HouseKeepingGenes.bed.gz/download>.

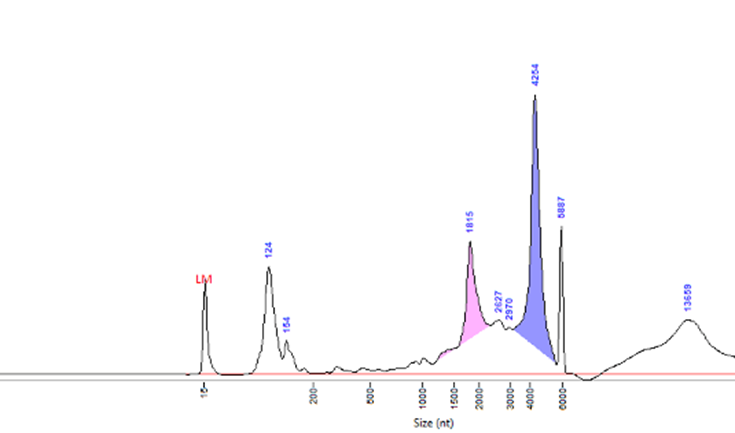

**Supplementary Figure 38.** Fragment Analyzer RNA trace for sample 1163 (Control female).

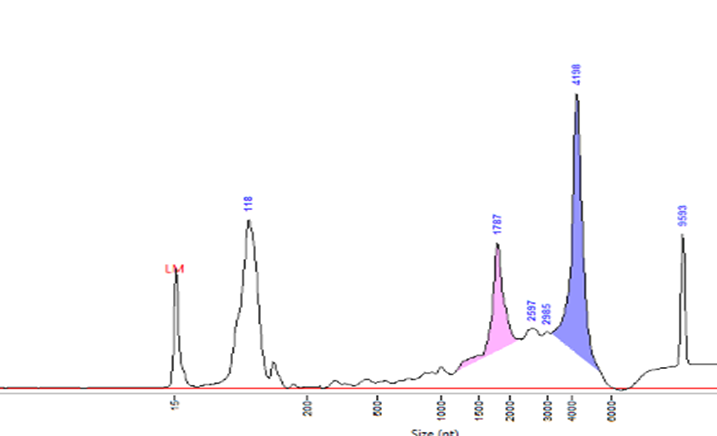

**Supplementary Figure 39.** Fragment Analyzer RNA trace for sample 1131 (Control female).

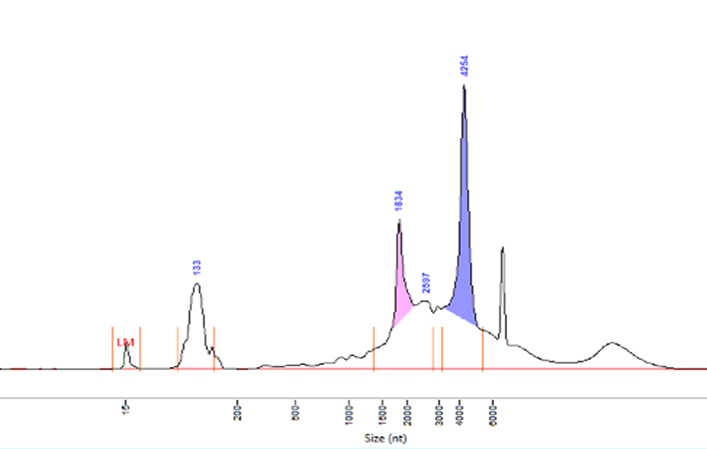

**Supplementary Figure 40.** Fragment Analyzer RNA trace for sample 1092 (Control female).

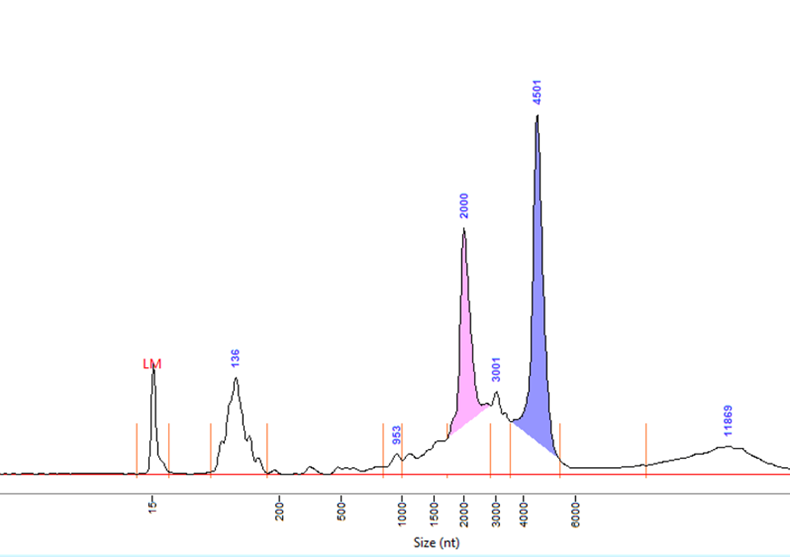

**Supplementary Figure 41.** Fragment Analyzer RNA trace for sample 5356 (Control male).

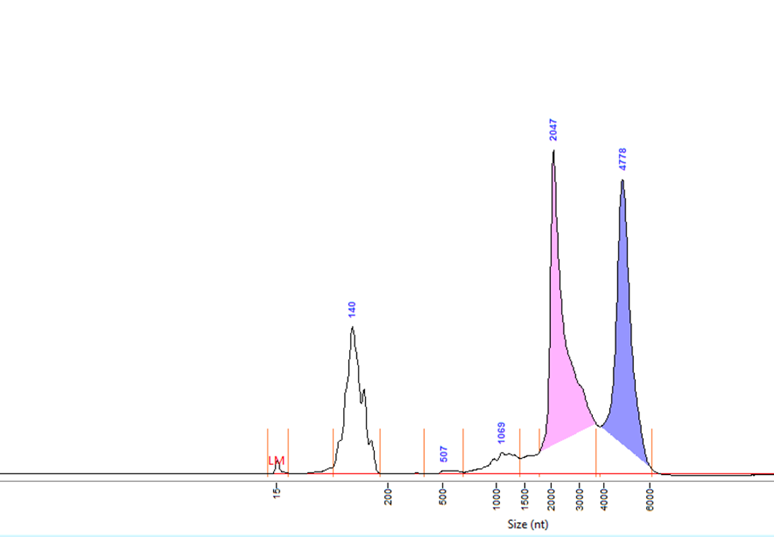

**Supplementary Figure 42.** Fragment Analyzer RNA trace for sample 1304 (Control male).

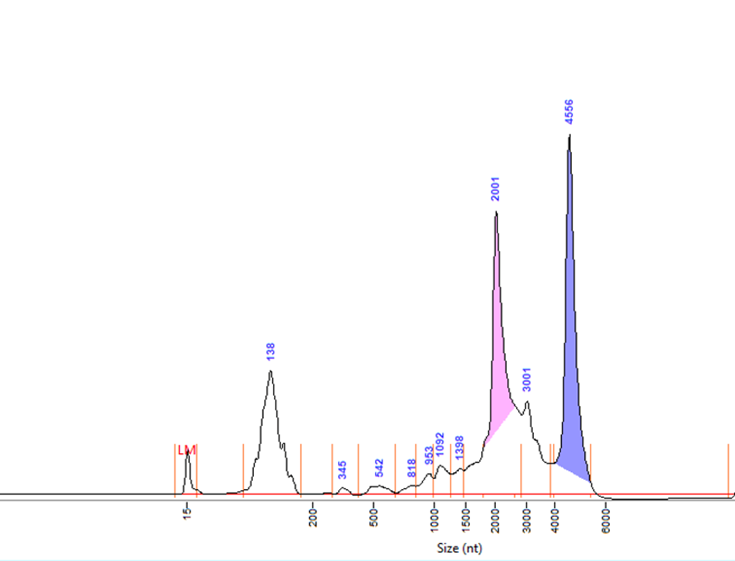

**Supplementary Figure 43.** Fragment Analyzer RNA trace for sample 1271 (Control male).

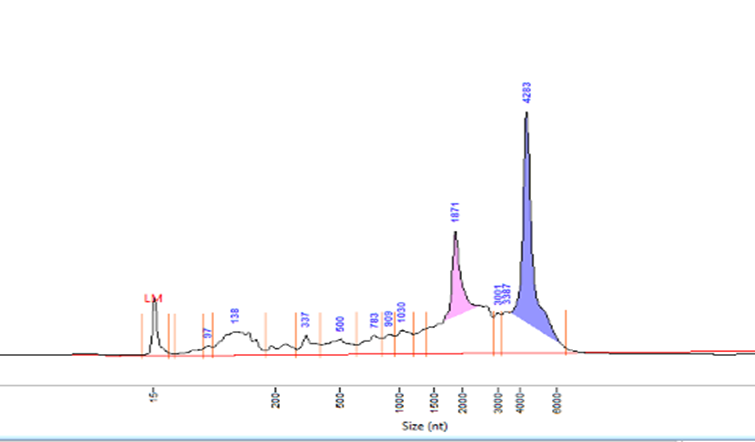

**Supplementary Figure 44.** Fragment Analyzer RNA trace for sample 1186 (Alzheimer’s disease female).

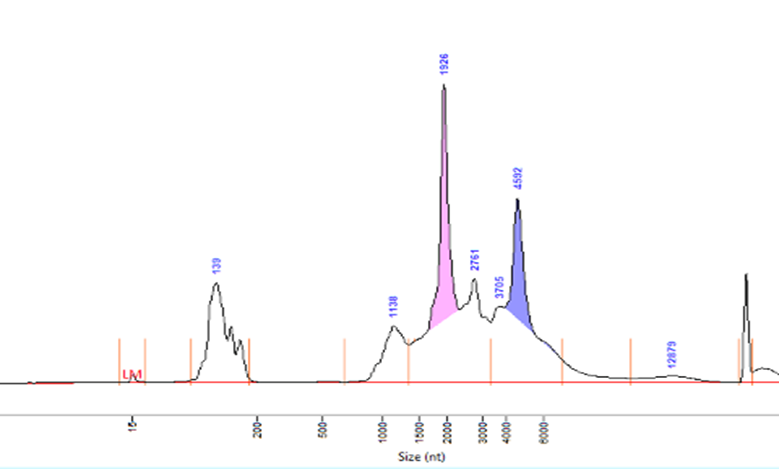

**Supplementary Figure 45.** Fragment Analyzer RNA trace for sample 1218 (Alzheimer’s disease female).

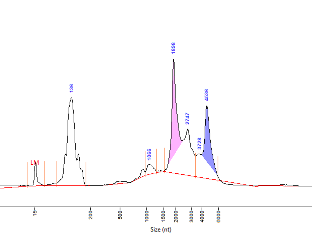

**Supplementary Figure 46.** Fragment Analyzer RNA trace for sample 5295 (Alzheimer’s disease female).

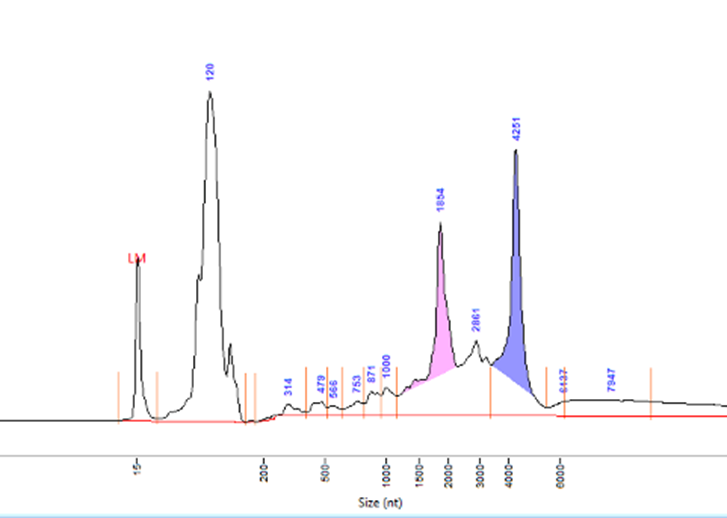

**Supplementary Figure 47.** Fragment Analyzer RNA trace for sample 5292 (Alzheimer’s disease female).

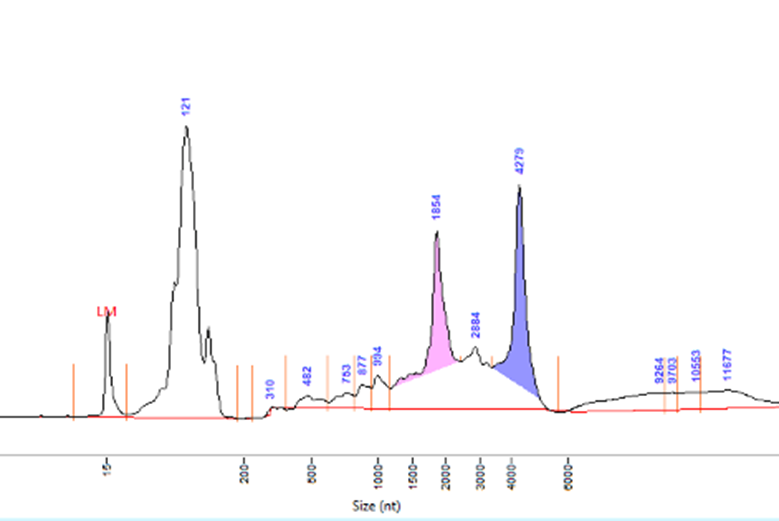

**Supplementary Figure 48.** Fragment Analyzer RNA trace for sample 579 (Alzheimer’s disease female).

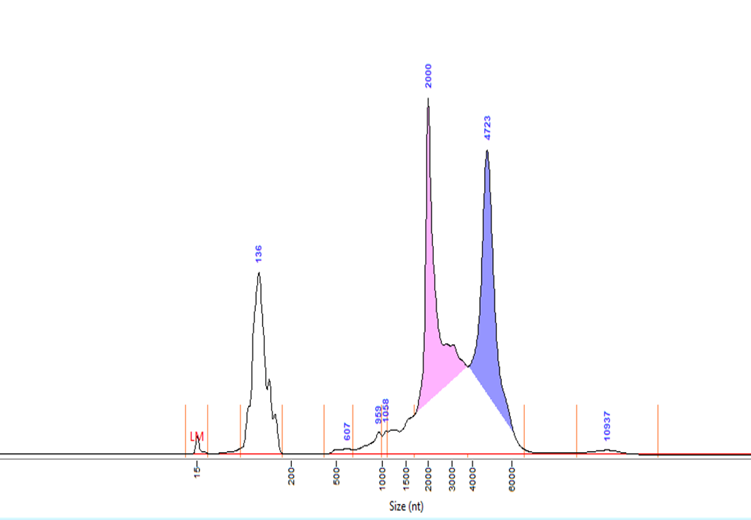

**Supplementary Figure 49.** Fragment Analyzer RNA trace for sample 1291 (Alzheimer’s disease female).

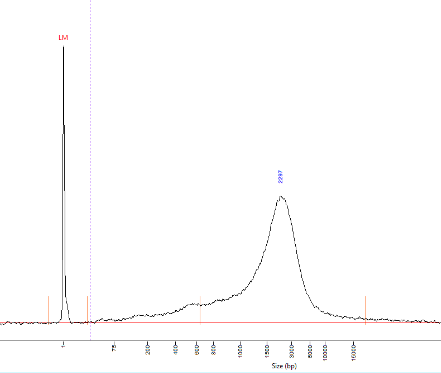

**Supplementary Figure 50.** Fragment Analyzer cDNA trace for sample 1163 (Control female). Peak at 2297 nucleotides.

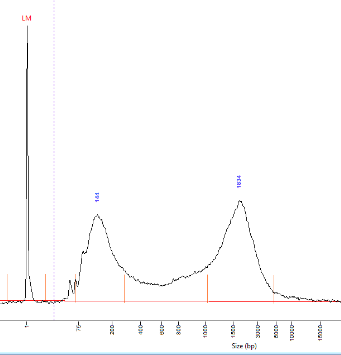

**Supplementary Figure 51.** Fragment Analyzer cDNA trace for sample 1131 (Control female). Lower peak at 144 nucleotides, higher peak at 1834 nucleotides

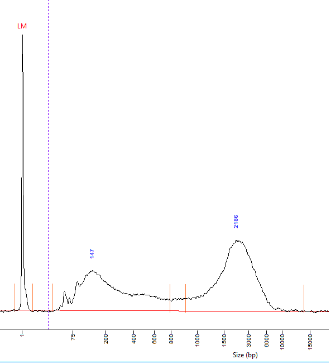

**Supplementary Figure 52.** Fragment Analyzer cDNA trace for sample 1092 (Control female). Lower peak at 147 nucleotides, higher peak at 2186 nucleotides

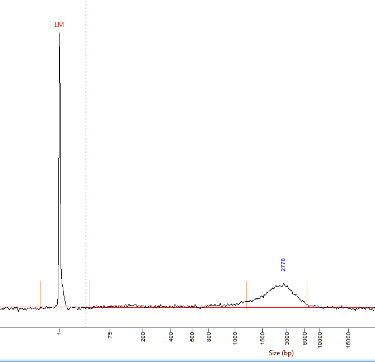

**Supplementary Figure 53.** Fragment Analyzer cDNA trace for sample 5356 (Control male). Peak at 2778 nucleotides.

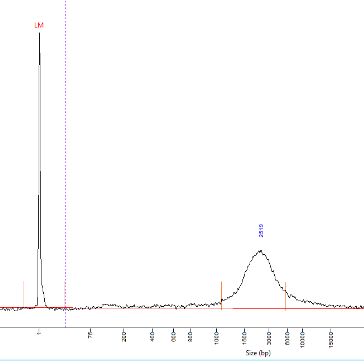

**Supplementary Figure 54.** Fragment Analyzer cDNA trace for sample 1304 (Control male). Peak at 2519 nucleotides.

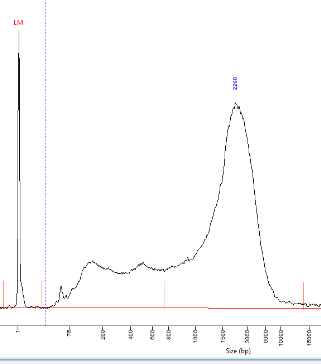

**Supplementary Figure 55.** Fragment Analyzer cDNA trace for sample 1271 (Control male). Peak at 2260 nucleotides.

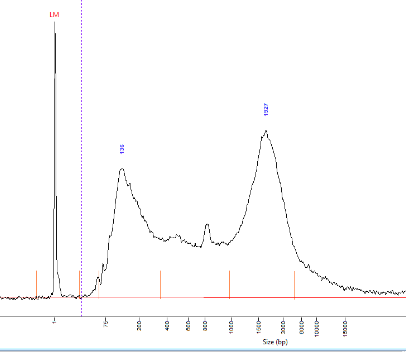

**Supplementary Figure 56.** Fragment Analyzer cDNA trace for sample 1186 (Alzheimer’s disease female). Lower peak at 136 nucleotides, higher peak at 1927 nucleotides.

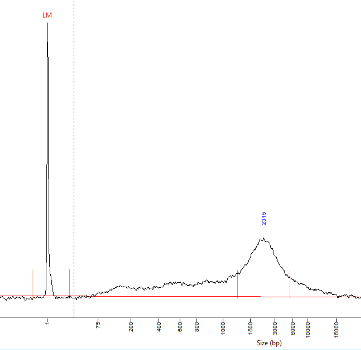

**Supplementary Figure 57.** Fragment Analyzer cDNA trace for sample 1218 (Alzheimer’s disease female). Peak at 2315 nucleotides.

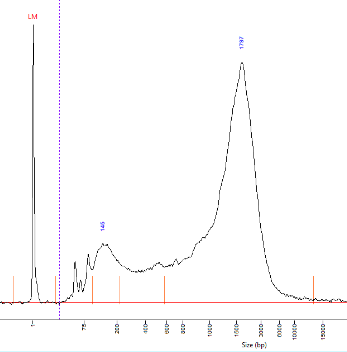

**Supplementary Figure 58.** Fragment Analyzer cDNA trace for sample 5295 (Alzheimer’s disease female). Lower peak at 145 nucleotides, higher peak at 1797 nucleotides.

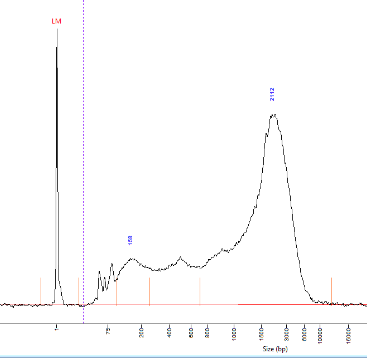

**Supplementary Figure 59.** Fragment Analyzer cDNA trace for sample 5292 (Alzheimer’s disease female). Lower peak at 158 nucleotides, higher peak at 2112 nucleotides.

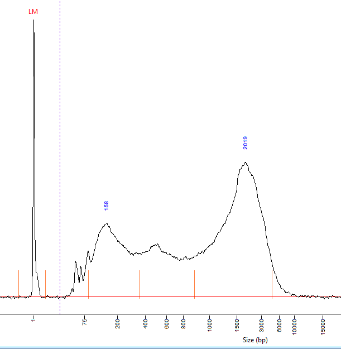

**Supplementary Figure 60.** Fragment Analyzer cDNA trace for sample 579 (Alzheimer’s disease female). Lower peak at 158 nucleotides, higher peak at 2019 nucleotides.

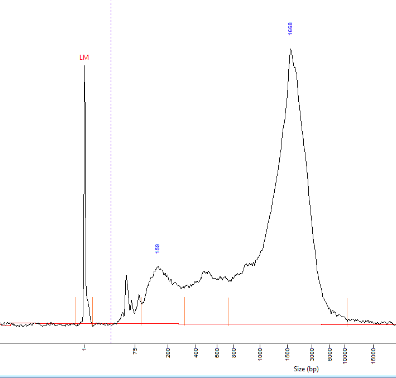

**Supplementary Figure 61.** Fragment Analyzer cDNA trace for sample 1291 (Alzheimer’s disease female). Lower peak at 159 nucleotides, higher peak at 1668 nucleotides.
