## Supplementary File 1 for "Using deep long-read RNAseq in Alzheimer’s disease brain to assess medical relevance of RNA isoform diversity"

LEXOGEN
The RNA Experts

### SPLIT™

Pure RNA for all your experiments

### RNA Extraction Kit
### User Guide

Catalog Number:
008 (SPLIT RNA Extraction Kit)

008UG005V0320

**FOR RESEARCH USE ONLY. NOT INTENDED FOR DIAGNOSTIC OR THERAPEUTIC USE.**
INFORMATION IN THIS DOCUMENT IS SUBJECT TO CHANGE WITHOUT NOTICE.
Lexogen does not assume any responsibility for errors that may appear in this document.

#### PATENTS AND TRADEMARKS

SPLIT is a trademark of Lexogen. Lexogen is a registered trademark (EU, CH, US, CN, AU, NO, BR).
Phase Lock Gel™ is a registered trademark of 5 Prime GmbH, RNA*later*™ and RNase*Zap*™ are registered trademarks of Ambion, Inc., RNasin® is a registered trademark of Promega Corporation. Bioanalyzer®, Fragment Analyzer™, and TapeStation® are trademarks of Agilent Technologies, Inc., DNA-ExitusPlus™ is a trademark of AppliChem GmbH, TRIzol® is a registered trademark of Molecular Research Center, Inc., MACS® is a registered trademark of Miltenyi Biotec B.V. All other brands and names contained in this user guide are the property of their respective owners.

Lexogen does not assume responsibility for violations or patent infringements that may occur with the use of its products.

#### LIABILITY AND LIMITED USE LABEL LICENSE: RESEARCH USE ONLY

This document is proprietary to Lexogen. The SPLIT RNA Extraction Kits are intended for use in research and development only. They need to be handled by qualified and experienced personnel to ensure safety and proper use. Lexogen does not assume liability for any damage caused by the improper use or the failure to read and explicitly follow this user guide. Furthermore, Lexogen does not assume warranty for merchantability or suitability of the product for a particular purpose.

The purchase of the product is subject to Lexogen general terms and conditions (www.lexogen.com/terms-and-conditions/) and does not convey the right to resell, distribute, further sublicense, repackage, or modify the product or any of its components. This document and its contents shall not be used or distributed for any other purpose and/or otherwise communicated, disclosed, or reproduced in any way without the prior written consent of Lexogen.

For information on purchasing additional rights or a license for use other than research, please contact Lexogen.

#### WARRANTY

Lexogen is committed to providing excellent products. Lexogen warrants that the product performs to the standards described in this user guide until the expiration date. Should this product fail to meet these standards due to any reason other than misuse, improper handling, or storage, Lexogen will replace the product free of charge or issue a credit for the purchase price. Lexogen does not provide any warranty if product components are replaced with substitutes.

Under no circumstances shall the liability of this warranty exceed the purchase price of this product.

#### LITERATURE CITATION

When describing a procedure for publication using this product, please refer to it as the SPLIT RNA Extraction Kit.

We reserve the right to change, alter, or modify any product without notice to enhance its performance.

---

#### CONTACT INFORMATION

**Lexogen GmbH**
Campus Vienna Biocenter 5
1030 Vienna, Austria
www.lexogen.com


**Support**


### 1. Overview

The SPLIT RNA Extraction Kit enables fast and highly efficient extraction of high quality, high purity RNA from a broad range of biological samples including cell culture, animal and plant tissue, and fluid samples. Thus the obtained RNA is ideal for seamless library preparation for Next Generation Sequencing and other demanding applications such as full-length reverse transcription, sample preparation for microarray analysis, or RT-qPCR. Furthermore, SPLIT recovers the complete RNA size ranges, including small RNAs (≥17 nt).

First, the sample is homogenized in a highly chaotropic isolation buffer which facilitates effortless and complete solubilization, and guarantees complete RNase inhibition.
Acidic buffer and acidic phenol are added to create a monophasic solution, a step that is essential for the efficient separation of genomic DNA into the organic phase. Chloroform is added, which creates a biphasic system, and phases are cleanly separated using Phase Lock Gel tubes. The use of these tubes mitigates the risk of contaminating the upper aqueous phase that contains RNA with the lower phenol phase that contains DNA and proteins.
The aqueous phase is mixed with isopropanol and the RNA is precipitated onto a silica column. Contaminants are efficiently cleared away by several washes and the eluted RNA is ready for any downstream application.

Please note that acidic phenol, chloroform, ethanol, and isopropanol have to be supplied by the user.

Protocols are given for RNA extraction from human cell culture, animal, and plant tissue, as well as fluid samples. Optionally, a modified protocol allows purification of small and large RNA-enriched fractions (see Appendix D, p.21).

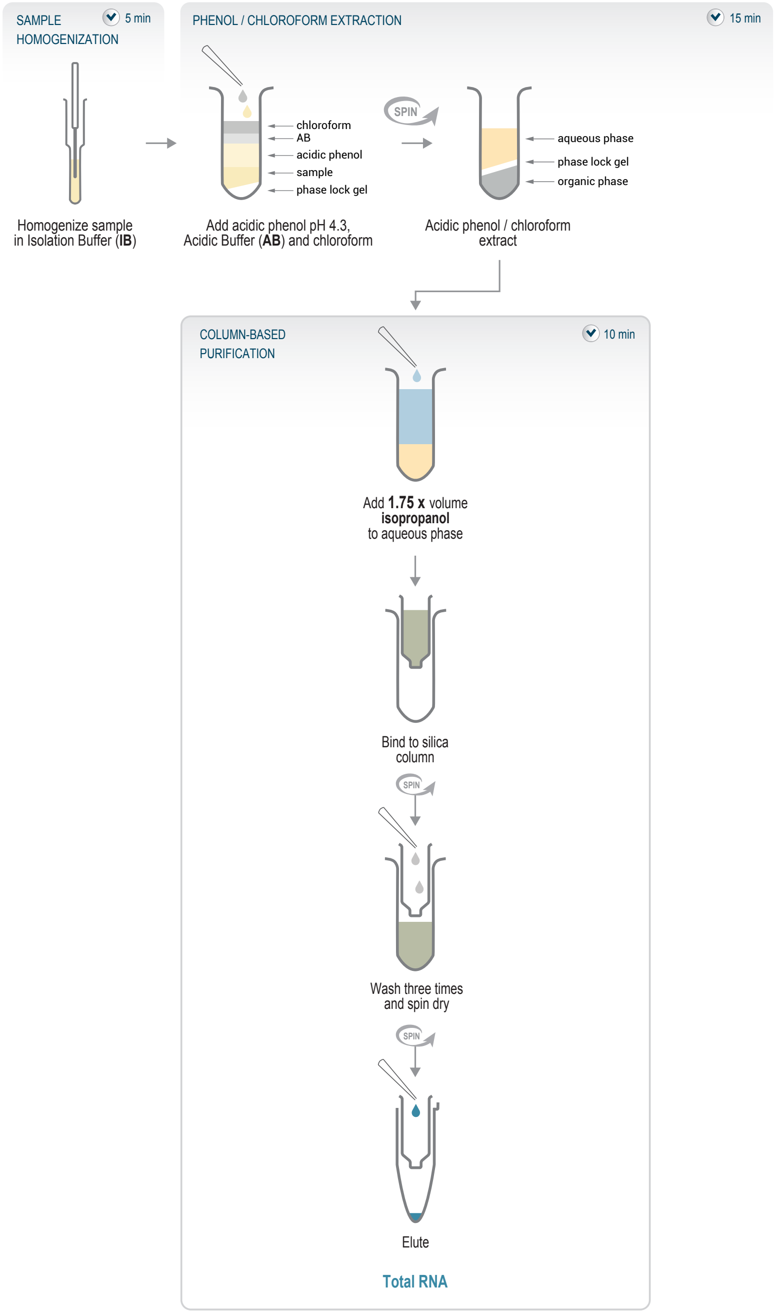

Figure 1. Schematic overview of the SPLIT workflow.

### 2. Kit Components and Storage Conditions

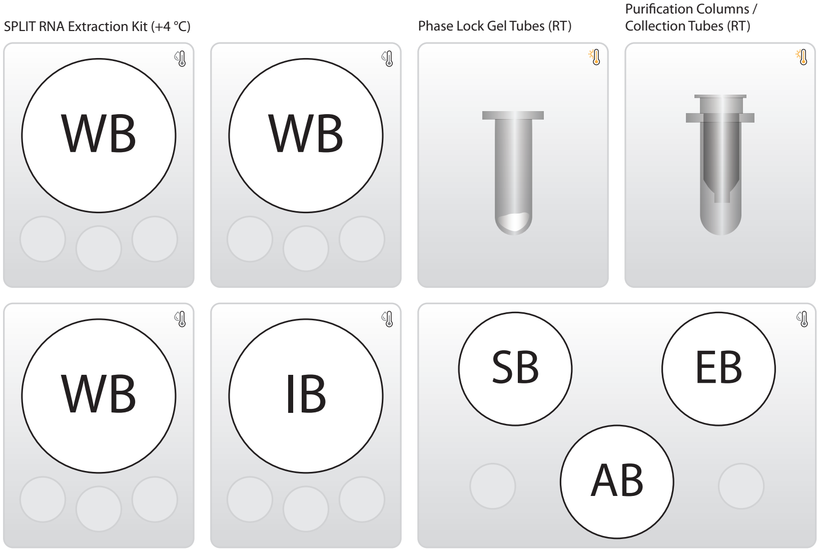

Figure 2. Location of kit components.

| Kit Component | Label | Volume | Storage |
|---|---|---|---|
| **Isolation Buffer** | **IB** | 21.1 ml [*] | +4 °C |
| **Acidic Buffer** | **AB** | 7.9 ml [*] | +4 °C |
| **Wash Buffer** | **WB** | 3 x 6.5 ml [1,*] | +4 °C |
| **Elution Buffer** | **EB** | 2.6 ml [2,*] | +4 °C |
| **Storage Buffer** | **SB** | 2.6 ml [2,*] | +4 °C |
| **Phase Lock Gel tubes** | **Phase Lock Gel tubes** | 48 | RT |
| **Purification columns** | **Purification columns** | 48 | RT |
| **Collection Tubes** | **Collection Tubes** | 48 | RT |

*including ≥10 % surplus

[1] Excluding ethanol (to be added by the user - see bottle for volume to add).
[2] For each RNA fraction, either EB or SB is required.

Upon receiving the SPLIT (Cat. No. 008) kit, store it at +2 to +8 °C.
**ATTENTION:** Phase Lock Gel tubes must not be frozen.

The Isolation Buffer (**IB**) contains guanidine isothiocyanate, an irritant, which upon protocol completion is also present in flow-through and wash fractions. This chemical is harmful when in contact with the skin, inhaled, or ingested. Do not add bleach or acidic solutions directly to solutions or sample preparation waste that contains guanidine isothiocyanate, as reactive compounds and toxic gases are formed.

**IB** is to be used at +4 °C. All other components should be equilibrated to room temperature before use.

Check the contents of **IB**, **AB**, **WB**, and **SB** which may precipitate during shipping and storage. If a white precipitate is visible, incubate at 37 °C until buffer components dissolve completely.

Add 25 ml absolute ethanol to each of the three bottles with Wash Buffer (**WB**) concentrate and shake to combine.

### 3. User-Supplied Consumables and Equipment

Check to ensure that you have all of the necessary materials and equipment before beginning with the RNA extraction. All reagents, equipment, and labware must be free of nucleases and nucleic acid contamination.

**ATTENTION:** Before starting this protocol, please read the General Guidelines for Lexogen Kits, which are available online. These provide a detailed overview of RNA and kit component handling, as well as general RNA input requirements.

#### Reagents

| Reagent | Volume | Comment |
|---|---|---|
| **Phenol solution pH 4.3** | 19.2 ml | e.g., Sigma-Aldrich (P4682-100ML) or VWR (0981-400ML) |
| **Chloroform** | 9.6 ml | |
| **Isopropanol** | ~ 50.4 ml | 2-Propanol |
| **Ethanol abs.** | 3 x 25 ml | Added to **WB** |

When working with the phenol solution and with chloroform, always work in a fume hood. Phenol is toxic and corrosive, and it should not come in contact with skin, eyes, or the respiratory tract and may cause chemical burns to the exposed area.

#### Equipment

- Fume hood for organic solvent handling.
- Refrigerated benchtop centrifuge (12,000 x g, rotor compatible with 1.5 ml and 2.0 ml tubes).
- Calibrated single-channel pipettes for handling 10 µl to 1,000 µl volumes.
- Vortex mixer.
- UV-spectrophotometer to quantify RNA.

#### Labware

- Suitable pipette tips (pipette tips with aerosol barriers recommended).
- 1.5 ml and 2.0 ml tubes with cap, low binding, certified ribonuclease-free.
- Benchtop cooler or ice pellets in ice box (for short-term storage of RNA).

#### Optional Equipment & Solutions

- Tissue grinder (hand-held homogenizer).
- Liquid nitrogen (for RNA extraction of plant tissue).
- Automated microfluidic electrophoresis station (Agilent Technologies 2100 Bioanalyzer).
- Agarose gels, dyes, and electrophoresis rig (for RNA quality control).
- DNA-ExitusPlus™ (AppliChem GmbH).
- RNaseZap.
- RNase inhibitor.

The complete set of materials, reagents, and labware for quality control is not listed.

### 4. Detailed Protocol

#### 4.1. Sample Homogenization

##### 4.1.1. Animal Tissue

##### Preparation

| Tissue | Weigh and Reduce Tissue | Homogenization |
|---|---|---|
| **Animal tissue** – freshly harvested or frozen or thawed at +4 °C if stored in RNAlater | **Tweezers** – sterile<br>**Scalpel** – sterile<br>**Gauze pad** – sterile | **Isolation Buffer** (IB) - at +4 °C or on ice |
| **Fume hood** or laminar-flow cabinet | **Precision balance** | **Tissue grinder** |

##### Homogenization

Tissue is homogenized in a highly chaotropic solution. This protocol is specific for hand-held tissue grinders (glass homogenizers with pestle) but can be easily adapted for other homogenization protocols. Optimally, the tissue should be stored at -20 °C in RNAlater (Ambion, Inc.). Tissue frozen without preservation must not be thawed before homogenization to maintain RNA integrity.

To prevent cross contamination, it is best to work in a fume hood or a laminar-flow cabinet that can be UV-irradiated.

1. Add 400 µl cold (+4 °C) Isolation Buffer (**IB**) into a glass tissue grinder.
   **NOTE:** Isolation Buffer (**IB**) contains detergent. Pipette carefully to avoid foaming.
2. Use sterile tweezers to transfer a tissue piece onto a fresh, sterile gauze pad. If RNAlater was used for conservation, dry the tissue by tapping onto the gauze pad.
3. Determine the weight of the tissue on a precision balance. The protocol is efficient for extraction of up to 10 µg of total RNA. See Appendix A, p.17 for details on input and extraction efficiency.
4. **OPTIONAL:** Hard to homogenize tissues such as tendons or cartilage can be reduced using a scalpel to facilitate solubilization in the next steps. Also, short incubation of tissue with Isolation Buffer (**IB**) prior to homogenization can help solubilization.
5. Using tweezers, transfer the tissue pieces quantitatively into the Isolation Buffer (**IB**) in the tissue grinder.

6 Homogenize the tissue by carefully moving the pestle up and down. Simultaneous rotation helps to dissolve also larger pieces. Do not pull out the pestle completely to avoid foaming. The tissue is usually homogenized within 2 - 3 minutes; avoid extended homogenization and warming up of Isolation Buffer (**IB**).

7 Continue immediately with the phenol / chloroform extraction at step 8 in 4.2., p.13

After use, clean the tissue grinder thoroughly with a detergent such as DNA-ExitusPlus™ (AppliChem GmbH), then rinse thoroughly with ultra-filtered water and finally with 75 % ethanol.

#### 4.1.2. Plant Tissue

##### Preparation

| Tissue | Weigh and Reduce Tissue | Homogenization |
|---|---|---|
| **Plant tissue –** freshly harvested or frozen at -80 °C or -20 °C in RNAlater or already ground and frozen in **IB** at -20 °C | **Tweezers** – sterile<br>**Scalpel** – sterile | **Liquid nitrogen**<br>**Isolation Buffer (IB)** - at +4 °C or on ice |
| **Fume hood** or laminar-flow cabinet | **Precision balance** | **Pestle and mortar** |

##### Homogenization

Plant material is disrupted whilst frozen (i.e., in liquid nitrogen, or over dry ice) and homogenized in a highly chaotropic solution. Optimally, the plant material should be extracted immediately after harvesting. If storage of plant material is required, flash-freeze the sample in liquid nitrogen and store at -80 °C, or at -20 °C in RNAlater. Already ground plant tissue can also be stored in Isolation Buffer (**IB**) at -20 °C. To prevent cross contamination, it is best to work in a fume hood or a laminar-flow cabinet that can be UV-irradiated.

**NOTE:** Disruption of plant material can also be done using other devices such as ball mills and homogenization methods may need to be optimized for different types of input plant material (e.g., waxy, low water content tissues). For more information please contact.

1 Determine the weight of the tissue on a precision balance, working under sterile conditions (e.g., use sterile tweezers for transfer). The protocol is efficient for extraction of up to 10 µg of total RNA. See Appendix A, p.17 for details on input and extraction efficiency.

2 Quickly cut the plant tissue into small pieces using a scalpel and freeze in liquid nitrogen.

3 Grind the tissue using pestle and mortar. This can be done in liquid nitrogen, or by grinding the frozen tissue in a 1.5 ml or 2 ml tube over dry ice.

4. Allow any liquid nitrogen to evaporate.

5. Resuspend the tissue in 400 µl cold (+4 °C) Isolation Buffer (**IB**). Make sure to completely cover the tissue with **IB**.
**NOTE:** Isolation Buffer (**IB**) contains detergent. Pipette carefully to avoid foaming.

6. **OPTIONAL:** Further homogenize the sample by carefully moving the pestle up and down. Do not pull out the pestle completely to avoid foaming. Avoid extended homogenization and warming up of Isolation Buffer (**IB**).

7. Continue immediately with the phenol / chloroform extraction at step 8 in 4.2., p.13. **OPTIONAL:** ☞ Safe stopping point. Ground plant tissue in Isolation Buffer (**IB**) can be stored at -20 °C at this point.

After use, clean the pestle and mortar thoroughly with a detergent such as DNA-ExitusPlus™ (AppliChem GmbH), then rinse thoroughly with ultra-filtered water and finally with 75 % ethanol.

#### 4.1.3. Cultured Cells

##### Preparation

| Cells | Solubilization |
|---|---|
| **Cells** – freshly harvested, FACS / MACS sorted or frozen | **Isolation Buffer (IB)** – at +4 °C or on ice |
| **Fume hood** or laminar-flow cabinet | |

##### Solubilization

Cells are solubilized in a highly chaotropic solution. Lyse the cells fully in the Isolation Buffer (**IB**) and proceed directly to RNA extraction. Cell lysates in **IB** can also be stored at -80 °C prior to extraction.
**NOTE:** Recommendations for freshly harvested and FACS / MACS sorted cells: spin down to pellet the cells and wash twice with Phosphate Buffered Saline (1x PBS). Remove PBS completely before adding IB. To prevent cross contamination, it is best to work in a fume hood or a laminar-flow cabinet that can be UV-irradiated.

**ATTENTION:** Do not store cells as cell pellets in IB without resuspending and lysing the cells beforehand. Cell lysis is required in order to maintain RNA integrity throughout storage in IB. Alternatively, snap freeze and store cell pellets after completely removing PBS and add IB when thawing to lyse cells before starting RNA extraction.

1. Harvest, pellet, and wash the cells. The protocol is suitable for extraction of e.g., $10^6$ cells of a human suspension cell culture. SPLIT RNA extraction has also been successfully performed with 100 cells input.

2 Add 400 µl cold (+4 °C) Isolation Buffer (**IB**) to the cells.
**NOTE:** Isolation Buffer (**IB**) contains detergent. Pipette carefully to avoid foaming.

3 Lyse the cells by carefully pipetting up and down. Cells are usually lysed within 1 - 2 minutes.

4 Continue immediately with the phenol / chloroform extraction at step 8 in 4.2., p.13.

#### 4.1.4. Fluid Samples

##### Preparation

| Fluid samples | Solubilization |
|---|---|
| **e.g., plasma** – freshly harvested | **Isolation Buffer (IB)** – at +4 °C or on ice |
| **Centrifuge** – at +4 °C<br>**Fume hood** or laminar-flow cabinet | |

##### Solubilization

The solubilization / homogenization step of the SPLIT protocol can be applied to a whole range of cells in fluids (aspirates, viral supernatants, plasma, urine etc.). Depending on the sample a homogenization step might be necessary or you can proceed directly to the phenol / chloroform extraction in 4.2., p.13, step 8. To prevent cross contamination, it is best to work in a fume hood or a laminar-flow cabinet that can be UV-irradiated.

**NOTE:** For sample types other than plasma, prior centrifugation may or may not be required. Use up to 200 µl of the liquid sample as input, then add 200 µl of Isolation Buffer (IB). For RNA extraction from whole blood, use up to 50 µl of a blood sample as an input.

**ATTENTION:** If the liquid sample volume is < 200 µl, add extra IB to bring the total volume of the sample / IB mix to 400 µl.

1 Centrifuge 300 - 400 µl of plasma at 12,000 x g for 5 minutes at 4 °C to pellet the cell debris.

2 Transfer 200 µl of the supernatant to a new tube. Take care to avoid carry-over of cell debris.

3 Add 200 µl Isolation Buffer (**IB**) and mix properly.
**NOTE:** Isolation Buffer (**IB**) contains detergent. Pipette carefully to avoid foaming.

4 Continue immediately with the phenol / chloroform extraction at step 8 in 4.2., p.13.

#### 4.2. Phenol / Chloroform Extraction

##### Preparation

| | For each sample | Temperature |
|---|---|---|
| **Phenol solution pH 4.3**[1]<br>**Acidic buffer (AB)**<br>**Chloroform**[1] | 400 µl<br>150 µl<br>200 µl | +4 °C<br>RT<br>RT |
| **Phase Lock Gel tube**<br>**2 ml tube** | 1<br>1 | RT<br>RT |
| **Centrifuge** – at 18 °C<br>**Fume hood**<br>**Vortex mixer** | | +18 °C |

[1] **Caution:** When working with phenol or chloroform always use a fume hood and discard waste according to applicable Health and Safety regulations.

##### Phenol / Chloroform Extraction

Utilizing a highly specific phenol / chloroform extraction, RNA is partitioned into the upper, aqueous phase whereas DNA and proteins are partitioned into the lower, organic phase. The Phase Lock Gel matrix will act as a barrier in between the two phases.

8 For each sample, centrifuge one Phase Lock Gel tube for 1 minute at 12,000 x g at +18 °C. This collects the gel on the bottom of the tube. **ATTENTION:** Phase Lock Gel tubes should be equilibrated for 30 minutes at room temperature before use!

9 Transfer the homogenized sample in Isolation Buffer (**IB**) into a Phase Lock Gel tube.

10 Add 400 µl phenol solution pH 4.3 and mix by inverting the tube 5 times. **NOTE:** The phenol solution should be used at its storage temperature of +4 °C.

11 Add 150 µl Acidic Buffer (**AB**) and mix by pipetting.

12 Add 200 µl of chloroform.

13 Mix thoroughly by repeatedly inverting the tubes for 15 seconds (do not vortex!). **ATTENTION:** Thorough mixing is essential to disperse the chloroform efficiently and effectively separate all the phenol that will contain gDNA and protein into the organic and interphase.

14 Incubate for 2 minutes at room temperature.

15 Centrifuge for 2 minutes at 12,000 x g at +18 °C. **ATTENTION:** Temperatures below +18 °C can negatively influence phase separation. Repeat centrifugation at +18 °C if phase separation is incomplete.

16 Transfer the upper phase to a new 2 ml tube by decanting. **ATTENTION:** Do not transfer the upper phase by pipetting to avoid carry-over of the Phase Lock Gel.

17 For the purification of total RNA, proceed with **step 18 in 4.3.1.** For the purification of the large and small RNA-enriched fraction, proceed with **step 18 in Appendix D, p.21.**

#### 4.3. Column-based Purification

##### Preparation

| | Total RNA | Temperature |
|---|---|---|
| **Isopropanol**<br>**Wash Buffer (WB)**[1]<br>**Elution Buffer (EB) or**<br>**Storage Buffer (SB)**[2] | ~1,050 µl<br>1,500 µl<br>50 µl | RT<br>RT<br>RT |
| **Purification column**<br>**Collection tube**<br>**1.5 ml tube** | 1<br>1<br>1 | RT<br>RT<br>RT |
| **Centrifuge**<br>**Vortex mixer** | | +18 °C |

[1] **Caution:** Discard waste containing guanidine isothiocyanate, phenol, and chloroform according to applicable Health and Safety regulations.

[2] See Appendix C, p.20 whether **EB** or **SB** should be used for elution.

**NOTE:** Repeat centrifugation or increase centrifugation time if sample did not pass filter completely.

##### 4.3.1. Column Loading of Total RNA

The total RNA is precipitated onto a silica column by addition of 1.75x volume of isopropanol.

**NOTE:** To isolate the large and small RNA-enriched fractions (cutoff at ~ 150 nt), please see Appendix D, p.21.

18 Determine the volume of the aqueous phase, which may vary, depending on the sample volume and volume transfer efficiency during homogenization and extraction. Add isopropanol at 1.75x of this volume. Mix by vortexing for 10 seconds.
**EXAMPLE:** Add 1,050 µl isopropanol to 600 µl sample.

19 Place a purification column in a collection tube.

20 Apply a maximum of 800 µl of the mixture from step 18 (aqueous phase with isopropanol) to the column.

21 Centrifuge for 20 seconds at 12,000 x g at +18 °C and discard the contents of the collection tube.

22 Repeat steps 20 - 21 until the mixture is loaded completely then proceed to column washing and elution at step 23 in 4.3.2., p.15.

#### 4.3.2. Column Washing and Elution of RNA

##### Column Washing

The RNA is further purified by washing on the column.

| | |
|---|---|
| 23 | Apply 500 µl of Wash Buffer (**WB**) to the column and centrifuge for 20 seconds at 12,000 x g at +18 °C. Empty the collection tube. Repeat this step twice for a total of three washes. |
| 24 | Centrifuge for 1 minute at 12,000 x g at +18 °C.<br>**ATTENTION:** This step is essential to remove all traces of ethanol. |
| 25 | Discard the collection tube and place the purification column in new 1.5 ml tube. |
| 26 | Make sure that no ethanol traces are carried to the new tube. |

##### Elution of RNA

The RNA is eluted into an elution or storage buffer.

| | |
|---|---|
| 27 | Pre-warm the Elution Buffer (**EB**) or Storage Buffer (**SB**) for 5 minutes at 70 °C. |
| 28 | Add 10 - 50 µl of the pre-warmed Elution Buffer (**EB**) or Storage Buffer (**SB**) to the column and incubate for 1 minute at room temperature. |
| 29 | Centrifuge for 1 minute at 12,000 x g at +18 °C. |
| 30 | At this point the total RNA is purified and ready for quality control (Appendix B, p.18) and downstream applications. |
| 31 | **OPTIONAL:** Add RNase inhibitor (not included). See Appendix C, p.20 for RNA storage. Note that the RNase inhibitor might absorb at 230 nm, therefore use buffer with RNase inhibitor added as blank for OD measurements. |

### 5. Short Procedure

#### Extraction of Total RNA

**ATTENTION:** All centrifugation steps are at 12,000 x g and +18 °C!

##### 20 min Homogenization and Phenol / Chloroform Extraction

| | |
|---|---|
| | **Homogenization** |
| ☐ | Homogenize sample in 400 µl **IB**. |
| | **Phenol / Chloroform Extraction** |
| ☐ | Centrifuge 1 Phase Lock Gel tube for 1 min. |
| ☐ | Transfer homogenate into a Phase Lock Gel tube. |
| ☐ | Add 400 µl phenol solution pH 4.3, mix by inverting the tube 5 times. |
| ☐ | Add 150 µl **AB**, mix by pipetting. |
| ☐ | Add 200 µl chloroform and mix by repeatedly inverting the tube for 15 sec. |
| | **ATTENTION:** Do not vortex! |
| ☐ | Incubate for 2 min at RT. |
| ☐ | Centrifuge for 2 min (or longer if phase separation is incomplete). |
| ☐ | Decant the upper phase into a 2 ml tube. |
| | **ATTENTION:** Do not transfer the upper phase by pipetting! |

##### 10 min Purification of Total RNA

| | |
|---|---|
| | **Column Loading Total RNA** |
| ☐ | Measure volume of transferred upper phase. |
| ☐ | Add 1.75x vol. isopropanol to the upper phase. |
| ☐ | Mix by vortexing for 10 sec. |
| ☐☐ | Load max. 800 µl onto purification column in collection tube. Centrifuge for 20 sec and discard flow-through. Repeat until mixture is loaded completely. |
| | **Column Washing** |
| ☐☐☐ | Apply 500 µl **WB** and centrifuge for 20 sec. Empty collection tube. Repeat this step twice for a total of three washes. |
| ☐ | Centrifuge for 1 min to spin dry column. |
| | **Elution** |
| ☐ | Place purification column in a 1.5 ml tube. |
| ☐ | Pre-warm **EB** or **SB** for 5 min at 70 °C. |
| ☐ | Apply 10 - 50 µl of pre-warmed **EB** or **SB**, incubate for 1 min at RT. |
| ☐ | Centrifuge for 1 min. |
| ☐ | **OPTIONAL:** Add RNase inhibitor (not included). |

**NOTE:** Repeat the centrifugation or increase centrifugation time if sample did not fully pass through the filter completely.

### 6. Appendix A: Sample Input and Extraction Efficiencies

If immediate RNA extraction is not possible, tissue samples can be either flash-frozen with liquid nitrogen and stored at -80 °C, or preserved in RNAlater (Ambion, Inc.) and stored at -20 °C or -80 °C. Tissues / cells frozen without RNAlater preservation must only be thawed during the homogenization step in cold Isolation Buffer (+4 °C) to keep RNases inactive.

RNA extraction efficiency for mouse liver is typically 4.0 - 4.5 µg total RNA / mg tissue (3.0 - 3.5 µg large RNA and 0.6 µg small RNA / mg tissue). The maximum binding capacity of the purification column is 10 µg RNA, which should not be exceeded for optimal results. For mouse liver tissue, this translates into an upper limit of 2.5 mg input per extraction. Other tissues have different RNA content, and the input might have to be adjusted accordingly.

The SPLIT RNA Extraction Kit has been used for isolation of RNA from different organisms including animal (e.g., mouse, human) and plant tissues (e.g., *A. thaliana*, *Picea abies*), insects (e.g., drosophila), cell lines (e.g., human), fluid samples (e.g., plasma), and others (jellyfish, fungi, bacteria). Please contact for information on protocol adaptations for other sample types.

### 7. Appendix B: RNA Quality Control

#### RNA Integrity

The integrity of an RNA sample can be assessed with a variety of methods (see Fig. 3). We recommend the use of a microfluidics assay such as the RNA6000 series for the 2100 Bioanalyzer (Agilent Technologies Inc.). However, RNA quality can also be assessed with denaturing agarose gel electrophoresis if such a device is not available. Most microfluidics platforms will carry out an automated peak analysis and generate a quality score (RIN or RQN) in addition to the 28S / 18S rRNA ratio. The quality of RNA extracted with the SPLIT RNA Extraction Kit almost exclusively depends on the extraction source: a RIN of 10 and a 28S / 18S rRNA ratio of 2.7 can be obtained from human cell culture homogenized according to 4.1.3. Extractions from tissue samples usually result in RNA with a RIN of 8.0 - 9.5.

#### Potential Contaminants

RNA samples should be free of salts, metal ions, and organic solvents, which can be carried over from the RNA extraction. Several sources of contamination can be detected with a UV-Vis spectrophotometer. An acceptably pure RNA sample should have an A260 / A280 ratio between 1.8 and 2.1. The A260 / A230 ratio should also be greater than 1.8. Several common contaminants including proteins, chaotropic salts and phenol absorb strongly between 220 and 230 nm and can often be identified as peaks in this region. Contamination with any of these generates a lower A260 / A230 ratio. Phenol also has an absorption maximum between 250 and 280 nm, which overlaps that of nucleic acid, so high 230 nm absorbance combined with a biphasic or broad peak between 250 and 280 nm may indicate contamination with phenol rather than chaotropic salts.

#### Genomic DNA Contamination

The SPLIT RNA Extraction Kit was designed for minimizing the genomic DNA (gDNA) content in the RNA sample. gDNA is indistinguishable from RNA on a spectrophotometer, and many of the dyes used in RNA microfluidics assays stain single-stranded nucleic acids much more intensely than double-stranded. Hence, low to moderate amounts of gDNA may not be readily visible with an RNA-specific microfluidics assay. We highly recommend examining all RNA samples on a denaturing agarose gel or using a fluorometric assay with DNA- and RNA-specific dyes to check samples for DNA contamination. On an agarose gel, gDNA can appear as either a dark mass which remains in the slot if relatively intact or as a high molecular weight smear if it has been sheared during extraction. For RNA-Seq, qRT-PCR, and other sensitive downstream applications, DNase I treatment in solution is recommended. For more information, please contact.

#### Typical Results

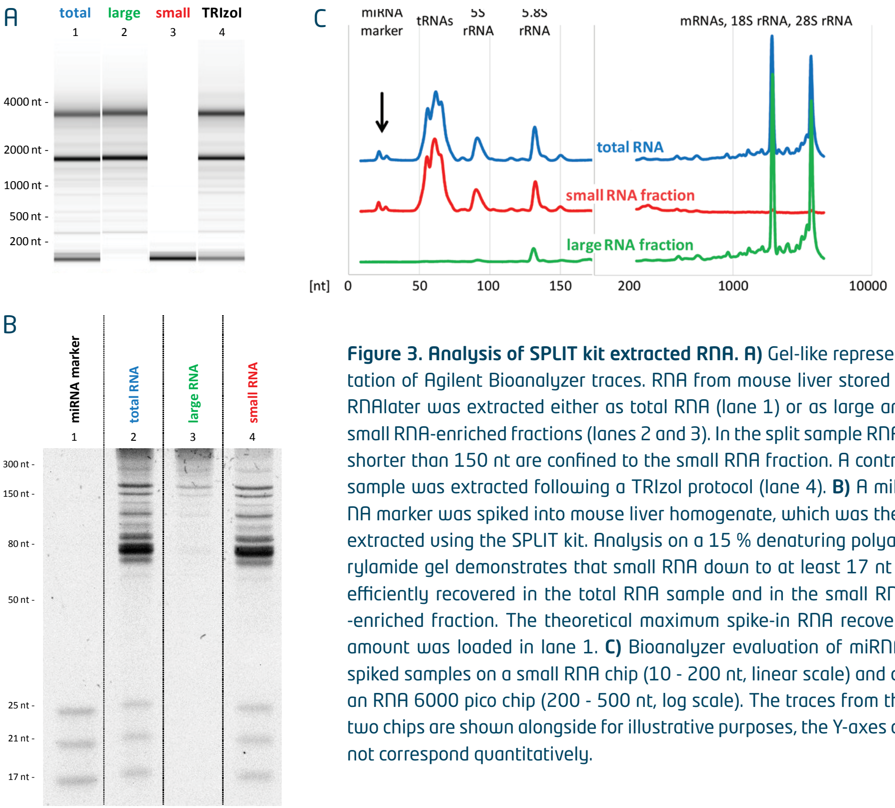

**Figure 3. Analysis of SPLIT kit extracted RNA. A)** Gel-like representation of Agilent Bioanalyzer traces. RNA from mouse liver stored in RNAlater was extracted either as total RNA (lane 1) or as large and small RNA-enriched fractions (lanes 2 and 3). In the split sample RNAs shorter than 150 nt are confined to the small RNA fraction. A control sample was extracted following a TRIzol protocol (lane 4). **B)** A miRNA marker was spiked into mouse liver homogenate, which was then extracted using the SPLIT kit. Analysis on a 15 % denaturing polyacrylamide gel demonstrates that small RNA down to at least 17 nt is efficiently recovered in the total RNA sample and in the small RNA -enriched fraction. The theoretical maximum spike-in RNA recovery amount was loaded in lane 1. **C)** Bioanalyzer evaluation of miRNA-spiked samples on a small RNA chip (10 - 200 nt, linear scale) and on an RNA 6000 pico chip (200 - 500 nt, log scale). The traces from the two chips are shown alongside for illustrative purposes, the Y-axes do not correspond quantitatively.

### 8. Appendix C: RNA Storage

After extraction, RNA can be stored in Elution Buffer (**EB**, 10 mM Tris-HCl pH 7.0) at -20 °C or -80 °C. This minimal buffer stabilizes the pH without any other components that might interfere with downstream applications. When eluting in **EB** we highly recommend the addition of RNase inhibitors to block any accidentally introduced RNases.

The Storage Buffer (**SB**, 10 mM Tris-HCl pH 7.0, 10 mM DTT, and 0.1 mM EDTA) supplied with these kits can be used for intermediate storage of the RNA at -20 °C or -80 °C. DTT (antioxidant) and EDTA (chelating agent) both minimize the threat of RNA degradation, especially at non-freezing conditions. For long-term storage, we recommend keeping aliquots of the RNA as sodium acetate / ethanol precipitate at -80 °C to avoid accidental RNase contamination as well as RNA degradation due to freeze-thaw cycles.

We suggest checking the RNA quality after extended periods of storage for changes in integrity and quantity e.g., on a microfluidics system.

### 9. Appendix D: Extraction of Large and Small RNA-enriched Fractions

RNA can be split into large and small RNA-enriched fractions following the optional column loading protocols below.

**ATTENTION:** The SPLIT RNA Extraction Kit contains reagents for the isolation of total RNA or the large RNA-enriched fraction from 48 samples, or small and large RNA-enriched fractions from 24 samples.

#### 9.1. Column Loading of Large RNA

The large RNA-enriched fraction is precipitated onto a silica column by adding 0.33x volume of isopropanol. The small RNA-enriched fraction will be in the flow-through and can be further purified (see the protocol below).

| | |
|---|---|
| 18 | Determine the volume of the aqueous phase, which may vary, depending on the tissue volume and volume transfer efficiency during homogenization and extraction. Add isopropanol at 0.33x of this volume (e.g., 200 µl isopropanol to 600 µl sample). Mix by vortexing for 10 seconds.<br>**ATTENTION:** For best reproducibility of the size cutoff it is essential to quantify the volume of the aqueous phase exactly. |
| 19 | Place a purification column in a collection tube. |
| 20 | Apply a maximum of 800 µl of the mixture from step 18 (aqueous phase with isopropanol) to the column. |
| 21 | Centrifuge for 20 seconds at 12,000 x g at +18 °C.<br>**ATTENTION:** To isolate the small RNA fraction, pipette the flow-through into a 2 ml tube. |
| 22 | Repeat steps 20 - 21 until the mixture is loaded completely and then proceed to column washing and elution at step 23 in 4.3.2., p.15.<br>**NOTE:** If only the small RNA-enriched fraction is of interest, discard the spin-column containing the large RNA and collect the flow-through into the 2 ml tube. Continue to Column Loading of Small RNA at step 18 in 9.2. (below). Else discard the flow-through. |

#### 9.2. Column Loading of Small RNA

The flow-through obtained in steps 21 - 22 of 9.1. (Column Loading of Large RNA) contains the small RNA fraction and is recovered by precipitation onto a new purification column with the addition of 1x volume of isopropanol.

| | |
|---|---|
| 18 | Determine the total volume of the flow-through in the 2 ml tube (from step 22, 9.1.) and add the same volume of isopropanol. Mix by vortexing for 10 seconds. **EXAMPLE:** 800 µl isopropanol to 800 µl flow-through. |
| 19 | Place a purification column in a collection tube. |
| 20 | Apply a maximum of 800 µl of the mixture from step 18 (aqueous phase with isopropanol) to the column. |
| 21 | Centrifuge for 20 seconds at 12,000 x g at +18 °C and discard the content of the collection tube. |
| 22 | Repeat steps 20 - 21 until the mixture is loaded completely then proceed to column washing and elution at step 23 in 4.3.2., p.15. |

#### 9.3. Short Procedure: Extraction of Large and Small RNA-enriched Fractions

**ATTENTION:** All centrifugation steps are at 12,000 x g and +18 °C!

##### 20 min Homogenization and Phenol / Chloroform Extraction

| | **Homogenization** |
|---|---|
| ☐ | Homogenize sample in 400 µl **IB**. |
| | **Phenol / Chloroform Extraction** |
| ☐ | Centrifuge 1 Phase Lock Gel tube for 1 min. |
| ☐ | Transfer homogenate into a Phase Lock Gel tube. |
| ☐ | Add 400 µl phenol solution pH 4.3, mix by inverting the tube 5 times. |
| ☐ | Add 150 µl **AB**, mix by pipetting. |
| ☐ | Add 200 µl chloroform and mix by repeatedly inverting the tube for 15 sec. |
| | **ATTENTION:** Do not vortex! |
| ☐ | Incubate for 2 min at RT. |
| ☐ | Centrifuge for 2 min. |
| ☐ | Decant the upper phase into a 2 ml tube. |
| | **ATTENTION:** Do not transfer the upper phase by pipetting! |

##### 15 min Purification of Large / Small RNA-enriched Fraction(s)

| | **Column Loading Large RNA-enriched Fraction** |
|---|---|
| ☐ | Measure volume of transferred upper phase. |
| ☐ | Add 0.33x vol. isopropanol to the upper phase. |
| ☐ | Mix by vortexing for 10 sec. |
| ☐☐ | Load max. 800 µl onto purification column in collection tube. Centrifuge for 20 sec. Repeat until mixture is loaded completely. |
| | **ATTENTION:** Keep flow-through and transfer into a 2 ml tube if small RNA extraction is desired. If only small RNA extraction is desired, discard the spin column. |
| | **Column Loading Small RNA-enriched Fraction** |
| ☐ | Measure flow-through volume in 2 ml tube and add 1x vol. isopropanol. Mix by vortexing for 10 sec. |
| ☐☐ | Load max. 800 µl onto new purification column in a collection tube. Centrifuge for 20 sec. Discard flow-through and repeat until mixture is loaded completely. |
| | **Column Washing** |
| ☐☐☐ | Apply 500 µl **WB** to each column and centrifuge for 20 sec. Empty collection tube(s). Repeat this step twice for a total of three washes. |
| ☐ | Centrifuge for 1 min to spin dry column(s). |
| | **Elution** |
| ☐ | Place purification column(s) in a 1.5 ml tube. |
| ☐ | Pre-warm **EB** or **SB** for 5 min at 70 °C. |
| ☐ | Apply 10 - 50 µl of pre-warmed **EB** or **SB**, incubate for 1 min at RT. |
| ☐ | Centrifuge for 1 min. |
| ☐ | **OPTIONAL:** Add RNase inhibitor (not included). |

**NOTE:** Repeat the centrifugation or increase centrifugation time if sample did not pass through the filter completely.

### 10. Appendix E: Revision History

| Publication No. / Revision Date | Change | Page |
|---|---|---|
| 008UG005V0320<br>Oct. 10, 2022 | Legal terms and conditions statements updated. | 2 |
| | Schematic overview updated. | 5 |
| | Kit components figure updated. | 6 |
| | Guidelines replaced with the hyperlink to General Guidelines for Lexogen Kit Use. | 8 |
| | Protocol for Extraction of Large and Small RNA-enriched Fractions moved to Appendix. | 21 - 23 |
| | SPLIT for Blood Cat. No. 099 removed from the User Guide. | 14 |
| 008UG005V0100<br>Aug. 19, 2013 | Initial Release. | |

The RNA Experts

Associated Products:

015 (QuantSeq 3' mRNA-Seq Library Prep Kit for Illumina (FWD))
016 (QuantSeq 3' mRNA-Seq Library Prep Kit for Illumina (REV) with Custom Sequencing Primer)
025, 050, 051, 141 (SIRVs Spike-in RNA Variant Control Mixes)
052 (Small RNA-Seq Library Prep Kit for Illumina)
070 (RS-Globin Block, *homo sapiens*)
095 (CORALL Total RNA-Seq Library Prep Kit)
113 - 115, 129 - 131 (QuantSeq 3' mRNA-Seq Library Prep Kit FWD with UDIs)
117 - 119, 132 - 134 (CORALL Total RNA-Seq Library Prep Kit with UDIs)
128 (TraPR Small RNA Isolation Kit)
144 (RiboCop rRNA Depletion Kit for Human/Mouse/Rat (HMR) V2)
145 (RiboCop rRNA Depletion Kit for Human/Mouse/Rat Plus Globin (HMR+Globin))
171 - 176 (CORALL RNA-Seq V2 Library Prep Kit with UDIs)

### SPLIT RNA Extraction Kit · User Guide

Lexogen GmbH
Campus Vienna Biocenter 5
1030 Vienna, Austria

© Lexogen GmbH, 2022

Lexogen, Inc.
51 Autumn Pond Park
Greenland, NH 03840, USA
www.lexogen.com
